# A liquid-to-solid phase transition of biomolecular condensates drives *in vivo* formation of yeast amyloids and prions

**DOI:** 10.1101/2023.11.17.566816

**Authors:** Maya K. Alieva, Sophia B. Nikishina, Igor I. Kireev, Sergei A. Golyshev, Pyotr A. Tyurin-Kuzmin, Liudmila V. Ivanova, Alexander I. Alexandrov, Vitaly V. Kushnirov, Alexander A. Dergalev

## Abstract

Liquid-liquid phase separation (LLPS) and liquid-solid phase transition (LSPT) of amyloidogenic proteins are now being intensively studied as a potentially widespread mechanism of pathological amyloids formation. However, the possibility and importance of such a mechanism in living systems is still questionable. Here, we investigated the possibility of such LSPT for a series of yeast prion proteins-based constructs overproduced in yeast cells lacking any pre-existing amyloid template. By combining fluorescence and electron microscopy with biochemical and genetic approaches, we have shown that three such constructs (containing the prion domains (PDs) of either Sup35, Rnq1 or Mot3 proteins) form amyloid fibrils via the intermediate stage of liquid-like condensates, that age over time into the more solid-like hydrogels and amyloid bodies. In turn, LSPT of these constructs triggers prion conversion of the corresponding wild-type protein. Two other constructs studied (Ure2- and Sap30-based) are unable to phase separate in vivo and their amyloidogenesis is therefore strongly suppressed. Using PrK-resistant amyloid core mapping, we showed that Sup35PD amyloids formed via LSPT have a different molecular architecture compared to those formed via amyloid cross-seeding. Finally, we showed that physiological LLPS of wild-type Sup35 protein can increase its prion conversion in yeast.

## Introduction

Biomolecular liquid-liquid phase separation (LLPS) is the spontaneous, metastable demixing of protein and/or nucleic acid solutions into highly concentrated and dilute phases. LLPS is usually driven by transient, multivalent interactions between the biomolecules enriched in the concentrated (dense) phase. This dense phase usually represents so-called “liquid droplets”, or “liquid-like condensates” - highly dynamic bodies with physicochemical properties characteristic of liquids: a dynamic shape, propensity to fuse upon contact, and high mobility of the constituent biomolecules - both within and across the boundaries of the condensate.

Discovered more than a century ago as a physicochemical phenomenon in the test tube (as colloids and micelles of biological molecules), LLPS came into the spotlight of the life sciences with the emerging conceptualization of LLPS as one of the most fundamental mechanisms of protein and nucleic acid compartmentalization in living cells (Alberti and Hyman, 2021). LLPS underlies the formation of numerous types of membrane-less organelles (such as the nucleolus, stress granules, P-granules, centrosomes, nuclear speckles, Cajal bodies, and many others (Banani et al., 2017)), membrane-associated protein complexes (such as the PrPC-Aβo complex Kostylev et al., 2018), and contributes to the proper 3D organization of chromatin inside the nucleus (Ulianov et al., 2021).

Many of the proteins that are able to form liquid-like condensates have regions that are intrinsically disordered and/or of low complexity. Remarkably, this feature is also characteristic of the majority of proteins that can aggregate into amyloids - fibrillar homopolymers with a cross-beta structure and the ability to replicate their conformation by self-catalytic sequestration of new monomers at their ends. Unlike liquid-like condensates, amyloids are highly ordered and immobilize the individual protein molecules (called “protomers”) within the fibril. Although the biological functions and effects of amyloids are diverse, many are involved in the pathogenesis of numerous human diseases, including neurodegenerative disorders, type II diabetes, and several others. Amyloids with the apparent ability to infect new cells and organisms in real-life situations are called prions. Prions from multicellular organisms usually manifest themselves as pathogenic agents, whereas yeast prions can be benign (or even beneficial) heritable protein conformers that encode certain epigenetic features (Oamen et al., 2020) - in effect, a proteinaceous form of the gene.

Two well-studied models of amyloid formation include (i) the spontaneous formation of small, ordered pre-fibrillar intermediates in a homogeneous monomeric protein solution, and (ii) the seeding of amyloid polymerization by the catalytic ends of pre-existing amyloid fibrils, which may consist of either the same protein (i.e., homotypic amyloid elongation) or a different protein type (amyloid cross-seeding). In recent years, a growing body of evidence has suggested an additional pathway for amyloid formation: phase-separated, liquid-like protein droplets may be an intermediate for the de novo appearance of amyloid-like polymers. Because of their metastable, highly dynamic nature, liquid-like condensates can evolve and rearrange over time into more solid-like granules - a process termed “Liquid-to-Solid Phase Transition” (LSPT). These aged condensates can be either amyloid-rich particles or so-called “hydrogels” (which, however, can also contain proteinaceous fibrillar structures inside). The formation of cross-beta fibrils in the interior (or on the surface of) the biomolecular condensates has been demonstrated for a number of disease-associated mammalian proteins and peptides (including PrP (Tange et al., 2021), tau (Wegmann et al., 2018), synuclein (Ray et al., 2020), huntingtin (Peskett et al., 2018), FUS (Patel et al., 2015; Murakami et al., 2015), TDP43 (Babinchak et al., 2019), hnRNPA1 (Molliex et al., 2015), IAPP (Pytowski et al., 2020), as well as for D. melanogaster functional amyloid-forming protein Orb2 (Ashami et al., 2021), the SARS-CoV2 nucleoprotein (Tayeb-Fligelman et al., 2021), and molecularly engineered silk proteins (Mohammadi et al., 2018). These examples include the formation of both “classical” irreversible amyloids (resistant to detergents, chaotropic agents, and mild heating) and “soft”, metastable amyloid-like fibrils (Murakami et al., 2015). However, most of these studies have investigated the LSPT of biomolecular condensates in vitro, in designed buffer conditions that are often far from physiological. Inside the living cells, whose interiors are densely crowded with thousand types of different biological macromolecules, the formation and evolution of liquid-like condensates may follow different trajectories governed by the different physicochemical environments. Therefore, the creation and validation of a convenient cellular model for studying the liquid-to-solid phase transition of protein-rich liquid droplets could help to move the ball forward in this field. In addition, although the mammalian prion protein PrP has already been shown to form amyloid fibrils via LLPS and LSPT (Tange et al., 2021), to our knowledge, it remains unclear whether the amyloids of such origin are capable of prionic (i.e., infectious or heritable) behavior. The structural characteristics of amyloids formed by the aging of liquid condensates are also generally uninvestigated.

In this work, we have studied in vivo LLPS and LSPT on a series of GFP-tagged chimeric constructs containing the prion domains of several yeast prionogenic proteins - Sup35, Rnq1, Ure2, Mot3 and Sap30 - overproduced in the yeast cells lacking a pre-existing amyloid template. Using a range of microscopic and biochemical techniques, we found that three of them - the constructs based on Sup35, Rnq1 and Mot3 prion-like domains - form liquid-like condensates, that further rearrange into more solid-like hydrogels and/or amyloid-rich aggregates. Ure2- and Sap30-based constructs did not form macroscopic condensates and showed either lacking or dramatically declined amyloidogenesis under the same conditions. Therefore, liquid-like condensates appear to be an essential intermediate for amyloidogenesis by such constructs in situations where the pre-existing amyloid template is not available. Ageing of Sup35- and Mot3-based constructs condensates significantly increased the formation of the prion form of Sup35 and Mot3 proteins, respectively. At least, for the Sup35 protein this mechanism of prion formation appears to be physiologically relevant, since wild-type, non-overproduced Sup35 protein increasingly switches to the prion form after prolonged incubation of the cells under conditions favoring the stress-inducible LLPS of this protein. Finally, we showed that amyloids formed by the Sup35-based construct via two different pathways - either LSPT of biomolecular condensates, or amyloid cross-seeding with another yeast prion - differ in both structural and physicochemical properties.

## Results

### Overproduced Sup35PD-GFP forms large liquid-like condensates in a concentration-dependent, stress-independent manner

Sup35 is a gold standard yeast prion protein, the heritable amyloid form of which is known as the [PSI+] prion. Sup35 has a large C-terminal globular domain (e.g. 254-685), which catalyses translation termination, and a disordered N-terminal region (e.g. 1-253), which has traditionally been divided into the Q/N-rich N domain and the highly charged central (M) domain. The N domain is usually referred to as the ‘prion domain’ - however, we and others have recently shown that the proximal part of the M domain (at least up to a.a. 153) contains segments involved in the rigid amyloid core for some of the prion variant-specific conformations (Dergalev et al., 2019; Ohhashi et al., 2018; Frederick et al., 2014). We therefore refer to the entire intrinsically disordered NM region as the “prion domain” (Sup35PD).

Sup35PD forms large linear or circular amyloid aggregates (so-called ‘rings’) when overproduced in yeast carrying the [RNQ+] prion, which acts as an imperfect physical template for amyloid cross-seeding (Zhou, 2001; Tyedmers, 2010; Saibil, 2012). Recent studies have shown that both full-length Sup35 and Sup35PD can also undergo liquid-liquid phase separation in cells lacking a pre-existing amyloid template, both in vitro and in vivo at different production levels (Franzmann et al., 2018; Khan et al., 2018). This process is triggered by cytosolic acidification, with the negatively charged distal part of the Sup35 M domain acting as a pH sensor (Franzmann et al., 2018). In this study, we aimed to investigate the metamorphosis of Sup35PD condensates in vivo during long-term incubation of the cells (as a stationary yeast culture). In yeast culture, the switch from logarithmic to stationary growth phase leads to acidification of both the culture medium and the cell cytosol, which decreases cell viability over the course of long-term incubation (Murakami, 2012). Therefore, in order to observe protein aggregation in viable cells, we decided to use a culture medium buffered at a less toxic pH and aimed to reduce the dependence of Sup35PD phase separation on the acidity of the solution. To this end, we created a Sup35PD allele with a deletion of the negatively charged cluster consisting of 14 a.a. (Δ240-253) in the distal part of the M domain (Figure 1A). This construct was C-terminally tagged with eGFP (henceforth, the resulting protein will be referred to as Sup35PD-GFP) and inserted into a multicopy plasmid under the control of a strong GAL1 promoter, resulting in approximately 200-fold overproduction relative to endogenous Sup35 levels (Figure S1A). Since Sup35 phase separation has been shown to be highly dependent on its N domain (Franzmann et al., 2018), and Sup35 amyloidogenicity is dependent on the region 1-153 a.a. (Dergalev et al., 2019), we created the derivative construct with the deletion of 152 amino acids from the Sup35 N-terminus (hereafter - Sup35(ΔPD)-GFP) to use as a negative control mutant that is incapable of both forms of assembly.

**Figure 1.**
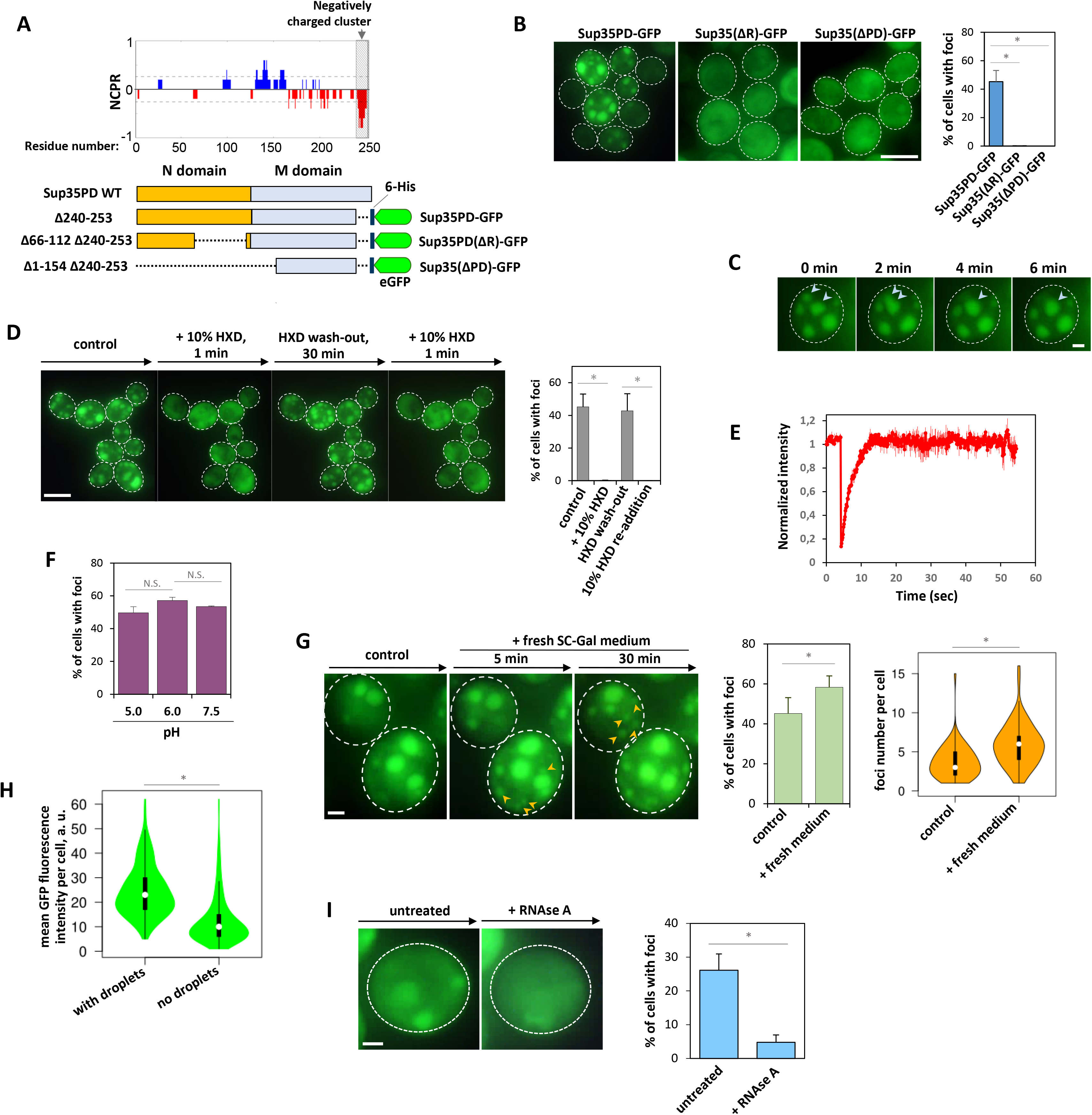
Sup35PD-GFP forms liquid-like condensates when overproduced in cells lacking a pre-existing amyloid template. **A.** The scheme of Sup35-based constructs used in the work. Sup35PD-GFP was used as a construct capable for both LLPS and amyloidogenesis, but with deletion of the proximal, negatively charged “pH sensor” cluster (a.a. 240-253); Sup35PD(ΔR)-GFP was used as a mutant that able to form amyloids and prions, but deficient for LLPS; and Sup35(ΔPD)-GFP was used as a mutant deficient for both amyloidogenesis and LLPS. The diagram of Net Charge Per Residue (NCPR) of Sup35NM region was created with CIDER algorythm. **B.** {Left} Representative maximum projected images of Δrnq1 cells overproducing either Sup35PD-GFP, Sup35(ΔR)-GFP or Sup35(ΔPD)-GFP construct for 24 hrs. The scale bar is 5 μM. {Right} Quantification of the percentage of cells carrying the assembled form of the indicated protein. Data are mean of 3 biological replicates, error bars are SEM. * p-value < 0.001 (two-tailed Student’s t-test). **C.** Fusion of two Sup35PD-GFP assemblies (blue arrowheads) upon contact. Scale bar is 1μM. **D.** Sup35PD-GFP assemblies can be rapidly and reversibly disrupted by 10% 1,6-hexanediol (HXD). {Left} Representative maximum projected images of digitonin-treated, concanavanine A-immobilised Δrnq1 cells overproducing Sup35PD-GFP for ∼18h. Cells were sequentially treated with 82 mM phosphate-citrate buffer (pH 6.5) with and without 10% HXD (as indicated). Scale bar is 5 μM. {Right} Quantification of cells with Sup35PD-GFP foci. The experiment was performed in 3 biological replicates, with 200 to 300 cells counted per replicate. Data are expressed as mean + SEM. * p-value < 0.01 (two-tailed Student’s t-test). **E.** Half-FRAP analysis of the Sup35PD-GFP assemblies formed after its 20-hrs long overproduction. Mean values (± SEM) are shown. **F.** Proportion of cells bearing condensates in suspensions of cells with Sup35PD-GFP overproduction, incubated in Na-citrate buffer with different pH values. To equate pH values inside and outside of the cells, 2 mM 2,4-dinitrophenol were added. Data are expressed as mean + SEM (n = 3). N. S. – non-significant (two-tailed Student’s t-test). **G.** Sup35PD-GFP condensates are not disassembled when cells recover from the stationary phase. {Left} Exemplary maximum projected images of concanavalin A-immobilised cells with Sup35PD-GFP overproduction over ∼18h. Cells were treated with fresh SC-Gal medium (pH 6.5). Sup35PD-GFP foci that appeared after the addition of fresh medium are indicated by the orange arrows. The scale bar is 1 μM. {Centre} Quantification of cells with Sup35PD-GFP foci (old medium compared with 30 min in fresh medium). The experiment was performed in 3 independent biological replicates, with 200 to 300 cells counted per replicate. Data are expressed as mean + SEM. {Right}: Normalized distribution of the number of Sup35PD-GFP foci per cell in populations of cells either treated for 30 min with fresh SC-Gal medium (n = 71 cells) or left in the old culture medium (n = 68 cells). * p-value < 0.01 (two-tailed Student’s t-test). **H.** Violin plots show the distribution of GFP signal level in the cells with and without Sup35PD-GFP condensates. White circles show the medians; box limits indicate the 25th and 75th percentiles as determined by R software; whiskers extend 1.5 times the interquartile range from the 25th and 75th percentiles; polygons represent density estimates of data and extend to extreme values. * p-value < 0.01 (two-tailed Student’s t-test). **I.** Sup35PD-GFP assemblies can be disrupted by the cell treatment with RNAse A (1 mg/ml) solution. {Left} Representative maximum projected images of concanavanine A-immobilised Δrnq1 cells overproducing Sup35PD-GFP for ∼18h. Cells were sequentially treated with 82 mM phosphate-citrate buffer (pH 6.5) with and without RNAse A (as indicated). Scale bar is 1 μM. {Right} Quantification of cells with Sup35PD-GFP foci. The experiment was performed in 3 biological replicates, with 200 to 300 cells counted per replicate. Data are expressed as mean + SEM. * p-value < 0.01 (two-tailed Student’s t-test).

To obtain Sup35PD-GFP condensation (instead of amyloid cross-seeding), we overproduced this protein in yeast cells lacking [RNQ+] prion. Since [RNQ+] prion can appear spontaneously in [rnq-] cells (Derkatch, 2000), we used a strain with a deletion of the gene encoding the Rnq1 protein (Δrnq1). After approximately 18 hrs of overproduction of Sup35PD-GFP in the stationary yeast culture, we observed relatively large (µm- or sub-µm-scale), quasi-spherical aggregates that appeared in ∼45% of the cells with Sup35PD-GFP signal. As a control, Sup35(ΔPD)-GFP did not form any visible aggregates, confirming that these aggregates are predominantly formed by interactions of the intrinsically disordered Sup35PD region, rather than potential oligomerisation of eGFP (Figure 1B). The aggregates were numerous and moved slowly in the cytosol with occasional coalescence (Figure 1C), followed by subsequent restoration of the quasi-spherical shape of the resulting aggregate, suggesting a liquid-like physical state. The shape of such assemblies was generally dynamic (Video S1), and in the budding cells such assemblies were sometimes observed squeezing through the narrow bud neck into the daughter cell (Figure S1B), which together also supports their liquid nature. The probing of these assemblies with 1,6-hexanediol (HXD) and FRAP analysis - two standard methods used to detect the material properties of protein assemblies (Kroschwald, 2017) - confirmed this suggestion: they were rapidly and reversibly dissolved by 10% HXD solution (Figure 1D), and photobleached Sup35PD-GFP molecules were promptly exchanged with unbleached ones from the surrounding cytosol (t1/2= 2.65 s, immobilised fraction = 0%, Figure 1E). As a negative control, the well-known Sup35PD-GFP amyloid ‘rings’ formed in the [RNQ+] strain (Tyedmers, 2010; Saibil, 2012) were not affected by HXD even after 30 min treatment (Figure S1F). Therefore, we concluded that the observed assemblies are liquid-like proteinaceous droplets, similar to those previously observed in cells with hyperproduced Sup35NM-mEos3.1 construct (Khan et al., 2019).

To determine whether Sup35PD-GFP droplets represent one of the physiological cytosolic yeast condensate types, we examined the colocalisation of condensed Sup35PD-GFP with the markers of either stress granules or P-bodies (Pab1 and Dcp2, respectively) fused to tagRFP. Pab1-RFP was enriched in Sup35PD-GFP condensates, whereas Dcp2-RFP formed separate aggregates and showed no colocalisation with Sup35PD-GFP (Figure S1C). However, in cells with overproduction of Sup35(ΔPD)-GFP (non-aggregating Sup35PD mutant), Pab1-RFP was either diffuse (75% of cells with GFP signal) or formed 1-2 dots per cell (which can be attributed to the “classic” stress granules), whereas in cells with Sup35PD-GFP condensates we observed more Pab1-RFP dots, some of which colocalised with Sup35PD-GFP condensates (Figure S1D). Thus, the joint formation of Sup35PD-GFP/Pab1-RFP foci is likely initiated by the homotypic and heterotypic interactions of Sup35PD-GFP molecules (with subsequent recruitment of Pab1-RFP), rather than by Sup35PD-GFP recruitment into the true stress granules. We also found that fresh Sup35PD-GFP condensates disassembled where the cells had been treated with RNAse A solution (Figure 1I), suggesting that RNA molecules are likely to be involved in these assemblies and even play a scaffolding role in their formation and maintenance.

Stationary phase yeasts are usually considered to be chronically stressed by nutrient depletion, cytosolic acidification and accumulation of toxic metabolites (Gray et al., 2004). Energy depletion has been shown to promote Sup35 phase separation at nearly endogenous levels (Franzmann et al., 2018). In the later work, Sup35-GFP foci were readily disassembled when energy depletion was relieved by the addition of fresh culture medium. Therefore, we asked whether energy depletion of cells in stationary phase culture was a key factor for Sup35PD-GFP condensation in our experimental setup. When cells with pre-formed Sup35PD-GFP droplets were exposed to fresh SC-Gal medium, none of the droplets disappeared even after an hour of incubation - instead, a number of additional fluorescent puncta appeared in many cells (Figure 1G). Furthermore, direct manipulation of the pH of the yeast cytosol (by treating the cells with pH-buffered solutions containing 2mM dinitrophenol as a cell membrane permeabilizer for protons) showed that Sup35PD-GFP droplet formation is almost unaffected by this parameter, at least in the pH range 5.0-7.5 (Figure 1F). Finally, overproduced Sup35PD-GFP also formed liquid-like condensates in the logarithmic phase yeast cells, with almost the same frequency as in the stationary phase yeasts (Figure S1E).

LLPS in protein solutions is generally considered to be a concentration-dependent process that occurs when the saturation concentration (C_sat_) is exceeded. To investigate the concentration dependence of Sup35PD-GFP condensation in yeast cells, we measured the mean GFP fluorescence intensity (as a proxy for Sup35PD-GFP concentration) in the images of individual cells with and without Sup35PD-GFP condensates. As expected, the subpopulation of droplet-positive cells has a mean Sup35PD-GFP level approximately 2-fold higher than those without droplets (Figure 1H). However, the distributions of Sup35PD-GFP levels in the droplet-positive and droplet-negative cell subpopulations overlap significantly, suggesting that the intracellular C_sat_ for Sup35PD-GFP is not fixed in yeast. Instead, it may be influenced by the physicochemical parameters of the cytosol in individual yeast cells and by interactions with other proteins.

### Sup35PD-GFP liquid droplets age into amyloid-containing hydrogels and amyloid ring-like aggregates upon long-term incubation of cells

Franzmann and colleagues have shown that Sup35 liquid droplets formed in vitro are prone to rapid gelification, becoming solid particles with a sponge-like underlying structure in nearly 60 minutes after formation (Franzmann et al., 2018). In our in vivo experimental system, the numerous Sup35PD-GFP droplets began to form 6-8 hours after the addition of galactose - and 10-16 hours later they still behaved as liquid-like bodies. We then asked whether prolonged incubation (at longer timescales) of the cells with the Sup35PD-GFP liquid droplets would lead to their gelification and possibly to massive formation of Sup35PD-GFP amyloids. To address this issue, we incubated the stationary yeast culture for up to 10 days in SC-GAL medium buffered at pH 6.5 to prevent acidification-induced cell death (Murakami, 2012).

First, we took time-lapse samples of yeast cultures, subjected them to live-cell fluorescence microscopy and tracked the dynamics of the number, size and morphology of Sup35PD-GFP assemblies. The droplets, which initially appear in the cell as numerous small foci, then tend to coalesce into one or several larger condensates per cell (Figure S2A). Over the course of cell incubation, these condensates gradually lost their sphericity and became more irregular in shape (Figure 2A, 2B, 2C). In contrast to fresh droplets with a relatively smooth surface, the surface of aged condensates becomes more ‘knobby’ (Figure 2B). In many cells, aged condensates tend to localise close to the cell surface and become flatter, probably due to squeezing between the cell membrane and the enlarged vacuole (the formation of such large vacuoles is a common phenotypic feature of starved yeasts). In contrast to fresh droplets, which are dynamic in shape and position in living cells, ageing condensates become more rigid and have an “arrested” shape (Video S1). Such changes in condensate morphology and dynamics could reflect the gelification of initially liquid-like aggregates, as in the liquid-solid phase transition of Sup35PD-GFP condensates. To test this hypothesis, we applied a number of techniques that address different physicochemical properties of proteinaceous particles. First, we measured the dynamics of condensate dissolution by 10% HXD at different stages of cell incubation. As the “chronological age” of the condensates increased, their dissolution slowed down (Figure 2J), which could indicate the formation of stronger intermolecular bonds within the condensates. Furthermore, in contrast to the ‘fresh’ condensates, the older condensates were characterised by the presence of an HXD-insoluble Sup35PD-GFP fraction, which appears to reach ∼50% in 7-day-old condensates. Furthermore, FRAP analysis revealed a steady decrease in the mobility of the molecules within the aged condensates (Figure 2I). Using this method, we also detected a compatible proportion of the immobile Sup35PD-GFP fraction accumulated within the condensates (at the respective time points) compared to the HXD-insoluble fraction in the time-lapse HXD treatment experiment. In addition, we found that from the second day of incubation in SC-Gal medium, Sup35PD-GFP condensates lose their sensitivity to RNAse A treatment (Figure S2D), suggesting that emerging strong protein-protein interactions may replace the proposed scaffolding role of RNA molecules in maintaining condensate integrity.

**Figure 2.**
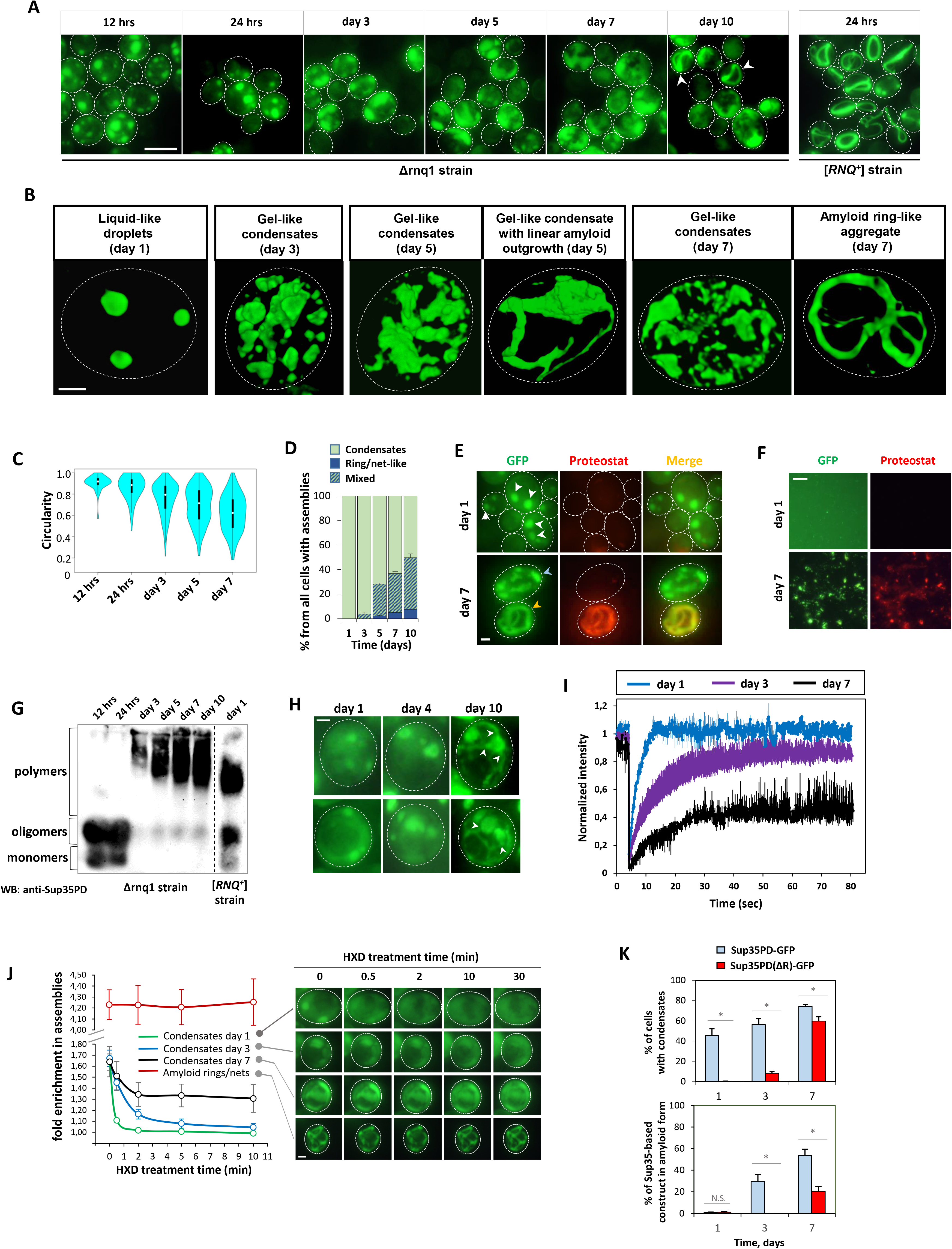
Sup35PD-GFP liquid condensates can age into the amyloid-rich aggregates. **A.** Time-lapse maximum intensity projections of Δrnq1 cells with overproduction of Sup35PD-GFP protein. White arrowheads indicate linear aggregates similar with “amyloid rings” formed by the same protein in [RNQ+] cells (which are shown on the right panel for comparison). Scale bar is 5 μM. **B.** Volume snapshots of 3D-rendered SIM images of fresh and aged Sup35PD-GFP condensates: liquid-like droplets, solidified gel-like condensates, and amyloid ring-like aggregates. Scale bar is 1 μM. **C.** Sup35PD-GFP condensates became less circular over time. The time-resolved distribution of circularity values of different individual Sup35PD-GFP condensates is shown with violin plots. Δrnq1 cells with overproduction of Sup35PD-GFP were incubated as a stationary culture over 7 days, z-stacked images were obtained at indicated time points, and circularity values of 200-300 individual Sup35PD-GFP assemblies (per single time point) were calculated with FIJI software. White circles show the medians; box limits indicate the 25th and 75th percentiles as determined by R software; whiskers extend 1.5 times the interquartile range from the 25th and 75th percentiles; polygons represent density estimates of data and extend to extreme values. **D.** Time-lapse proportion between cells with different categories of Sup35PD-GFP assemblies: liquid-/gel-like condensates, linear/circular amyloid aggregates and mixed type (typically, gel condensates with linear outgrowth). Experiment was performed in 3 biological replicates, error bars represent SEM. **E.** Staining of Sup35PD-GFP assemblies, formed in Δrnq1 cells via LLPS and LSPT, with amyloid-specific fluorescent dye proteostat. In contrast to freshly-formed droplets (white arrowheads) and aged hydrogels (blue arrowhead), the linear/circular aggregates (orange arrowhead) stain with proteostat. Scale bar is 1 μM **F.** Proteostat-stained lysates of Δrnq1 cells with Sup35PD-GFP overproduced for either 1 or 7 days. Cells were lysed by shaking with the glass beads. Scale bar is 5 μM. **G.** Time-lapse SDD-AGE analysis of the dynamics of amyloid polymers formation in Δrnq1 cells with overproduction of Sup35PD-GFP. Cell lysates were normalized to the amount of Sup35PD-GFP. As a positive control for amyloid mobility, we used lysates from [RNQ+] cells overproducing Sup35PD-GFP for 1 day. The experiment was performed in 3 biological replicates, the representative image is shown, and the quantification of amyloid formation rate is presented in Figure 6I. **H.** Time-lapse fluorescence images of the single cells overexpressing Sup35PD-GFP. After 24 h incubation in SC-Gal medium, cells were spotted on agarose pad (made on the citrate-phosphate buffer, pH = 6.5) and single cells were imaged at the indicated time points. White arrowheads indicate linear protrusions emerging from the initial condensate. Scale bar is 1 μM. **I.** Time-resolved FRAP analysis of the ageing Sup35PD-GFP condensates (1^st^, 3^rd^ and 7^th^ day of Sup35PD-GFP overproduction). Mean values (± SEM) are shown (2-5 replicates per each incubation time). **J.** {Left} Quantification of Sup35PD-GFP time-lapse fold enrichment within ageing condensates and ring-like aggregates (calculated as the ratio of GFP fluorescence intensity inside/outside particles) in cells under 10% HXD treatment. Cells overproducing Sup35PD-GFP for different time periods (1, 3 and 7 days) were immobilised with concanavalin A, exposed to 10% HXD solution and imaged at 0, 0.5, 2, 5 and 10 min after HXD addition. The time-resolved inside/outside fluorescence intensity ratio (IR) was calculated for 8-10 individual assemblies of each type, the data are expressed as mean ± SEM. {Right} Representative images of HXD-treated cells. Scale bar is 1 μM. **K.** Amyloid switch of Sup35-based constructs in Δrnq1 cells correlates with their ability to phase separate. (UP) Comparison of condensate-forming ability of Sup35PD-GFP and Sup35PD(ΔR)-GFP constructs overproduced in Δrnq1 strain. Proportion of cells with assembled form of respective protein (from all GFP-positive cells) was calculated for 4 independent biological replicates at different time points ranging from 1 to 7 days of incubation in SC-GAL medium. (DOWN) Time-lapse proportion of Sup35-based construct in amyloid (SDS-insoluble) form determined by SDS-PAGE with boiling followed by Western Blotting. At indicated time points, the lysates of Δrnq1 cells with overproduction of either Sup35PD-GFP or Sup35PD(ΔR)-GFP were subjected to SDS-PAGE followed by Western Blotting with immunostaining against GFP. Soluble form was detected as a protein band that enter the gel initially, the amyloid form – as a band that enter the gel just after boiling (see Material and Methods). The intensities of the protein bands were quantified with FIJI software, and the data are expressed as mean + SEM, based on four independent biological replicates. * p < 0,05, N.S. – non-significant (two-tailed Student’s t-test). The blot images are presented in the Figure S2F.

We also noticed that in addition to ageing condensates, another type of Sup35PD-GFP aggregates starts to appear in long-term incubated yeast cells. This type of assembly looks similar to the well-known Sup35PD amyloid ‘rings’ - ring- or ribbon-like aggregates that have been shown to be large and highly ordered bundles of parallel amyloid filaments (Kawai-Noma, 2010; Saibil, 2012). In our setup, these ring- or ribbon-like aggregates typically started to appear from day 3 of Sup35PD-GFP overproduction. Their proportion gradually increased until the final time point of the experiment (day 10), reaching 7.6 ± 0.15% of aggregate-bearing cells (Figure 2D). From day 3, we also observed some cells with a mixed pattern, where the aged condensates had one or more linear protrusions (Figure 2B, 2H). The proportion of cells with this pattern also increased significantly over the course of cell incubation (Figure 2D). Based on their morphology, we hypothesised that such linear aggregates consist exclusively of long amyloid fibrils. Staining of cells with proteostat, an amyloid-specific red fluorescent dye, supported this hypothesis (Figure 2E). In addition, time-lapse HXD treatment showed that Sup35PD-GFP molecules in such aggregate type are linked by strong protein-protein interactions, as these aggregates did not fuse when cells were treated with this chemical (Figure 2J).

Since the interior of the cell is a densely crowded environment in which the rate of molecular diffusion can be influenced by many factors, we tried to confirm the changes in the physical state of liquid condensates in an alternative way, by placing such aggregates in different physicochemical environments. To this end, we spheroplasted cells with either fresh or aged condensates and then filmed the process of cell bursting due to hypoosmotic-induced water uptake. As expected, the fresh Sup35PD-GFP droplets quickly disappeared within seconds after the loss of cell integrity, whereas the aged condensates were able to persist as integral fluorescent particles outside the cells - although they also lost part of their fluorescent signal (Video S2). Curiously, we were unable to observe and film the process of exploding cells with a ring-like Sup35PD-GFP aggregation pattern, although such solid-like aggregates were also found in large numbers outside the cells, and their linear size (in the extracellular space) typically exceeded the size of a typical yeast cell (Figure S2H). This led us to believe that cells with this type of aggregation could rupture almost immediately after digestion of the cell wall - probably due to the mechanical pressure that over-packed amyloid bundles could potentially exert on the cell membrane from the inside.

To elucidate the dynamics of Sup35PD-GFP amyloid accumulation in yeast culture with Sup35PD-GFP condensates, we subjected the lysates of yeast incubated for different time periods to SDS-AGE. The large SDS-resistant polymers started to appear from the 3rd day of incubation and their load gradually increased until the end of the time-lapse experiment (Figure 2G). As a control to estimate the dependence of amyloid aggregation on the initial phase separation, we used a Sup35PD-GFP-based construct with deletion of the region from the 3rd to the 5th oligonucleotide repeat, designated Sup35PD(ΔR)-GFP (Figure 1A). Such a construct under the control of the GAL1 promoter has an average production level approximately two times higher than a Sup35PD-GFP construct (Figure S1A). In [RNQ+] cells, it forms amyloid aggregates at the level compatible with Sup35PD-GFP (Figure S2E). In Δrnq1 cells, overproduced Sup35PD(ΔR)-GFP was initially (day 1) barely able to phase separate (Figure 1B, 2K). However, it formed condensates in approximately 8% of cells incubated for 3 days (Figure 2K). In contrast, Sup35PD-GFP formed condensates in more than 40% of the cells from the first day of incubation. Using the “boiled gel” assay, we found that Sup35PD-GFP formed a significant proportion of the amyloid form by day 3, whereas Sup35PD(ΔR)-GFP was converted to the amyloid form with a significant delay (a significant amount of Sup35PD(ΔR)-GFP amyloid was detected at day 7) and in lower proportions (Figure 2K). Thus, for both constructs, LLPS initially precedes amyloid aggregation and the intensity of amyloidogenesis correlates with the intensity of phase separation.

To examine the ultrastructure of aged Sup35PD-GFP condensates, we took advantage of Correlative Light and Electron Microscopy (CLEM). As can be seen from the electron tomography slice, the hydrogel area is more electron dense than the cytosol outside (Figure 3). Inside the hydrogel we can see amyloid fibrils, both single and in bundles, protruding in different directions. Thus, the Sup35PD-GFP hydrogels are indeed the amyloid-containing bodies - but not all the Sup35PD-GFP inside the hydrogels is in the amyloid form.

**Figure 3.**
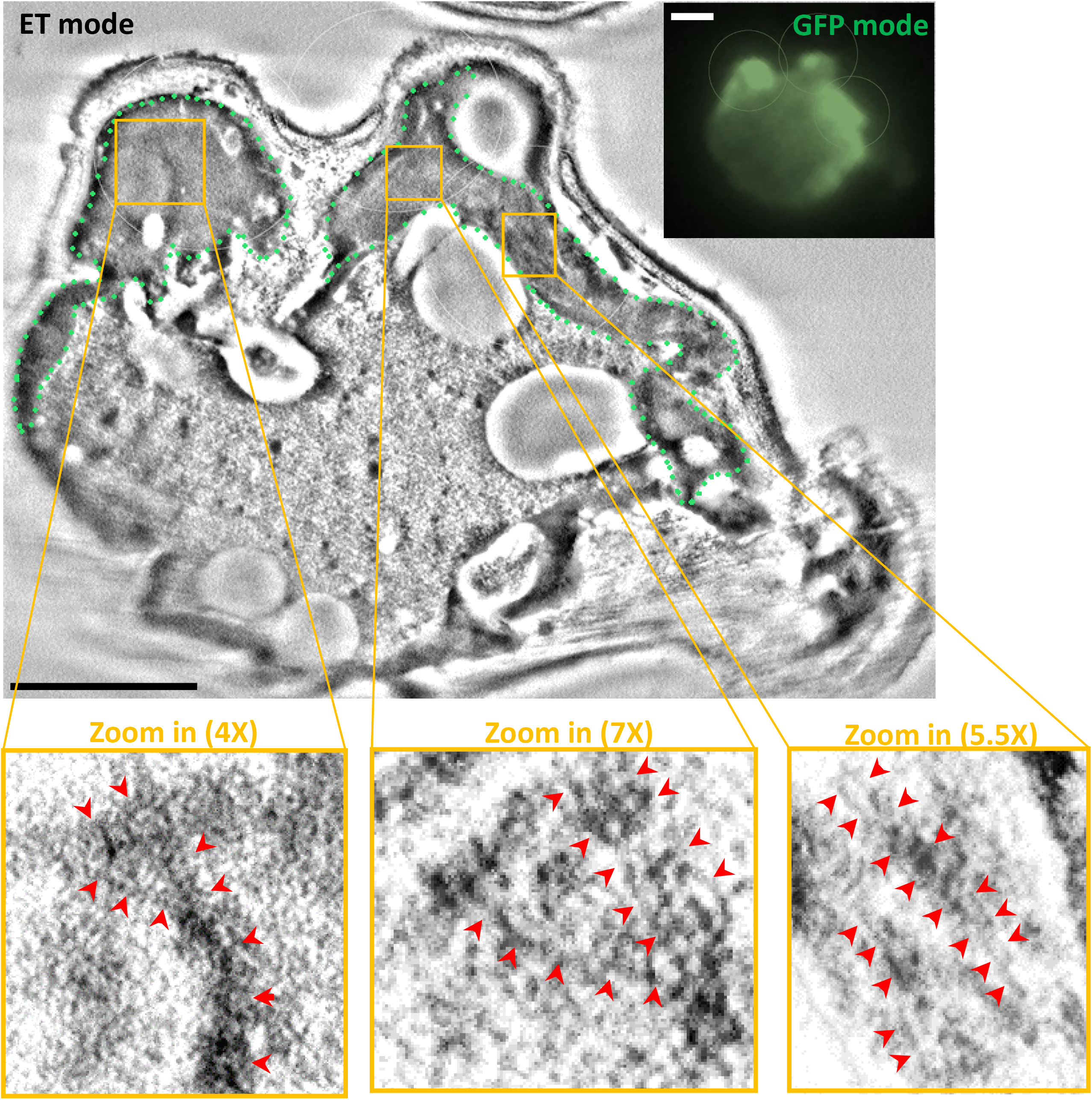
Ultrastructure of Sup35PD-GFP hydrogel (10 day cell) as found by correlative light and electron microscopy. Transmission electron tomography (ET) slice is shown on the left, fluorescence microscopy image is shown on the right. The hydrogel area on ET image is highlighted with the green dots, the amyloid filaments are highlighted with the red arrowheads. Scale bar is 1μM.

### Condensation and liquid-to-solid phase transition of Sup35PD-GFP construct drives the prion switch of endogenous Sup35 protein

Inside yeast cells, Sup35 amyloids can exist as a cytosolic population of numerous and relatively short fibrils that are stably transferred to daughter cells during cell division. Such highly heritable Sup35 fibrils represent a [PSI+] prion and tend to confer a number of epigenetic traits, mainly related to translation termination efficiency (Kochneva-Pervukhova et al., 2001). Another type of Sup35 fibrils, which form large amyloid rings in the [RNQ+] strain, are longer (Saibil et al., 2012) and have lower heritability (Tyedmers et al, 2010) - but can still rearrange over time into a more fragmented and heritable form, giving rise to the [PSI+] prion (Mathur et al., 2010).Since Sup35 condensates undervent LSPT to form amyloid-containing particles, we asked whether the [PSI+] prion state could be established in some progeny cells. A rough estimation of amyloid fibril size from their mobility on SDD-AGE showed that Sup35PD-GFP amyloids formed by LSPT are initially (3rd day of overproduction) much larger than typical [PSI+] prion polymers, but later (7th day) their size was partially shifted towards the smaller values characteristic of [PSI+] prion fibrils (Figure 4A). Based on this observation, we expected that LSPT-derived Sup35PD-GFP amyloids could behave as prions themselves, and possibly confer such prion behaviour on wild-type endogenous Sup35, which was previously shown to co-aggregate with overproduced Sup35PD in its amyloid form (Sharma et al., 2017).

**Figure 4.**
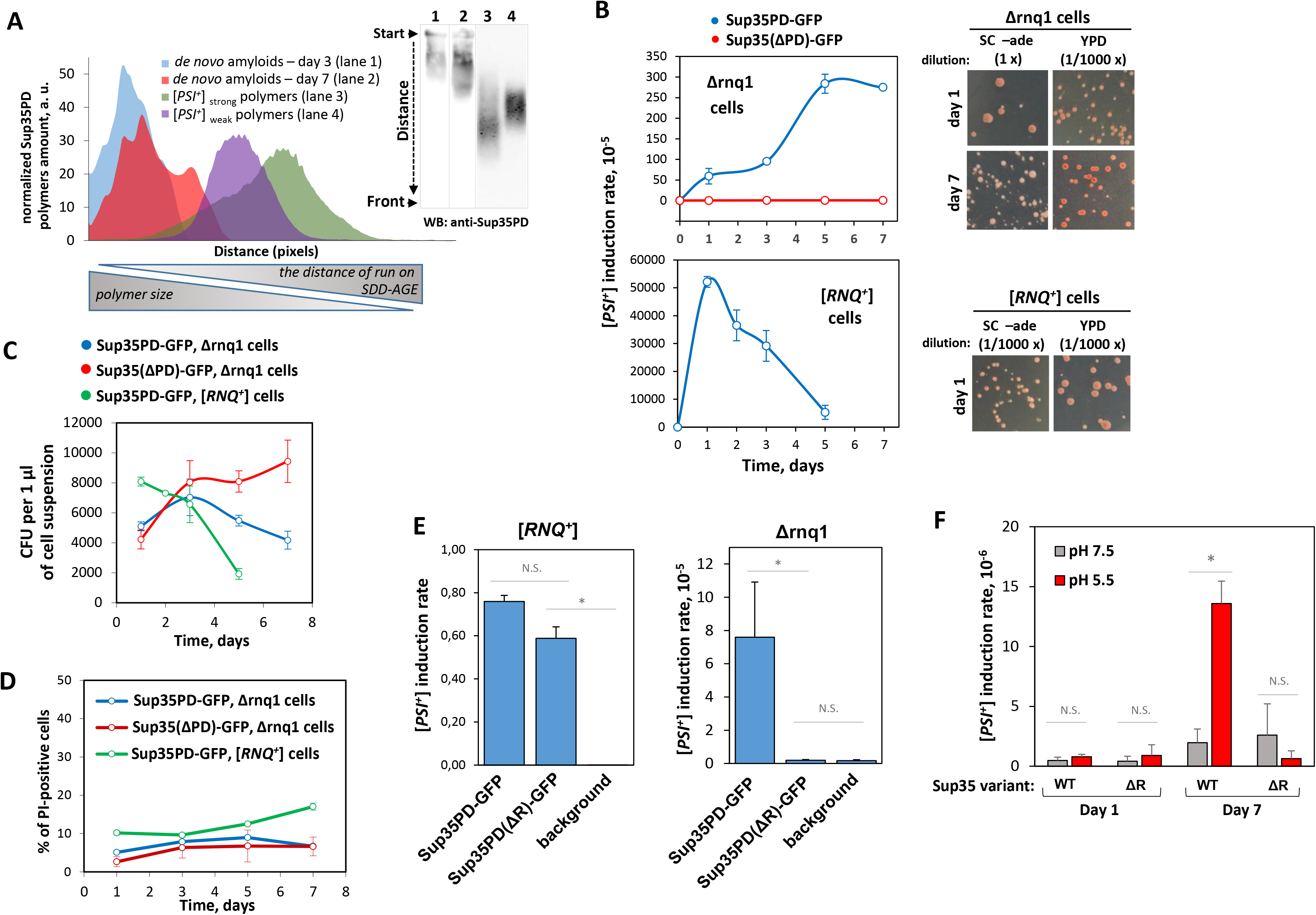
Sup35PD-GFP LSPT can drive de novo formation of [PSI+] prions. **A.** The size distribution of Sup35PD-GFP amyloid polymers of different origin (either [*PSI+*] prion polymers, or *de novo* formed in cells with aged Sup35PD-GFP condensates), estimated via their mobility on 1,8% agarose gel electrophoresis. SDD-AGE analysis of the lysates of [*psi-*] Δrnq1 cells (at 3rd and 7th day of Sup35PD-GFP overproduction), and [*PSI+*] cells (strong and weak prion variants, w/o Sup35PD-GFP overproduction) was performed, and the distribution of the Sup35PD-GFP signal along the lane was quantified using FIJI software. Long-term incubation of yeast with Sup35PD-GFP condensates leads to a shift of the size of the emerging Sup35PD-GFP amyloid polymers into the area corresponding to that of the established [*PSI+*] prion polymers. **B.** Time-lapse measurement of [PSI+] prion formation in cells overproducing Sup35PD-GFP, where this protein is involved in either LLPS (Δrnq1 cells) or heterologous amyloid cross-seeding by Rnq1 polymers ([RNQ+] cells). Sup35(ΔPD)-GFP, a non-polymerising Sup35 mutant, was overproduced in Δrnq1 cells as a negative control to assess the background level of [PSI+] prion formation. The stationary cell cultures were incubated for up to 7 days and plated on either SC-Glu medium lacking adenine or adenine-rich YPD medium at the indicated time points. The plated cells were then grown at 30°C for either 3 days (YDP) or 10 days (SC-Glu w/o adenine) and the number of CFU was counted. Ade+ colonies were confirmed to be [PSI+] prions by curing their nonsense-suppression phenotype on YPD plates containing 4 mM GuHCl (Figure S4A). The mean percentage of [PSI+] clones was calculated for 4-5 biological replicates. Error bars show SEM. **C.** Time-lapse dynamics of the viable cell density, measured as the number of colony-forming units (CFU) yielded from 1 μl of cell suspension streaked on YPD plates. The mean CFU/μl values were calculated for 4-5 independent biological replicates. Error bars show SEM. The decrease in the CFU/μl values in [RNQ+] and Δrnq1 cells with Sup35PD-GFP overproduction reflects the death of cells accumulating Sup35PD-GFP amyloid. **D.** Time-lapse dynamics of necrotic cell death rate, measured as a percentage of cells positively stained with propidium iodide (PI). The cell suspensions were incubated with 2 µg/ml PI solution for 10 min and then the PI signal was measured in 10000 cells (per single time point) using a flow cytometer. The mean percentage of PI-positive cells was calculated as the average of 3 independent biological replicates. Error bars show SEM. **E.** Proportion of [PSI+] clones formed in the cell suspension with 1-day long overproduction of either Sup35PD-GFP or Sup35PD(ΔR)-GFP. Sup35(ΔPD)-GFP, a non-polymerising Sup35 mutant, was overproduced as a negative control to assess the background level of [PSI+] prion formation. The data are expressed as mean ± SEM. * p < 0,05, N.S. – non-significant (2-tailed Student’s t-test). **F.** Proportion of [PSI+] clones formed in the suspensions of [rnq-] cells without overproduction of Sup35PD-GFP, incubated over 1 and 7 days in a citrate-phosphate buffer at pH either 5.5 or 7.5. At indicated time-points, cells were streaked at the plates with and without adenine, and proportions of Ade+ clones were calculated in four independent biological replicates. As a negative control, the same experiment was performed with a strain where wild-type Sup35 was deleted and replaced by LLPS-deficient Sup35 mutant with deletion of a.a. 65-112 (indicated as ΔR), expressed from centromeric plasmid under the control of SUP35 promoter. The data are expressed as mean + SEM; * p < 0,05, N.S. – non-significant (two-tailed Student’s t-test).

To estimate the rates of [PSI+] prion formation during the aging of Sup35PD-GFP condensates, we took advantage of the ade1-14 mutation-based prion detection system, which allows the selection of [PSI+] positive cells by their ability to grow and form colonies on the solid medium lacking adenine. To our surprise, we found that even one day of Sup35PD-GFP overproduction in Δrnq1 cells resulted in a decent increase in the proportion of prion-positive clones compared to the negative control (Δrnq1 cells overproducing the non-aggregating construct Sup35(ΔPD)-GFP, Figure 4B). For comparison, in the [RNQ+] strain with overproduced Sup35PD-GFP (the amyloid cross-seeding), approximately half of the cells were converted to the [PSI+] state at this time point. Further aging of the Sup35PD-GFP condensates (3-7 days of incubation) was accompanied by a significant increase in the proportion of [PSI+] clones, reaching the level of approximately 0.3% prion-positive clones on the 5th day of overproduction. In the negative control, the prion formation rate did not exceed 7·10^-6^ throughout the incubation period. However, further incubation of the cells with Sup35PD-GFP condensates (up to day 7) did not result in an additional increase in [PSI+] positive cells. In contrast, a strong and rapid decrease in [PSI+] cell abundance was observed in the [RNQ+] strain, with the decline starting after day 1 of Sup35PD-GFP overproduction. We hypothesised that such a phenomenon was a consequence of the preferential mortality of cells loaded with Sup35PD-GFP amyloids, emerged via both LSPT (to a lesser extent) and cross-seeding by [RNQ+] prions (to a greater extent). Indeed, normalised colony forming unit (CFU) numbers decreased more rapidly in [RNQ+] cells (where Sup35PD-GFP amyloids emerge most rapidly), more slowly in Δrnq1 cells with Sup35PD-GFP condensates, and did not decrease in negative control cells with overproduction of non-amyloidogenic Sup35-based construct (Figure 4C). It appears that such amyloid-induced cell death is mostly non-necrotic in nature, as the number of propidium iodide (PI)-positive cells did not increase or increased only slightly over the course of cell incubation (Figure 4D).

To demonstrate that the formation of [PSI+] prions in Δrnq1 cells with Sup35PD-GFP overproduction is related to their phase separation, we aimed to compare the rate of prion formation in such cells with those with overproduction of the LLPS-deficient construct Sup35PD(ΔR)-GFP. To do this, we chose a time point in our time course experiments with a sharp contrast in the proportions of cells with condensates - one day after the start of overproduction. In Δrnq1 cells with Sup35PD(ΔR)-GFP overproduction, the rate of [PSI+] formation did not exceed the background level (measured on cells with Sup35(ΔPD)-GFP overproduction), whereas the overproduction of Sup35PD-GFP results in an approximately 38-fold increase (Figure 4E). Conversely, in the cells with a pre-existing template for amyloid conversion ([RNQ+] prion), overproduction of either of these constructs has led to similar levels of [PSI+] formation - meaning that Sup35PD(ΔR)-GFP amyloid can interact with WT Sup35 and convert it to prion form to nearly the same extent as Sup35PD-GFP. We can therefore conclude that, in the absence of amyloid cross-seeding, phase-separated Sup35PD-GFP assemblies appear to be a key intermediate for [PSI+] prion formation.

### LLPS of wild-type Sup35 protein may contribute to [PSI+] prion formation in nearly-physiological conditions

To investigate whether [PSI+] prion formation via Sup35 condensation could be a physiologically relevant phenomenon for yeast, we aimed to estimate the correlation between LLPS and prionisation of an endogenous full-length Sup35 expressed in the cells at physiological levels. Consistent with previously published data (Franzmann et al., 2018), the endogenous full-length Sup35-GFP phase separates more extensively in the cells incubated in the pH 5.5 buffer than in the cells incubated at pH 7.5 (Figure S4B). At the same time, incubation of [psi-] [rnq-] cells with wild-type endogenous Sup35 protein at pH 5.5 for 7 days resulted in an approximately 7-fold increase in [PSI+] formation compared to the same cells incubated at pH 7.5 (Figure 4F). This increase was not observed for cells in which wild-type Sup35 was deleted and replaced by the episomal allele Sup35Δ66-112 (i.e. the analogue of the LLPS-deficient Sup35PD(ΔR)-GFP construct). Due to the absence of [RNQ+] prion, this type of spontaneous [PSI+] formation could not be explained by the previously described mechanism (Speldewinde et al., 2017), and the rate of [PSI+] formation positively correlates both with the proportion of cells carrying Sup35-GFP condensates (affected by pH) and with the increment of incubation time. Therefore, it appears that LLPS of wild-type Sup35 may contribute to spontaneous [PSI+] formation in the cells carried under mildly acidic conditions.

### Sup35PD-GFP amyloids formed via Liquid-to-Solid Phase Transition have an unusual conformation and biophysical properties

In the process of selecting [PSI+] prions from the population of cells with aging Sup35PD-GFP condensates (Figure 4B), we noticed that the majority of selected [PSI+] isolates have a white or light pink colony colour phenotype, which is usually characteristic of strong [PSI+] variants. In contrast, according to published data, Sup35NM overproduction in [RNQ+] cells usually results in the predominant formation of weak [PSI+] isolates (associated with dark-pink colony colour phenotype; Sharma and Liebman, 2013). As the [PSI+] associated color phenotype may be affected by different modifiers, we decided to use a recently developed genetic test to characterize a spectrum of nascent [PSI+] isolates by the ‘strength’ of their phenotype (Dergalev et al., 2019). This test clearly distinguishes strong and weak class members by their opposite response to overexpression of the SUP35 and HSP104 genes. By crossing the [psi-] tester strain (with a multicopy plasmid carrying either SUP35 or HSP104 gene) with a nascent [PSI+] isolate, we found that about 94% of the [PSI+] isolates formed by LSPT of Sup35PD-GFP condensates (Δrnq1 strain, 7-day incubation) belong to the strong class, compared to only 36% of those formed via amyloid cross-seeding ([RNQ+] strain, 24-hour incubation) (Figure 5A). We have also shown that such a bias in the strong/weak variant ratio is not an artefact of either prolonged cell incubation in stationary phase (with hypothetical competition between strong- and weak-type folds formed in the same cell) or the methodology of prion detection (based on selection of cells that can grow faster in medium without adenine).

**Figure 5.**
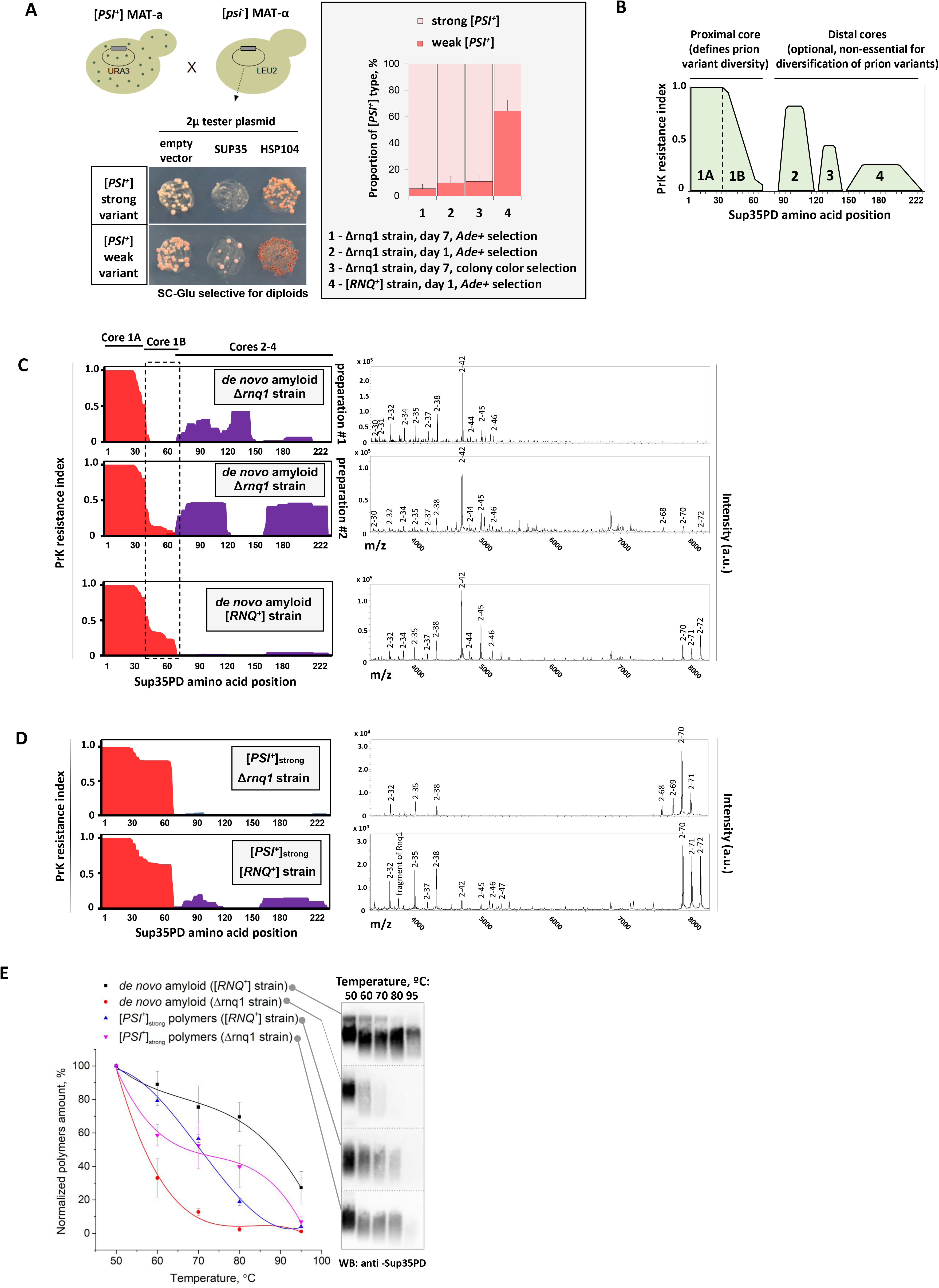
Sup35PD-GFP amyloids formed via LSPT differ from those formed via amyloid cross-seeding. **A.** Screening for proportion between strong and weak variants among [PSI+] clones emerged through either Sup35PD-GFP condensates aging (Δrnq1 strain, 7-day cell incubation, n = 95 isolates) or heterologic amyloid cross-seeding by [RNQ+] prion ([RNQ+] strain, 1-day cell incubation, n = 73 isolates). [PSI+] clones were selected as the cells with GuHCl-curable Ade+ phenotype. The tested clones carrying URA3 plasmid (pYES2-Sup35PD-GFP) were mated with the tester strain (64-D694 [psi-] [rnq-]) carrying either one of tester 2µ plasmid (YEplac181-Sup35 or YEplac181-Hsp104, LEU2 marker), or “empty” vector YEplac181. Diploids were selected on SC-Glu medium lacking uracil and leucine. To rule out a possible selective advantage for strong [PSI+] variants in case of Δrnq1 strain, we tested in the same way 36 Δrnq1 [PSI+] clones selected at 1st day of Sup35PD-GFP overproduction as Ade+ colonies, and 9 Δrnq1 [PSI+] clones selected at 7th day of Sup35PD-GFP overproduction as a white/pink colonies on YPD-red medium. **B.** An integral scheme of PrK-resistant cores composition of Sup35PD-GFP prion fibrils, generalized from the analysis of more than 20 [PSI+] isolates (Dergalev et al., 2019). N-terminal (proximal) Core 1 is ubiquitous, defines the variant-specific prion phenotype (either strong or weak), and can be subdivided into Core 1A (fully PrK-protected) and Core 1B (partially PrK-protected). Distal Cores 2-4 are optional, can be presented in different combinations and do not have an apparent effect on the variant-specific phenotype. **C-D.** PrK-resistant cores of either de novo formed bulk Sup35PD-GFP amyloids (**C**), or Sup35PD-GFP prion polymers from selected [PSI+] isolates (**D**), emerged in either Δrnq1 or [RNQ+] cells. Sup35PD-GFP fibrils, sonicated and isolated from yeast cell lysates in the presence of sarcosyl, were treated for 1 hr with 25 μg/ml proteinase K, and proteinase K-resistant peptides were analyzed with MALDI-TOF. The left panels show PrK resistance index profiles, and the right panels show a part of the MALDI-TOF mass spectra (linear mode) containing the Core 1 peptides. Unlabelled peaks represent parts of Cores 2-4. Note that the N-terminal methionine of Sup35PD-GFP has been completely removed and replaced with an acetyl group in the yeast cells, and therefore all Core 1 peaks start from the amino acid residue №2. **E.** The thermal stability assay for isolated Sup35PD-GFP amyloid polymers. Amyloid samples were incubated with SDD-AGE sample buffer (containing 2% SDS) for 8 min at specified temperatures and then loaded onto SDD-AGE. The integrated intensities of the Sup35PD-GFP amyloid smers were quantified using FIJI and plotted against temperature. Each sample of isolated amyloid was tested in 3 technical replicates and data are expressed as mean ± SEM.

Such a dramatic difference in the proportion of strong/weak [PSI+] variants may reflect the different conformations of amyloid fibres (or different distributions of conformations in the pool of amyloid fibrils), depending on the mechanism of their de novo formation in yeast. To investigate their organisation, we applied our recently developed protocol for mapping the densely packed, proteinase-resistant fibril regions in ex vivo isolated amyloids (Dergalev et al., 2019). In the original paper, we showed that Sup35 amyloid tends to have up to four proteinase-resistant regions (so-called “cores”), with the architecture of Core 1 (that typically spanns from 2 to up to 72 a.a. residues) being the most important correlate of the genetic properties of such amyloids in yeast (Figure 5B). We compared an architecture of Sup35PD-GFP amyloid formed either by amyloid cross-seeding ([RNQ+] strain, 24h Sup35PD-GFP overproduction) or by LSPT (Δrnq1 strain, 7d Sup35PD-GFP overproduction). In two of the four preparations of LSPT-derived amyloid, Core 1 was significantly shorter than in the cross-seeded amyloid (2-47 a.a. and 2-72 a.a., respectively) and lacked Core 1B (Figure 5C). In the other two preparations, the entire core 1 extended up to 72 aa, but core 1B was less PrK resistant than the corresponding region in the cross-seeded amyloids.

We further mapped the architecture of Sup35PD-GFP prion polymers extracted from [PSI+] isolates selected from strains with either amyloid cross-seeding ([RNQ+] strain) or LSPT (Δrnq1 strain). To this end, Sup35PD-GFP was overproduced for 18 hours in a set of [PSI+] isolates obtained from either [RNQ+] or Δrnq1 cellular background (two strong and two weak isolates from each strain, the strength was confirmed by a genetic test - Figure 5A). Strong [PSI+] isolates selected from both backgrounds generally had similar Core 1 architecture Figure 5D). Among studied weak isolates, one of LSPT-derived ones also had similar Core 1 patterns with amyloid cross-seeding-derived weak isolate, while the second one had almost absent Core 1B, which is quite similar with Core 1 structure of the bulk LST-derived amyloid (Figure S5).

To prove that observed difference in the Core 1 architecture reflect the prominent difference in the physical properties of studied amyloids, we performed the thermal stability assay. As expected, the LST-derived de novo amyloids (preparation #1) were shown to be much less thermal resistant that ones derived by the amyloid cross-seeding. In contrast, the amyloids purified from two strong [PSI+] isolates emerged via either LSPT or cross-seeding, had similar profiles of thermal resistance, which correlates with their similar Core 1 structures.

### Transient liquid condensates drive the amyloid formation and prion switch for other yeast prionogenic proteins

To generalize the observations made with the Sup35-based construct, we then asked whether LLPS might be an important trigger for the emergence of heritable and non-heritable amyloids formed by other prionogenic yeast proteins. More than 40 yeast protein PDs (selected by sequence similarity to the known prion proteins) have been shown to form amyloids when overproduced in yeast cells carrying [RNQ+] prion (Alberti et al., 2009). Some of them have also been shown to confer the heritable epigenetic switch of the corresponding endogenous protein into a prion form. However, their propensity to aggregate and form prions in the absence of the [RNQ+] prion template has not been addressed. Based on our observations with Sup35, we expected that at least some of these proteins, when overproduced in Δrnq1 cells, would form amyloids and possibly prions through aging of initially formed liquid-like condensates. On the other hand, the second class of PDs, which would not phase separate even at high concentrations in vivo, would not be able to form amyloids in such a template-free manner.

To test our assumptions, we select a set of four yeast prionogenic proteins. Ure2 and Rnq1 are among the “classical” yeast prion proteins (Wickner, 1994; Derkatch et al., 1997); Mot3 has been identified as a prionogenic protein in a large-scale screen, and Sap30 has been shown to form amyloids in the [RNQ+] strain (Alberti et al., 2009) and to adopt a prion-like state (Du et al., 2015). Ure2, Mot3 and Sap30 proteins, which possess N-terminal PDs, were cloned into chimeric constructs with the C-terminally fused Sup35M domain as a spacer (with deletion of the negatively charged cluster as described above for the Sup35PD-GFP construct) followed by eGFP (Figure 6A). Because the Rnq1 protein has several putative amyloidogenic stretches distributed throughout the protein (with a tendency to be located in the C-terminal and middle parts of the polypeptide (Stein and True, 2014)), we constructed an N-terminal fusion of the full-length Rnq1 sequence with the sfGFP reporter. All four constructs were cloned into a multicopy pYES2 vector under the control of the GAL1 promoter (Figure 6A). Based on the observation that a high content of glutamine (but not asparagine) residues tends to promote polypeptide chain LLPS (Halfmann et al., 2011), we expected that overproduction of Rnq1- and Mot3-based constructs (with Q/N-rich sequences) in Δrnq1 cells would lead to LLPS of these proteins, whereas overproduction of Ure2- and Sap30-based constructs (with N-rich PDs) would not lead to phase separation (Figure S6A). In addition, we were interested in whether ongoing amyloid aggregation and possibly prion formation of these constructs in the long-term incubated cells would correlate with their ability to form condensates.

**Figure 6.**
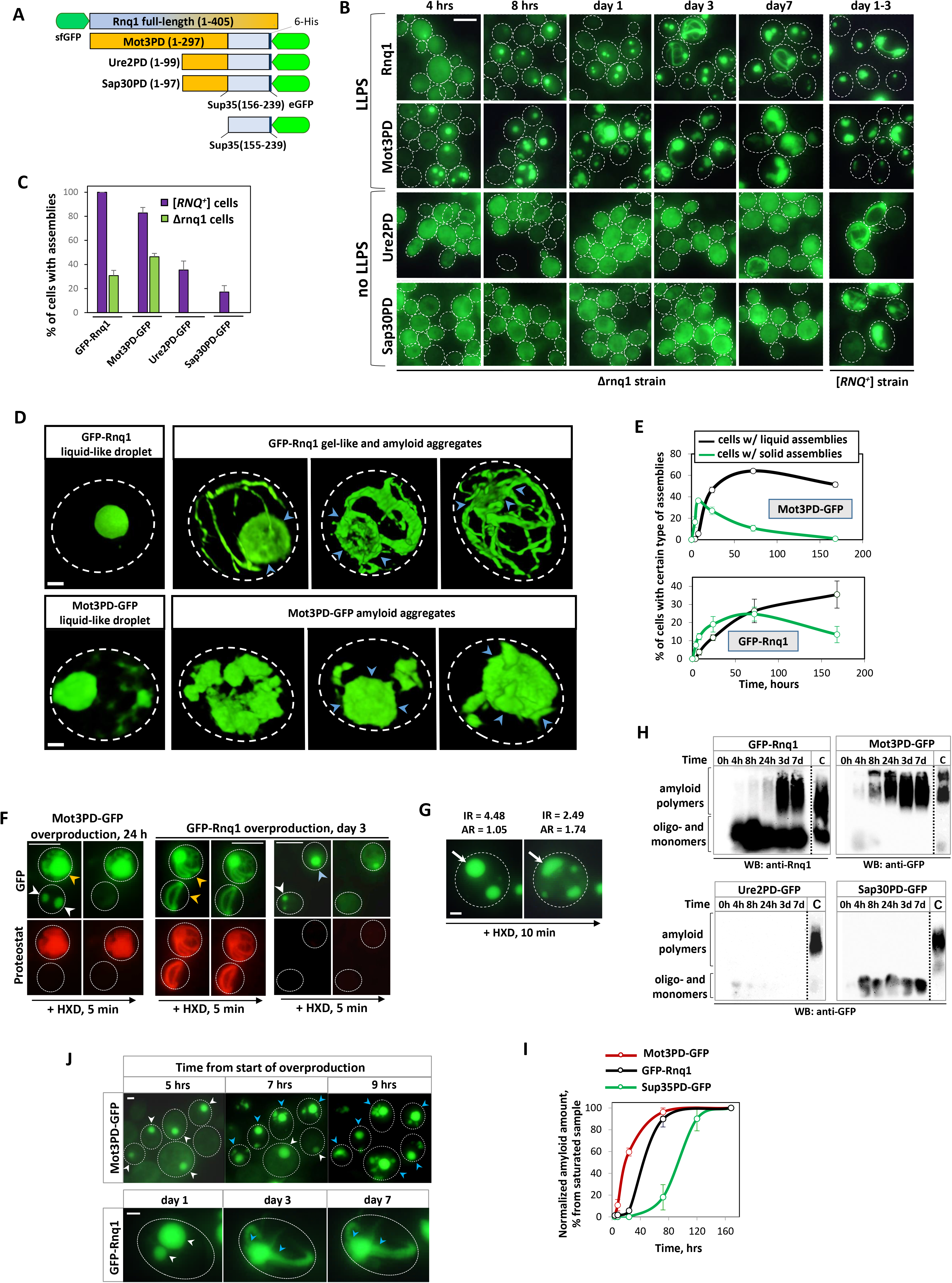
GFP-Rnq1 and Mot3PD-GFP form amyloids through LSPT. **A.** Schematic representation of the chimeric constructs used in this work. Orange colour highlights putative prion-forming regions, blue colour highlights putative non-prion-forming regions. The fragments of the Sup35 M domain (e.g. 156-239) were used as spacers between the prion-forming region and the GFP tag in certain chimeric constructs. **B.** Time-lapse maximum projected images of Δrnq1 cells overproducing either GFP-Rnq1, Mot3PD-GFP, Ure2PD-GFP or Sap30PD-GFP proteins. [RNQ+] cells overproducing the same constructs (time = 1 day for GFP-Rnq1, Mot3PD-GFP and Ure2PD-GFP, and 3 days for Sap30PD-GFP construct) are shown for comparison. Scale bar is 5 μM. **C.** Quantification of the proportion of cells with assemblies. Note that all four proteins assemble in the cells with [RNQ+] prion (i.e., via amyloid cross-seeding), but only two of them form foci in GuHCl-treated Δrnq1 cells (i.e., via LLPS). The data represent mean + SEM (n = 3). **D.** Volume snapshots of 3D-rendered super-resolution SIM images of fresh and aged condensates of GFP-Rnq1 and Mot3PD-GFP proteins: liquid-like condensates (1 day incubation for GFP-Rnq1 and 8 h incubation for Mot3PD-GFP) and solid-like aggregates (3-5 days incubation for GFP-Rnq1 and 1 day incubation for Mot3PD-GFP). Blue arrows indicate solidified bodies emerging from liquid-like droplets. **E.** Time-lapse dynamics of liquid- and solid-like assemblies formed in Δrnq1 cells by GFP-Rnq1 and Mot3PD-GFP proteins. At indicated time points, the suspensions were imaged in presence or absence of 10% HXD. As HXD disrupt initially formed liquid-like condensates of Mot3PD-GFP and GFP-Rnq1, but not solid-like hydrogels and amyloid bodies, the cells with intact aggregates under HXD treatment were counted as “cells with solid-like assemblies”. The proportion or cells with liquid-like assemblies was calculated as difference between the proportion of cells with any type of assembly (either liquid- or solid-like, imaged in absence of HXD) and proportion of cells with solid-like assemblies in each replicate. The experiment was performed in 3 independent biological replicates, the data presented as mean ± SEM. **F.** Maximum intensity projections of Δrnq1 cells with Mot3PD-GFP and GFP-Rnq1 assemblies stained with amyloid-specific fluorescent dye proteostat, and probed with 10% 1,6-hexanediol solution. Alike freshly-formed droplets (white arrows) and aged hydrogels (blue arrows), the linear aggregates and aggregates with outgrowths (orange arrow) stain with proteostat. Scale bar is 5 μM. **G.** GFP-Rnq1 hydrogels can change their shape and redistribute part of their interior protein into the surrounding cytosol under the treatment with 10% solution of 1,6-hexanediol. White arrow indicate GFP-Rnq1 hydrogel particle that became more elongate (aspect ratio (AR) increases from 1.05 to 1.74) and less enriched with GFP-Rnq1 protein (GFP intensity ratio (IR) inside/outside hydrogel decreases from 4.48 to 2.49). Scale bar is 1 μM. **H.** Time-lapse SDD-AGE analysis of amyloid polymers formation dynamics in Δrnq1 cells with overproduction of either GFP-Rnq1, Mot3PD-GFP, Ure2PD-GFP or Sap30PD-GFP constructs. As a positive control for amyloids mobility (denoted as «C»), we used lysates of [RNQ+] cells with 1-day-long overproduction for Rnq1-, Mot3- and Ure2-based constructs, and 3-day-long overproduction for Sap30-based construct. Each experiment was performed in 3 independent biological replicates, and representative images are shown. **I.** Quantification of amyloid accumulation dynamics for Sup35PD-GFP, GFP-Rnq1 or Mot3PD-GFP constructs in Δrnq1 cells. The data represent mean ± SEM. Note that SDD-AGE immunoblot images for Sup35PD-GFP amyloid accumulation are shown in Figure 3G. **J.** Time-lapse imaging of liquid-to-solid phase transition in condensates formed by either Mot3PD-GFP or GFP-Rnq1 proteins in the single cells. Δrnq1 cells with a corresponding plasmid were grown to a stationary phase in SC-Glu medium, diluted 4-fold in SC-Gal medium, incubated for either 4 (Mot3PD-GFP) or 24 (GFP-Rnq1) hours, streaked to the single cells on the agar pad with pH buffered at 6.5, and incubated at 30C° in a wet chamber. Cell images were taken at the indicated time points. Liquid-like assemblies (indicated by white arrows) rearrange into solid-like bodies or evolve the linear outgrowth (blue arrows). Scale bar is 1 μM.

We overproduced these four constructs in stationary cultures of the GuHCl-cured Δrnq1 strain on the SC-GAL medium. Except for GFP-Rnq1, all other constructs reached higher expression levels than Sup35PD-GFP under the control of the GAL1 promoter (Figure S1A). Consistent with previously published data (Kroschwald et al., 2015), the overproduced GFP-Rnq1 construct initially (4-8 hrs of overproduction) forms large, quasi-spherical foci that rapidly dissolve after the addition of 10% HXD and can thus be classified as liquid-like condensates. A similar observation was made with the Mot3PD-GFP construct (Figure 6B). The liquid-like droplets of GFP-Rnq1 and Mot3PD-GFP are generally more enriched with the constitutive protein (relative to a “dilute” phase in the cytosol) than Sup35PD-GFP droplets (median IR = 3.07, 2.38 and 1.55, respectively, Figure S6B). On the other hand, both Ure2PD-GFP and Sap30PD-GFP constructs did not form visible aggregates in the early stages of overproduction (0-3 days; Figure S6I). However, Ure2PD-GFP formed the ring- or rod-like, HXD-insoluble, proteostat-positive aggregates in a small proportion (up to 0.5±0.2%) of cells in the late stages of cell incubation (3-7 days; Figure S6F, S6I).

At the 8th hour of Mot3PD-GFP overproduction, we observed the emergence of the second class of assemblies, which were either globular, linear or irregular in shape, depleted of almost all constituent proteins in the cell, positively stained with proteostat, and completely (or partially) resistant to HXD treatment (Figure 6B, 6D, 6F). All these properties are consistent with the amyloid-like nature of these assemblies. After 24 hours of overproduction, the majority of Mot3PD-GFP assemblies were found to be of the second class. In analogy to Sup35 gels and rings, we hypothesized that these solid-like aggregates might mostly arise as derivatives of initial liquid-like assemblies that are very transient and readily switch to the amyloid state. To prove this, we initiated Mot3PD-GFP overproduction by plating cells on the solid SC-Gal medium pad and followed aggregation events in a number of individual cells. These observations confirmed that first class assemblies (liquid-like droplets) tend to transform into second class assemblies (solid-like aggregates), which often have linear or irregular protrusions and sequester almost all of the diffuse Mot3PD-GFP in the cell.

We observed a similar but slower LSPT process for GFP-Rnq1 condensates. Typically, it was expressed as the formation of linear, sometimes ring-shaped protrusions (growing from the initial spherical condensate), exclusively ring-shaped aggregates, or spherical solid-like bodies (Figure 6B, 6D, 6F, 6J). Similar to Sup35PD-GFP (and in contrast to Mot3PD-GFP), GFP-Rnq1 liquid-like droplets can age into gel-like condensates that are partially soluble by HXD treatment (Figure 6F, 6G). These hydrogels are either proteostat-negative (Figure 6F) or partially stained, with a tendency for the periphery to be more stained than the interior (Figure S6E). Probably as a result of this tendency of amyloid growth near the condensate surface, we can also observe solid-like bodies with hollow morphology (Figure 6D, S6F).

To investigate the dynamics of the liquid-like and solid-like assemblies of the studied constructs, we determined the time-resolved proportions of cells with either HXD-sensitive or HXD-resistant (partially or completely) assemblies (Figure 6E). The proportions of cells with HXD-sensitive (i.e., liquid-like) assemblies peak at either 8 hours (for Mot3PD-GFP) or 72 hours (for GFP-Rnq1) and then steadily decline. The proportions of cells with HXD-resistant (i.e. solid or gel-like) aggregates of both proteins increase in antiphase with respect to cells with HXD-sensitive assemblies, confirming that the latter act as a transient intermediate.

To confirm the accumulation of amyloid forms of Rnq1 and Mot3PD (but not Ure2PD and Sap30PD), we performed SDD-AGE with time-lapse samples from stationary yeast cultures overexpressing each of these constructs. For all four chimeras, this biochemical analysis confirmed the picture painted by fluorescence microscopy: the SDS-resistant polymers of GFP-Rnq1 and Mot3PD-GFP appeared from the first day of overproduction and gradually increased in number as the incubation progressed, whereas Ure2PD-GFP and Sap30PD-GFP did not form detectable amounts of amyloid (Figure S6I). Of note, unlike LSPT-derived Sup35PD-GFP and Mot3PD-GFP macroaggregates, GFP-Rnq1 amyloid macroaggregates appeared to contain predominantly large fibrils, which did not enter the 1.8% agarose gel - and we therefore had to sonicate such samples (Figure S6J).

To assess the potential prion switch of endogenous Rnq1, Mot3 and Ure2 proteins, we developed a novel bi-prion reporter system in yeast (Figure 7A; see description in the Materials and Methods section), based on the so-called “multicopy suppression” phenomenon (Salnikova et al., 2005), in which multicopy expression of the SUP35 gene in cells carrying another prion results in a transient nonsense-suppressed phenotype on medium lacking adenine. For cells overproducing the GFP-Rnq1 and Mot3PD-GFP constructs, we observed a significant increase in the proportion of Ade+ clones, whereas Ure2PD-GFP overproduction did not show such an effect (Figure 7B). Several dozen Ade+ clones selected after GFP-Rnq1 and Mot3PD-GFP overproduction were confirmed to be [PRION+] clones by guanidine assay (Figure S7C). The level of [MOT+] prion appearance was compatible with that induced by Sup35PD-GFP LSPT, reaching approximately 0.2% of the total cell population (approximately a 600-fold increase over the basal level). The level of [RNQ+] prion formation at the late stage of cell incubation was about 20-fold lower, despite the fact that overproduced GFP-Rnq1 underwent a very intense LLPS and LSPT. This could be explained by the high toxicity of GFP-Rnq1 overproduction in cells with formed [RNQ+] prion - however, the decrease in CFU number in cells overproducing GFP-Rnq1 was not dramatically higher than in cells overproducing Mot3PD-GFP (Figure 7D). Another line of explanation may be based on our observation that the fibrils formed by LSPT of GFP-Rnq1 are much larger and less fragmentable, as assessed by their mobility on the agarose gel (Figure S6J), which may reduce their heritability.

**Figure 7.**
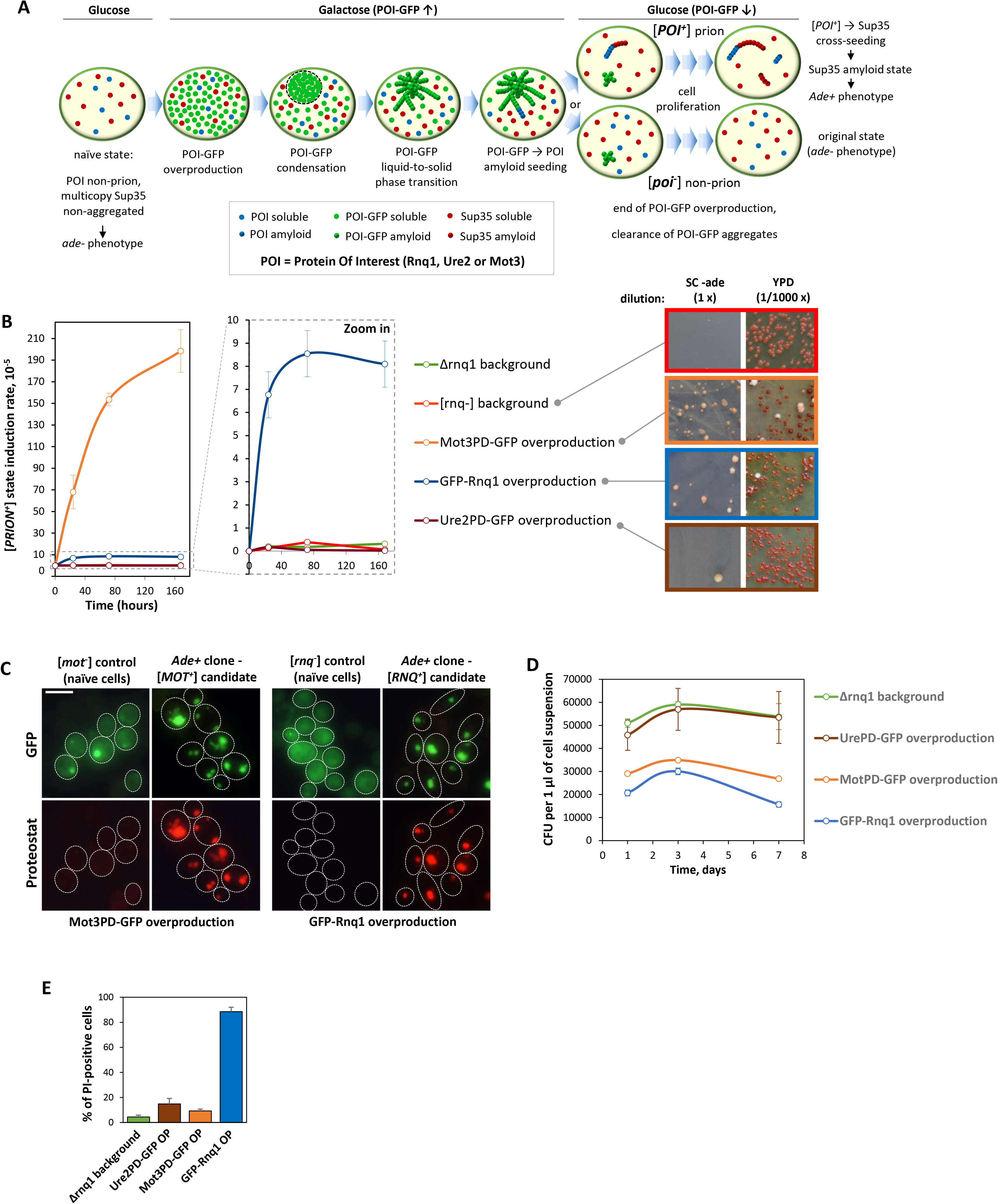
GFP-Rnq1 and Mot3PD-GFP can form prions through LSPT. **A.** Scheme of phenotypic detection of [RNQ+], [MOT+] and [URE+] prions arising in a strain with no pre-existing conformational template. When the protein of interest is switched to the prion form, it cross-seeds the amyloid polymerisation of moderately overproduced Sup35 protein, conferring on cells the ability to establish the Ade+ phenotype on medium lacking adenine. **B.** Time-resolved measurement of [PRION+] clones induction level upon GFP-Rnq1, Mot3PD-GFP or Ure2PD-GFP overproduction, scored as Ade+ clones in the genetic assay depicted in Figure 8A. GFP-Rnq1 construct was overproduced in [rnq-] strain with multicopy SUP35 gene, Mot3PD-GFP and Ure2PD-GFP constructs were overproduced in Δrnq1 strain with multicopy SUP35 gene. As a negative control (i.e. background level of [PRION+] clones formation), Sup35(ΔPD)-GFP construct (in fact, non-aggregating fragment of Sup35 M domain tagged with GFP) was overproduced in the same genetic backgrounds over the same time. **C.** GuHCl-curable Ade+ clones, selected after the transient overproduction of either GFP-Rnq1 or Mot3PD-GFP proteins, possess [RNQ+] and [MOT+] prions respectively, as revealed by GFP-Rnq1 or Mot3PD-GFP aggregation patterns. Maximum intensity projections of cells with 4 hrs long overproduction of either Mot3PD-GFP (Δrnq1 strain) or GFP-Rnq1 (RNQ1 strain) proteins, stained with amyloid-specific fluorescent dye proteostat are shown. Scale bar is 5 μM. **D.** Time-lapse dynamics of the viable cell density, measured as the number of colony-forming units (CFU) yielded from 1 μl of cell suspension streaked on YPD plates. The mean CFU/μl values were calculated for three independent biological replicates. Error bars show SEM. The decrease in the CFU/μl values in the cells with Mot3PD-GFP and GFP-Rnq1 overproduction reflects the death of cells accumulating amyloid form of these proteins. **E.** Cells with 7-day-long overproduction of Mot3PD-GFP and Ure2PD-GFP die mostly via apoptosis-like mechanism, while cells with overproduction of GFP-Rnq1 construct die via necrosis-like mechanism. The cell suspensions were incubated with 2 µg/ml propidium iodide (PI) solution for 10 min and then 400-700 cells per suspension were subjected to fluorescent microscopy. The mean percentage of PI-positive cells was calculated as the average of 3 independent biological replicates. Error bars show SEM.

## Discussion

### Liquid-liquid phase separation of the constructs based on yeast prionogenic proteins

In the present work, we investigated the in vivo phase behaviour of five GFP-tagged chimeric proteins containing an amyloidogenic domain of either Sup35, Rnq1, Ure2, Mot3 or Sap30 proteins. These constructs were overproduced in yeast cells and reached intracellular concentrations 2-3 orders of magnitude higher than physiological levels of corresponding wild-type proteins. According to numerous reports, overproduction of similarly constructed proteins in cells containing another amyloid-based prion (especially, [*RNQ+*]) has usually led to its fast amyloidization via a cross-seeding mechanism (Alberti et al., 2009; Bardill and True, 2009). To avoid this circumstance, we performed the majority of such experiments in the yeast strain devoid of a pre-existed amyloid template - i.e., in the GuHCl-cured cell line, usually with the deletion of RNQ1 gene.

In previous works, the phase separation of Sup35 protein was characterized as dependent on electrostatic interactions, as higher salt concentrations generally inhibit this process (Franzmann et al., 2018). However, our experimental data suggest that weak hydrophobic interactions also make a great contribution to Sup35 condensates formation, as the latter were dissolved by a chemical agent disruptive for such interaction type (Figure 1D). Similar types of interactions are likely underlie the condensation of GFP-Rnq1 and Mot3PD-GFP constructs, as their LLPS was also reversed by cell treatment with 1,6-hexanediol (Figure 6F, 6E). In addition, we have also discovered that at early stage of their formation, the condensates of all three proteins are sensitive to RNAse A treatment, which could imply a scaffold role of mRNA in their LLPS, and probably makes these condensates somewhat similar with the physiological cytoplasmic condensates of yeasts - stress-granules and P-bodies. To clarify their inter-relation, we probed the colocalization of Sup35PD-GFP, GFP-Rnq1 and Mot3PD-GFP condensates with the RFP-tagged markers of stress-granules and P-bodies. While the condensates of GFP-Rnq1 and Mot3PD-GFP colocalize with specific markers of both types of physiological condensates (Figure S6C), Sup35PD-GFP condensates contain only stress-granules marker Pab1-RFP, but not P-bodies marker Dcp2-RFP (Figure S1C). However, Pab1-RFP forms numerous separate (not enriched with Sup35PD-GFP) assemblies in cells with Sup35PD-GFP condensates, and forms less numerous granules in the cell without Sup35PD-GFP condensation (Figure S1D), we can conclude that neither of these three condensates appear to be true stress-granules or P-bodies - instead, they could be characterized as the artificial cytosolic condensates with a complex, multicomponent composition. This trait is greatly distinguishes them from in vitro reconstituted condensates (which are typically consist of the one type of protein, sometimes scaffolded with the RNA molecules) and makes their behaviour more unpredictable.

The liquid-like condensates of three studied proteins differ in number (per cell), size and density of the main constituent protein. Overproduced Sup35PD-GFP initially (8-12 hrs of overproduction) forms small numerous (up to 10-20 condensates per cell) foci which tend to fuse in several larger complexes, while GFP-Rnq1 and Mot3PD-GFP typically form from 1 to 3 larger condensates per cell. Sup35PD-GFP protein has the lowest enrichment in its freshly-formed condensates (the median value ∼1.6 fold), Mot3PD-GFP has higher enrichment (∼2.4 fold), and GFP-Rnq1 is the most enriched in its condensates (∼3.1 fold, Figure S6B). We can also note that Mot3PD-GFP droplets emerge at more early stages of galactose-induced overproduction (that ones of Sup35PD-GFP and GFP-Rnq1), with ∼40% Mot3PD-GFP overproducing cells contain droplets at 8th hour after galactose addition (Figure 6E). This may indicate that Mot3PD-GFP is likely to have a lower C_sat_ for condensate formation compared to Sup35PD-GFP and GFP-Rnq1.

We can also notice, that the observed production levels of all three constructs capable to phase separate in vivo (Sup35PD-GFP, Mot3PD-GFP and GFP-Rnq1) are instantly lower that those measured for four non-phase-separating constructs used in this work (Sup35PD(ΔR)-GFP, Sup35(ΔPD)-GFP, Ure2PD-GFP and Sap30PD-GFP), and the difference may reach an about 3,8 folds (for the Ure2PD-GFP/GFP-Rnq1 pair, Figure S1A). Possibly, the formation of liquid-like condensates by such constructs is, in general, somewhat unbenificial (or even toxic) to yeast, exerting a selective pressure towards reducing the production level of the phase-separating constructs.

### Liquid-to-Solid Phase Transition of Sup35PD-GFP, Mot3PD-GFP and GFP-Rnq1 condensates

The liquid condensates composed of Sup35PD-GFP and GFP-Rnq1 constructs transit to solid-like, amyloid-containing assemblies more or less gradually. In majority of cells with Sup35PD-GFP overproduction, its liquid condensates reorganizes into assemblies that can be referred to as hydrogels - the aggregates with the properties somewhat intermediate between liquid-like droplets and amyloid bodies: semi-solubility in HXD (Figure 2J); roughly 50% of constituent Sup35PD-GFP is mobile in the FRAP assay (Figure 2J); static, irregular shape, knobbing morphology. In the “cell burst” assay (Video 2I), we can see that after the hydrogel release in extracellular space, a part of the constituent protein dissipate (as it was in case of liquid-like condensate), while the rest of Sup35PD-GFP remain to be an integrative body. The biochemical experiments showed that significant proportion of Sup35PD-GFP in the cells containing aged hydrogels (days 3-7 of cell incubation) is in the form of SDS-insoluble amyloid polymers (Figures 2G and 2K). Finally, CLEM showed that aged hydrogel is more electron-dense area (comparing with the rest of the cytosol) containing both individual fibrils and fibril bundles (Figure 3). Therefore, combining all these data, we may conclude that Sup35PD-GFP hydrogels have an amyloid “sceleton” surrounded by the more dense and viscous liquid component (comparing with initially formed liquid condensates). This non-amyloid component of the hydrogels appears to act as a protective shell for amyloid fibrils, as the hydrogels were not stained with proteostat in the intact cells (Figure 2E), but were stained in the cell lysates (Figure 2F) and in cells pretreated with HXD (Figure S2J).

In contrast to Sup35PD-GFP hydrogels, its ring- or net-like aggregates have expressed properties of the amyloid bodies (Figure 2E, 2J), and appear to be very similar to ring-like aggregates formed by the same protein in [RNQ+] cells via the amyloid cross-seeding (Figure 2A, S1F). How could such aggregates emerge in the cells with phase-separated Sup35PD-GFP? We hypothesize that in cells with the single (or few) center of amyloid nucleation the fibrils can form a bundle, which could act as a primordium of ring-like aggregate. This bundle could them quickly grow, sequestering nearly all monomeric Sup35PD-GFP and forming a ring-/net-like pattern. However, in the majority of Sup35PD-GFP condensates, the centers of amyloid nucleation could be multiple, providing more or less even filling of the condensate volume with fibrils. In many cells we can also see the linear outgrowths protruding from the gel-like condensates (Figure 2B, 2H).

Similar to the Sup35PD-GFP condensates, GFP-Rnq1 droplets can evolve into the particles with the properties intermediate between liquid-like and all-amyloid assemblies (Figure 6G, S6E). As we can see in numbers the hollow GFP-Rnq1 particles (Figure S6E), as well as the spherical particles with only the surfase zone stained with proteostat (Figure S6E),it appears that GFP-Rnq1 amyloids tend to form more readily on the surface of the condensate than in its depth. Similar effect was described for some other amyloids (Pytowski et al., 2020). In the cell suspension with GFP-Rnq1 LSPT, the cells carrying typical amyloid rings are not very frequent, but we can often see the spherical particles (probably, the droplets “skeletonized” by the amyloid fibrils) with long linear or ring-like protrusions (Figure 6D, 6F, 6J). As GFP-Rnq1 LSPT and amyloid formation proceed faster than LSPT of Sup35PD-GFP (Figure 6I), the condensates of GFP-Rnq1 got skeletonized and rigid more quickly and conserve their spherical shape, while Sup35PD-GFP condensates got squeezed between cell membrane and growing central vacuole prior to adopt solid-like properties.

Unlike Sup35PD-GFP and GFP-Rnq1, Mot3PD-GFP assemblies appear to switch from liquid-like condensate to solid-like amyloid state quite fast and abruptly, as was visualized with the single-cell fluorescent microscopy by tracking the fate of the single assemblies (Figure 6J). Throughout all time-lapse experiments, we were not able to find Mot3PD-GFP assemblies with the properties intermediate between liquid condensates and amyloid-like particles - being it the morphological traits, instability under the 1,6-hexanediol treatment, or proteostat staining. Also, we saw a lot of cells with the hollow Mot3PD-GFP aggregates, which indicates that in these condensates amyloids have formed preferentially in the surface area.

Why LSPT of Sup35PD-GFP, GFP-Rnq1 and Mot3PD-GFP constructs proceed with such prominent difference in speed? The possible cause of such difference may originate from the different local concentrations of these three constructs within their condensates. Indeed, Sup35PD-GFP and GFP-Rnq1 constructs were produced at similar levels (Figure S1A), but the mean inside/outside concentration ratio was much higher for GFP-Rnq1 condensates (Figure S6B), resulting in its higher concentration inside its dense phase. In contrast, the mean inside/outside concentration ratios are similar for GFP-Rnq1 and Mot3PD-GFP, but the production level of Mot3PD-GFP is higher that those of GFP-Rnq1, which give us higher local concentration of Mot3PD-GFP comparing with GFP-Rnq1. In addition, the individual, sequence-based properties of Sup35PD-GFP, GFP-Rnq1 and Mot3PD-GFP in respect to amyloid formation are also likely to contribure in their individual LSPT rates.

Based on all our microscopic and biochemical observations, we can reconstruct the integral scheme summarizing all possible pathways of amyloid formation by LLPS and LSPT or three studied proteins in yeast (Figure 8A). Depending on the given protein and the different circumstances, initially formed liquid-like condensates could undergo morphological and conformational transformation leading to the formation of either ring-like aggregates, flattened hydrogels with near-surface localization, spherical hydrogels with cytosol localization, hollow amyloid bodies, amyloid bodies with linear/ring-like protrusions, and amyloid globes with irregular shape. For each or three proteins studied, we observed a specific combination of some of these pathways, represented with different frequencies in the cellular population.

**Figure 8.**
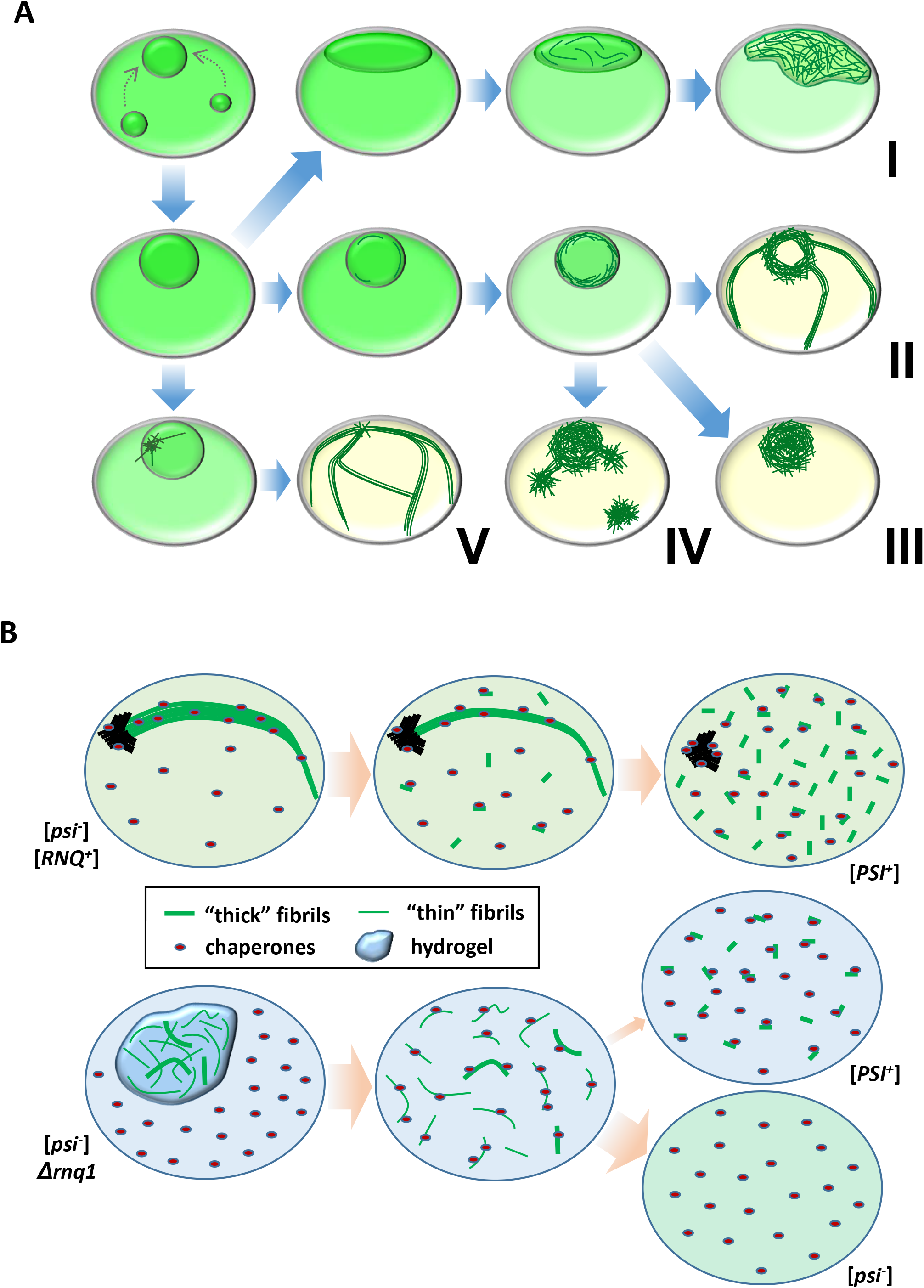
Schematic summary and conceptualization of the findings. **A.** Schematic representation of the different scenarios, or «pathways», of condensate ageing into the amyloid-containing particles. The liquid-like droplets may transform into a hydrogel containing meshwork of fibrils (I), into an amyloid-built hollow body with protrusions built by fibril bundles (II), into an amyloid-built globe (III), into an amyloid-built globe with several additional amyloid bodies and small protrusions (IV), and into a meshwork of ring or ribbon-like structures formed by fibril bundles (V). The condensates of Sup35PD usually age through I, II or V «pathways», the condensates of Rnq1 age through II, III or V «pathways», and Mot3PD condensates tend to age through II, III or IV «pathways». **B.** Model reconstructing formation of different Sup35PD-GFP amyloid conformations via either cross-seeding by [RNQ+] prion or liquid-to-solid phase transition of Sup35PD-GFP condensates. Hypothesized “thick” fibrils, that are more firm, thermodynamically stable and gave rise to [PSI+] prion, appear to be more represented in [RNQ+] cells with Sup35PD-GFP amyloid “rings”, while more fragile and unstable “thick” fibrils may appear inside aged Sup35PD-GFP condensates (hydrogels), where they could be protected from the action of cellular disaggregase machinery.

All features characterizing time-resolved phase behavior of three phase-separating (and two non-phase separating) constructs used in this work are summarized in the Table S1.

### Prion formation via Liquid-to-Solid Phase Transition of yeast prionogenic proteins

Prions are infective (transmissible, or heritable upon the cell division) protein-based agents, mostly represented by easily propagating and spreading amyloid fibrils. However, not every type of amyloids can act as a prion determinant in the living systems. Inside the cells, many type of amyloids are subjected to fragmentation by certain cellular chaperones, being it Hsp70 and VCP chaperones in mammalian cells, or Hsp104 assisted by Ssa1 and Sis1 in the yeasts (Kushnirov et al., 2021; Saha et al., 2023). To behave as a prion, the amyloid conformer should fall into a narrow interval between too strong and too weak fragmentation - because too slowly fragmented fibrils will eventually be oversized and too sparse for stable transmission into the daughter cells, and excessively fragmented fibrils could eventually be completely disaggregated to a monomer state. Therefore, despite the growing body of evidence exemplifying the pathway of amyloid formation via ageing or “maturation” of proteinaceous liquid condensates - it is still an open question, whether the amyloids of such origin could have the proper conformation to endure the multiple cycles of growth, fragmentation and transmission - or, instead, they can exist only as a part of misfolded protein deposits.

In the present work, we detected a progressive rate of [*PSI+*] prion formation upon Sup35PD-GFP overproduction, condensation and ongoing LSPT of Sup35PD-GFP condensates (Figure 4B). The importance of the condensate stage for prion formation was underlined by using a Sup35PD mutant with impaired ability to LLPS, but a nearly-intact propensity to provoke [PSI+] formation upon transient overproduction in [*RNQ+*] strain (via a cross-seeding mechanism; Figure 4E). Interestingly, that the moderate increase in [*PSI+*] formation was detected even after 1-day-long Sup35PD-GFP overproduction, even though the amyloid form of this protein was not detected by either biochemical or fluorescence microscopy tests. This may indicate that the single, sparse amyloid fibrils, which amounts is insufficient for the detection by the most methods, could likely emerge even in relatively “fresh” condensates (18-24 hrs of Sup35PD-GFP overproduction).

### Structural properties of Sup35PD-GFP amyloid formed via LSPT

The common path for amyloid and prion formation in yeast is heterologous cross-seeding with another pre-existing amyloid (usually the [RNQ+] prion). The latter probably acts as an imperfect mechanical template that physically interacts with the specific monomeric proteins and catalyzes their amyloid conversion (Derkatch et al., 2004; Salnikova et al., 2005; Vitrenko et al., 2007; Keefer et al., 2017). In this way, such a mechanical template is likely to transfer at least some of its conformationally encoded information to the newly formed amyloid fibril of another protein. If this assumption is correct, then the fibrils formed by cross-seeding would differ in structure from those formed under similar conditions but without a mechanical template. And indeed, we found that Sup35PD-GFP fibrils formed in cells with [RNQ+] prion have a different profile of PrK resistance than those formed via LSPT in cells lacking any amyloid-based prions (Figure 5C). The most important difference is the length and relative PrK resistance of the N-terminal amyloid core, which was previously shown to define the variant-specific prion properties of Sup35 amyloid (Dergalev et al., 2019). Combined with the thermal stability data of different Sup35PD-GFP amyloid types (Figure 5E), we can hypothesize that fibrils formed via LSPT are generally thinner, more fragile, and more susceptible to chaperone-mediated disaggregation than those formed via cross-seeding (Figure 8B). Such a difference may be due not only to the presence or absence of heterologous cross-seeding, but also to the situation where fibrils formed within solidifying condensates could hypothetically be fully or partially protected from the cellular disaggregase machinery (a variety of cellular chaperones that have been hypothesized to have evolved as a cellular defense system to preventing the formation and accumulation of deleterious amyloids; Wickner et al., 2022).

## Materials and Methods

### Yeast strains and cultivation conditions

Most experiments described in this manuscript were carried out on different derivatives of the 74-D694 yeast strain (MATa *ade1-14, trp1-289, his3Δ-200, ura3-52, leu2-3,112*), which has an adenine biosynthesis pathway corrupted by the nonsense mutation in the ADE1 gene. This allowed us to detect a nonsense codon readthrough (a basic phenotypic trait for [*PSI+*] prion) as a switch to *Ade+* phenotype (the ability to grow on the medium lacking adenine) and color phenotype (colony color shifting from red to white through the multiple intermediate shades, can be observed on the medium with lowered adenine content). In different experiments, 74-D694 derivatives with different combinations of deletions and epigenetic elements - [*PSI+*] [*RNQ+*], [*psi-*] [*RNQ+*], [*psi-*] [*rnq-*], [*psi-*] Δrnq1(*RNQ1::HIS3*) - were used. For the mating procedure, we used the 74-D694 strain derivative with MATα mating type. As a negative control in the experiment showing the [*PSI+*] formation without Sup35PD-GFP overproduction, we used 74-D694 derivative where genomic SUP35 gene was deleted (*SUP35::TRP1*), and SUP35 allele with deletion of a.a. 66-112 (Sup35ΔR) was introduced with pRS313 centromeric vector (under the control of SUP35 promoter).

The experiments studying the colocalization of the proteins of interest with cellular compartments, as well as the experiment visualizing LLPS of the full-length Sup35 protein, were performed on the BY4741-based strains (MATa *his3-1 leu2-0 met15-0 ura3-0*) derived from the whole-genome C-terminal GFP-fusion library collection (Huh et al., 2003). For the Sup35 LLPS experiment, the cell line, where the genomic Sup35 was C-terminally tagged with S65T-GFP, was grown for ∼50 generations on YPD plates with 4 mM GuHCl, to cure any Hsp104-dependent prion that could pre-exist in this strain. For colocalization experiments, the strains carrying either DCP2-GFP, PAB1-GFP, or HSP42-GFP genes, were transformed by a cassette that replaces S65T-GFP with tagRFP in the respective fusion genes (Alexandrov et al., 2009). The transformants were selected with URA3 marker and confirmed by replacement the green fluorescent signal for the red one with the same cellular distribution pattern. To re-introduce the uracil auxotrophity, the selected transformants were transformed by integrative cassette that replace the URA3 gene with the HIS3 gene. Finally, the resulting uracil-auxotrophic clones were transformed by a set of pYES2 plasmids encoding the GFP-tagged protein of interest under the control of GAL1 promoter. Before growing on GAL1 medium and imaging, the transformants were grown for ∼50 generations on the synthetic medium without uracil, containing 4 mM GuHCl to ensure the absence of any Hsp104-dependent prions.

The media used for growing cells without protein overproduction were: YPD (yeast extract 1%, peptone 2%, glucose 2%, and agar 2% when appropriate), and SC-Glu (yeast nitrogen base, 0.17 g/l; ammonium sulfate, 5 g/l; glucose, 2%; agar 2% when appropriate; a mixture of amino acids and nucleosides, appropriate for a selection of cells carrying genetic constructs with either URA3, LEU2 or HIS3 markers, or the cells with the re-gained ability to synthesize adenine). When necessary, a solid medium was prepared by adding 2% agar. For the overproduction of proteins of interest under the control of the GAL1 promoter, we used the SC-Gal medium, where glucose was replaced by 2,4% galactose. Because of some observations that accumulation of a “red pigment” (an adenine synthesis intermediate) may decrease the cell viability and even interfere with amyloidogenesis of some proteins (Nevzglyadova et al., 2011), in order to prevent its accumulation, we used SC medium with adenine concentration 100 mg/l, a 5-fold higher than standard one. To avoid cell death in the experiments with long-term incubations of cell cultures, which was shown to be provoked by medium acidification (Murakami, 2012), we buffered the SC medium pH at the value 6.5 by adding a phosphate-citrate buffer (64.2 mM Na2HPO4, 17.9 mM citric acid). For a more intense appearance of the “colony color” phenotype, a solid YPD-red medium (yeast extract (Oxoid) 0,6%, peptone 2%, glucose 2%, agar 2%) was used. The cells were grown at 30°C, and the liquid cultures were shaken at 160-200 rpm (depending on the suspension volume, and the vessel used).

### Yeast genetics

Yeast transformations with DNA plasmids or integrative cassettes was carried out according the standard LiAc-PEG method (Gietz and Schiestl, 2007).

For introducing a “tester” plasmid (overproducing either Sup35 or Hsp104 proteins) in [*PSI+*] isolates, we used a mating procedure. A MATα 74-D694 strain (preliminarily cured with 4 mM GuHCl to have [*psi-*] [*rnq-*] phenotype) was transformed with either YEplac181-SUP35, YEplac181-HSP104 or “empty” YEplac181 vector (LEU2 marker). The resulted transformants were crossed with newly-formed [*PSI+*] isolates (74-D694 strain with MATa mating type), carrying pYES2-Sup35PD-GFP plasmid (Sup35PD-GFP expression was repressed on glucose-containing medium). The mating partners were independently streaked on YPD plates, grown for 1 day, then mixed on another YPD plate with additional 1-day incubation, and then diploid cells were selected by growth on SC-Glu solid medium lacking uracil and leucine. For distinguishing [*PSI+*]/[*psi-*] states in diploids via their colony color phenotype, the adenine was added at a concentration of 6.5 mg/l, which is about 33% of its standard concentration.

To select Ade+ clones and determine their proportion in bulk cell population, cell suspensions were plated on solid medium with and without adenine in different dilutions. Typically, a 30 µl aliquot of non-diluted suspension was plated on SC-Glu medium lacking adenine (-Ade), and 30 µl aliquot of 1000-fold diluted suspension was plated on the solid YPD medium (+Ade). Cells +Ade medium were grown at 30°C for 2-3 days, and cells plated on -Ade medium were grown at 30°C for 7-10 days because of significantly decreased grow rate. The colony number for each sample and condition was counted semi-authomatically with OpenCFU software (Geissmann, 2013).

To confirm a prion status of *Ade+* clones, we probed them by growing on YPD-red medium with 4 mM GuHCl, a selective inhibitor of amyloid fragmentation activity of Hsp104. A several dozens of *Ade+* colonies were streaked on YPD-red plate, grown for 2 days at 30°C, then incubated 1 day at 4°C (for better development of color phenotype) and photographed. After that, cells were velvet-replica-plated on YPD-red plate with 4 mM GuHCl, incubated for 2 days at 30°C and then for 1 day at 4°C. Prion status was assessed by colony color switch - from white/pink on the “normal” YPD-red medium, to red on GuHCl-containing medium.

### An assessment of GFP-tagged proteins production levels with flow cytometry

To estimate and compare the levels of Sup35PD-GFP, GFP-Rnq1, Mot3PrD-GFP, Ure2PrD-GFP and Sap30PrD-GFP production under the control of the GAL1 promoter, we used GFP fluorescence level as a proxy. We have also used a production level of genomic Sup35-GFP as a proxy for a native Sup35 production level.

GFP fluorescence level (FITC channel, signal amplification = 1) and cell size (FSC channel, signal amplification = 200) were measured with flow cytometer CytoFlex S (Beckman Counter) and CytExpert Software in 10000 cells (per each sample) from stationary phase yeast culture after 18 h incubation in SC-Gal medium. FITC channel and FSC channel values for each individual cell were extracted with FlowJo software, for each cell FITC value was based on FSC value. To make a correction for the cell’s autofluorescence level in FITC channel, the same procedure was applied to the “empty” cells (i. e., cells not carrying any genes encoding GFP-tagged construct) grown under the same conditions. Using the dataset collected on 10,000 “empty” cells, we created a compensation matrix that calculates the anticipated autofluorescence value of an individual cell based on its size (i. e., FITC channel value based on FSC channel value). The values collected on the cells producing GFP-tagged chimeric constructs were corrected with this compensation matrix. The data were visualized as violin plots with BoxPlotR tool (http://shiny.chemgrid.org/boxplotr).

### An assessment of necrotic cell death rate with propidium iodide (PI)

A small aliquots of cell suspensions incubated over various time (10 µl per each sample and incubation time) were transferred to 90 µl of TBS buffer, and then PI was added for a final concentration 2 µg/ml. Cell were incubated with PI for 10 minutes at dark, and then 10000 cells per sample were subjected to flow cytometry analysis. The data were acquired and analyzed with flow cytometer CytoFlex S (Beckman Counter) and CytExpert Software. PI signal was detected in PE channel (signal amplification = 245). For setting a threshold for dead (PI-positive) cells quantification, we boiled the suspension of the same cells for 5 min, and then subjected them for the same procedure.

### Fluorescence microscopy-based experiments

Images and videos of the yeast cells were obtained using an Nikon Ti2 inverted microscope and a Photometric CoolSnap HQ 12-bit CCD camera. For some experiments we also used Axioscop 40 fluorescence microscope (Zeiss, Oberkochen, Germany). Z stacks of several fields of view were collected for each sample. To immobilize the live cells for imaging, we have used the agar pad technique (Alexandrov and Dergalev, 2019). Representative cells or groups of cells were selected for figures. For the quantitative experiments, typically 200-400 cells (or individual proteinaceous particles, if applicable) were counted per each replicate.

For a life cell imaging in a “single cell” mode (i.e., time-lapse imaging of the exact cell or the group of cells subjected to any type of chemical treatment) we have used a glass-bottom Petri dishes with the cells immobilized on the glass with concanavalin A.

To perform a super-resolution structural illumination microscopy (SIM), we fixed cells with 2.4% formaldehyde for 15 min, embedded them into Moviol resin, incubated for 24 hrs at +4°C and then for 24 hrs at room temperature, and then examined using an N-SIM microscopic system (Nikon Instech Co., Tokyo, Japan) with an immersion objective 100×/1.49 NA, excitation laser wavelength of 488 nm. Image stacks (with a *z*-axis step of 0.12 μm) were acquired with an EMCCD camera (iXon 897, Andor, effective pixel size 60 nm) in the 3D-SIM mode. Serial optical sections of the same cell, taken in the wide field mode, were processed using the AutoQuant blind deconvolution algorithm. Image acquisition and SIM reconstruction were performed using the NIS-Elements 4.2 software (Nikon Instech Co., Tokyo, Japan).

Determination of mean GFP intensity per cell, fluorescent particles analysis (shape, size, number, particle/cytosol intensity ratio) was performed with either FIJI or NIS-Elements software, using image segmentation and particle analysis tools.

### SDD-AGE analysis of protein polymers

To detect the formation of the amyloid polymers, the cells collected at different time points (∼10 ml of stationary yeast culture suspension per each sample) were lysed with glass bead in 100 μl TBS buffer supplemented with 5 mM PMSF, 1 mM DTT and cOmplete protease inhibitor cocktail (Roche). The lysis was performed for 15 min at 4°C. Cell lysates were mixed with SDD-AGE sample buffer and, where indicated, sonicated for 30 sec (50% amplitude, 5/5 sec pulse/relaxation) to crush the super-aggregates resistant to 2% SDS, and generate a smaller-sized amyloid fibrils able to enter a 1,8% agarose gel. The SDD-AGE procedure was performed as follows (Kryndushkin et al., 2003). The protein samples resolved in agarose gel were vacuum-transferred onto the nitrocellulose membrain (Porablot, MACHEREY-NAGEL), and subjected to a standard Western-Blotting procedure with either anti-Sup35NM, anti-Rnq1 or anti-GFP rabbit polyclonal antibodies. The integrated intensity of the protein-of-interest amyloid shmeres, as a measure of the amyloid amount, was then quantified using FIJI software.

### Determination of soluble/amyloid Sup35PD-GFP proportion in cell lysates (“boiled gels” assay)

To investigate, which proportion of Sup35PD-GFP (or Sup35PD(ΔR)-GFP) molecules was converted to amyloid in the cells incubated for up to 7 days, we employed a modified SDS-PAGE (5% acrylamide in the concentration gel, 10% acrylamide in the resolving gel) followed by Western Blotting. The cell lysates prepared on TBS buffer with protease inhibitors cocktail (cOmplete, Pierce), 5 mM PMSF and 1 mM DTT were briefly sonicated (10 s, 50% amplitude), mixed with SDS-PAGE sample buffer, cell debris was spun down at 2348 g 1 min, and the supernatant was loaded on an SDS-PAGE gel. The electrophoresis was run under 110 V of current for about 40 min. Importantly, at this stage only monomeric form of Sup35PD-GFP can run inside the resolving gel, while SDS-insoluble amyloid polymers are stacked in the wells or in the upper part of the concentration gel. After this, we loaded a 5 μl of SDS-PAGE sample buffer to the sample wells, boiled the gel for about 5 min in the water bath, cooled it, and then additionally run electrophoresis for about 40 min at 110 V. At this stage, amyloid fibrils dissolve and released Sup35PD-GFP monomeric molecules which can enter the gel. This portion of Sup35PD-GFP molecules then runs in the same lane with Sup35PD-GFP monomers, but as a distinct protein band (with a shorter run distance) - and, therefore, can be quantified separately from non-amyloid form. This procedure was followed by standard Western Blotting with Sup35NM primary antibodies, the band intensities were quantified with FIJI.

### PrK resistance analysis of Sup35PD-GFP amyloid fibrils

For mapping the densely-packed regions (“amyloid cores”) of Sup35PD-GFP fibrils, we extracted and purified the detergent-resistant fractions from a large amount of yeast cell pellets (typically 3-5 grams per preparation), as described in the original work (Dergalev et al, 2019). In brief, yeasts were glass beads-beaten, the heavy fraction of lysate was mixed with sarkosyl (up to the final concentration = 5%), sonicated, loaded on 20-65% sucrose gradient and ultracentrifuged at 268000 g for 4 hours. The fraction with bright GFP fluorescence was collected, and the obtained Sup35PD-GFP sample (diluted to the final concentration 200 μg/ml) was digested with PrK (25 μg/ml) for 1 hour at RT. After that, the PrK-resistant part of fibrils was preciped with 40% acetone, resuspended in bidistilled water, boiled (which causes the fibrils disassembly to the monomeric peptides) and analyzed with MALDI-TOF/TOF mass spectrometer UltrafleXtreme (Bruker, Germany). Importantly, in our reaction conditions, PrK has no sequence specificity and digests all parts of Sup35PD molecules except ones folded in densely packed cross-β structures, and, partially, the β-barrel of GFP molecule (the latter was not taken into account when analyzing the raw MALDI data). Peptides were identified by tandem mass spectroscopy (MS-MS) and/or as groups of related peaks. The raw MALDI data were analyzed with FlexAnalysis software, and PrK resistance charts were calculated and drawn as a function of the normalized PrK resistance index for a given sequence position. The contribution of each peptide in the PrK resistance map of a given preparation was proportional to its MS peak area.

### The “cell burst” test

The 50 μl aliquots of stationary yeast culture at different stages of incubation were spun down, the cells were washed twice with bidistilled water, resuspended in 100 μl of 80 mM citrate-phosphate buffer (pH = 6.5) containing 0.8 M sorbitol and 0.04 mg/ml lyticase (Sigma), and incubated 60 min at 30°C with gentle agitation. After that, a small aliquot (5 μl) of spheroplasts in the sorbitol-containing buffer was loaded on a 2% agar pad prepared on the citrate-phosphate buffer with 0.3 M sorbitol. The aliquot was immediately covered by a cover glass and subjected to widefield fluorescence microscopy. As the sorbitol diffuses from the aliquot into the pad, the gradual drop in its concentration creates hypo-osmotic conditions for yeast spheroplasts, leading them to hyper-hydration and subsequent bursting, releasing the proteinaceous particles into the extracellular space. After that, the liquid-like Sup35PD-GFP droplets dissolve quickly (∼5 sec) because of the sudden drop of Sup35PD-GFP concentration in the droplet’s exterior, while Sup35PD-GFP hydrogels remain stable in extracellular spase over more than several minutes.

### Thermal stability assay

Amyloid samples isolated as described above (for PrK resistance analysis of Sup35PD-GFP amyloid polymers) were mixed with SDD-AGE sample buffer (total volume = 30), transferred to PCR tubes, and covered with vaseline oil to prevent evaporation. The samples were then heated in a Tercik (DNK-Technologii) multi-channel DNA amplifier for 8 minutes at the appropriate temperature (either 50, 60, 70, 80, or 95°C) and then cooled down to 10°C. Cooled samples were then loaded on 1.8% agarose gel and analyzed by SDD-AGE followed by Western blotting (as described above) with anti-Sup35PD rabbit polyclonal antibodies. The integrated intensity of Sup35PD-GFP shmeres, as a measure of the amount of Sup35PD-GFP amyloids that withstand the heating, was then quantified using Fiji software. For each preparation of amyloid, the experiment was performed in 3 technical replicates.

### Detection of [RNQ+], [MOT+] and [URE+] prions formation via LSPT pathway

A wide-spread approach to detecting prions in yeast cells is to adapt the ADE1 or ADE2 gene-based detection system initially elaborated for [*PSI+*] prion. This implies a fusion of respective protein PD with the Sup35 C-terminal domain (the translation termination subunit II), replacement of episomal SUP35 gene with such chimeric construct expressed from a centromeric plasmid in the strain with a nonsense mutation in either ADE1 or ADE2 gene (like *ade1-14* or *ade2-1*), and detection of Ade+ clones with white/pink coloration and ability to grow on the medium lacking adenine. However, the detection of prion state with such Sup35C-tagged chimera might be not appropriate for Rnq1 protein, because its putative amyloidogenic segments are located predominantly in the C-terminal part of the protein, and its PrK-resistant amyloid core was also reported to be C-terminal (Dergalev et al., 2019). Therefore, the C-terminal fusion of Rnq1 with Sup35C, a nearly 55 kD polypeptide, may alter its amyloid and prion behavior - while Sup35C, fused with Rnq1 from its own C-terminus, could likely be not fully functional in the translation termination catalysis.

For Mot3 and Ure2 prions, we initially aimed to construct a detection system as described above. Starting from centromeric pRS315 plasmid with SUP35 gene under its endogenous promoter, we replaced Sup35PD for respective PD of either Ure2 or Mot3 protein. The resulting plasmids were shuffled with pRS316-SUP35 plasmid in the *ade1-14* strain, where the endogenous SUP35 gene was deleted (SUP35::TRP1). However, for both Mot3PD and Ure2PD, the strains obtained in this way have spontaneously switched in the Ade+ phenotype (white/pink colony color) at a high rate (∼50%), which made them inappropriate for the detection of [*PRION+*] clones formed via LLPS and LSPT.

Therefore, we needed an alternative approach to detect the prion switch of our proteins of interest (POI’s). Eventually, we developed a novel, indirect detection system relied on the nearly-universal ability of one amyloid-based yeast prion to cross-seed amyloid polymerization of another yeast prionogenic protein when the latter is overproduced (Du and Li, 2014; Arslan et al., 2015; Du et al., 2015). Salnikova and coauthors have shown that when SUP35 in moderately overexpressed (as a multicopy gene under the control of native promoter) in cells carrying [*RNQ+*] prion, it forms the non-heritable amyloids - which, however, have switched into the unstable [*PSI+*] prion-like state when cells were plated on the medium lacking adenine (observed as colonies with unstable Ade+ phenotype; Salnikova et al., 2005). As Sup35 was expressed at a level significantly lower than one achievable with GAL1 promoter (and required to form the phase-separated assemblies), the cross-seeding by either [*RNQ+*] or any other amyloid-based prion (including [*MOT+*] and [*URE+*]) is likely the only way for such Ade+ phenotype to emerge with observable frequency. Sap30 was leaved apart this experiment, because Sap30PD-GFP was reported to form prion-like polymers that can not transfer their prion state onto the wild-type endogenous Sap30 (Du et al., 2015).

The experiments were performed as follows. 74-D694 ade1-14 [psi-] [rnq-] strain was transformed with two plasmids simultaneously - one, YEplac181-SUP35 (multicopy vector, SUP35 promoter), and second, pYES2 encoding the POI (either GFP-Rnq1, Ure2PD-GFP, or Mot3PD-GFP) under the control of GAL1 promoter. For the [RNQ+] induction experiment, we used [rnq-] strain with intact RNQ1 gene; for the [MOT+] and [URE+] induction experiment, we used Δrnq1 strain to prevent possible co-appearance of [RNQ+] prion together with either [MOT+] or [URE+] (the similar phenomenon was described for [PSI+] and [RNQ+] co-occurrence Derkatch, 2000). The transformants were grown for ∼50 generations on the selective SC-Glu plate with 4 mM GuHCl, to ensure that any amyloid-base prion is not present in the cells at the start of experiment. Then the cells were grown for 24h to stationary culture in the liquid SC-Glu medium, diluted 4-fold by the selective SC-Gal medium with pH buffered at 6.5, and then incubated for up to 7 days. After 1, 3, and 7 days of incubation, 1 μl of each suspension was inoculated to the 1000 μl aliquot of selective SC-Glu medium (repression of POI overproduction), and cells were grown for about two days up to stationary phase state (∼10 generations). At this stage, GAL1-mediated overproduction of the protein of interest was shut down, the cells with solid-like aggregates of the POI were about 1000-fold diluted, and such aggregates themselves are expected to be disassembled by chaperones, as was demonstrated for the macroscale amyloid aggregates formed by overproduced Sup35PD-GFP (Kumar et al., 2016). Thus, only the amyloid forms of wild-type endogenous POI (either Rnq1, Ure2, or Mot3), that were stably inherited for 10 generations without overproduction of its GFP-tagged PD (i.e., an established prion), would eventually be scored. The glucose medium-grown cells were probed for *Ade+* clones formation by plating them on SC-Glu medium without adenine (30 μl of non-diluted suspension) and on YPD medium containing adenine (30 μl of 1500-fold diluted suspension). *Ade+* clones were tested for their [*PRION+*] status by growing them on YPD-red plate containing 4 mM GuHCl.

To ensure that GuHCl-curable *Ade+* clones, selected after GFP-Rnq1 and Mot3PD-GFP overproduction, are indeed positive for [RNQ+] and [MOT+] priones (respectively), we subcloned them on YPD plates, selected *ade-leu-* subclones (i.e., dark-red clones that have lost YEplac181-SUP35 plasmid and do not contain prion-like form of Sup35), grew them in 100 μl of selective SC-Glu medium and then induced repeatative overproduction of GFP-tagged POI by adding 300 μl of SC-Gal medium. As a negative control, we overproduced the corresponding constructs in the naïve cells. After 4h incubation with galactose, cells were stained with proteostat (as described above) and subjected to fluorescence microscopy. In the naïve cells, POI-GFP was either distributed diffusely or separated between diffuse state in the cytosol and bright spherical bodies, which were not stained with proteostat (i.e., represent the condensates). In *Ade+* clones, nearly all POI-GFP was assembled in large non-spherical bodies stained with proteostat (i.e., containing amyloids). The latter pattern is typical for yeast cells with endogenous POI in the prionic form and overproduced POI-GFP (Kumar et al., 2016).

## Acknowledgments

The authors thank the Shared-Access Equipment Centre “Industrial Biotechnology” of FRC “Fundamentals of Biotechnology” (RAS) and Nikon Center of Excellence at A.N. Belozersky Institute of Physical and Chemical Biology for providing research infrastructure.

## Funding Statement

This research was funded by the Russian Science Foundation, grant #23-74-00062.

## Author Contributions

Conceptualization: AAD; methodology: AAD, IIK, VVK, AIA; acquisition of data: MKA, AAD, SBN, IIK, SAG, PAT, LVI; analysis of data: AAD, MKA, SBN, LVI, VVK; writing — original draft: AAD; writing — review and editing: AAD, AIA; visualization: AAD, MKA; supervision: AAD; funding acquisition: VVK. All authors have read and agreed to the published version of the manuscript.

## Notes

### Competing Interest Statement

The authors have declared no competing interest.

