## Supplementary material for "A liquid-to-solid phase transition of biomolecular condensates drives *in vivo* formation of yeast amyloids and prions": Figure S1

### Slide 1
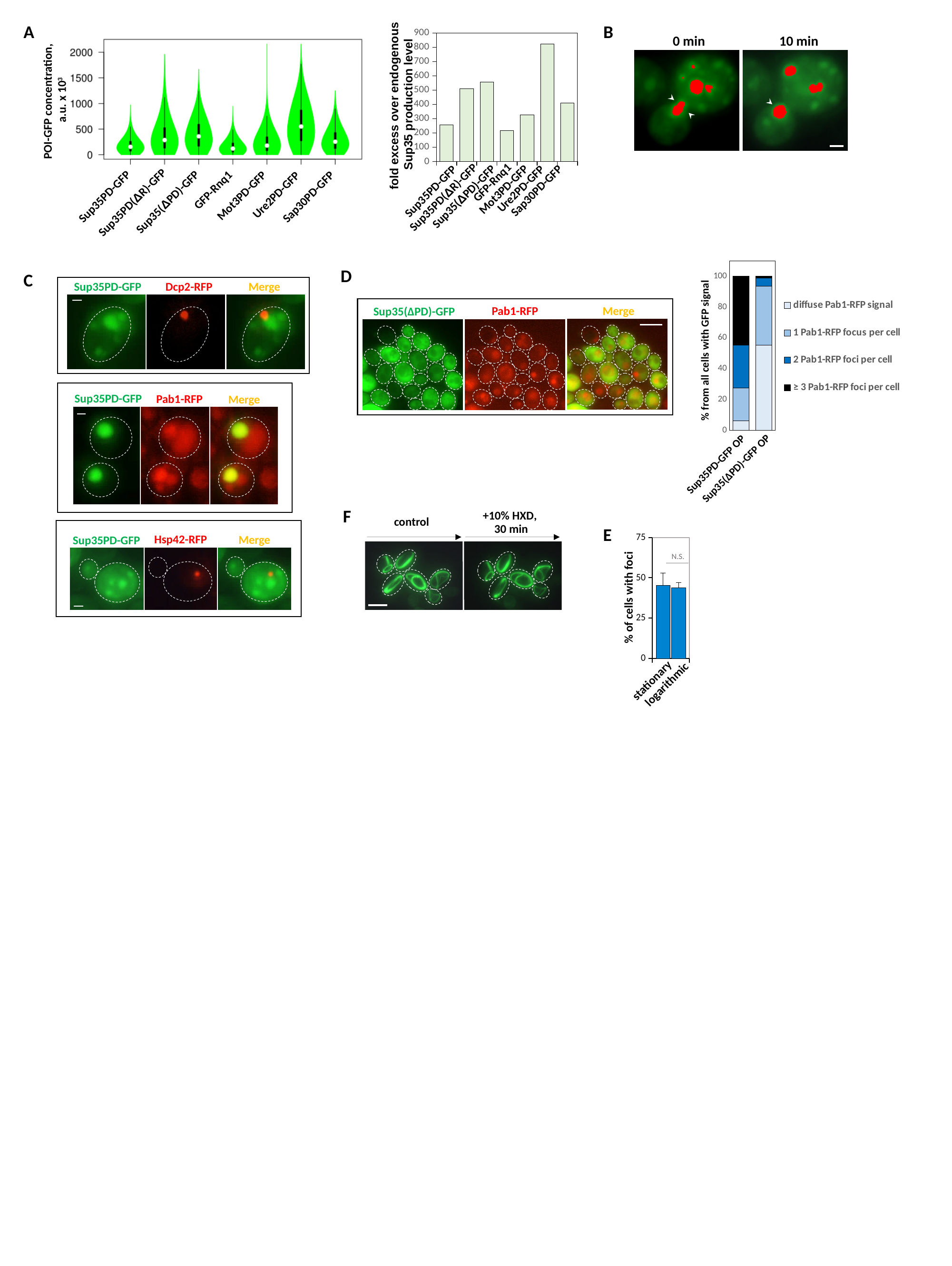

#### Chart
| Category | |
|---|---|
| NMG | 254.6396425098936 |
| R2 | 508.56981019752965 |
| -152 | 556.6351689748262 |
| Rnq | 217.021470035471 |
| Mot | 327.5328198339403 |
| Ure | 822.4363270042471 |
| Sap | 409.4050836976114 |fold excess over endogenous
Sup35 production level
GFP-Rnq1
Sap30PD-GFP
Mot3PD-GFP
Sup35PD-GFP
Ure2PD-GFP
Sup35(ΔPD)-GFP
Sup35PD(ΔR)-GFP
A
GFP-Rnq1
Sap30PD-GFP
Mot3PD-GFP
Sup35PD-GFP
Ure2PD-GFP
Sup35(ΔPD)-GFP
Sup35PD(ΔR)-GFP
POI-GFP concentration,
a.u. x 10^3^
B
0 min 10 min
#### Chart
| Category |
|---|
| 1 | 6.25 | 21.153846153846153 | 27.884615384615387 | 44.71153846153847 |% from all cells with GFP signal
Sup35PD-GFP OP
Sup35(ΔPD)-GFP OP
D
Pab1-RFP
Merge
Sup35(ΔPD)-GFP
C
Dcp2-RFP
Merge
Sup35PD-GFP
Sup35PD-GFP
Pab1-RFP
Merge
Hsp42-RFP
Merge
Sup35PD-GFP
F
+10% HXD,
30 min
control
#### Chart
| Category | Stationary | Log |
|---|---|---|N.S.
% of cells with foci
stationary
logarithmic
E

### Slide 2
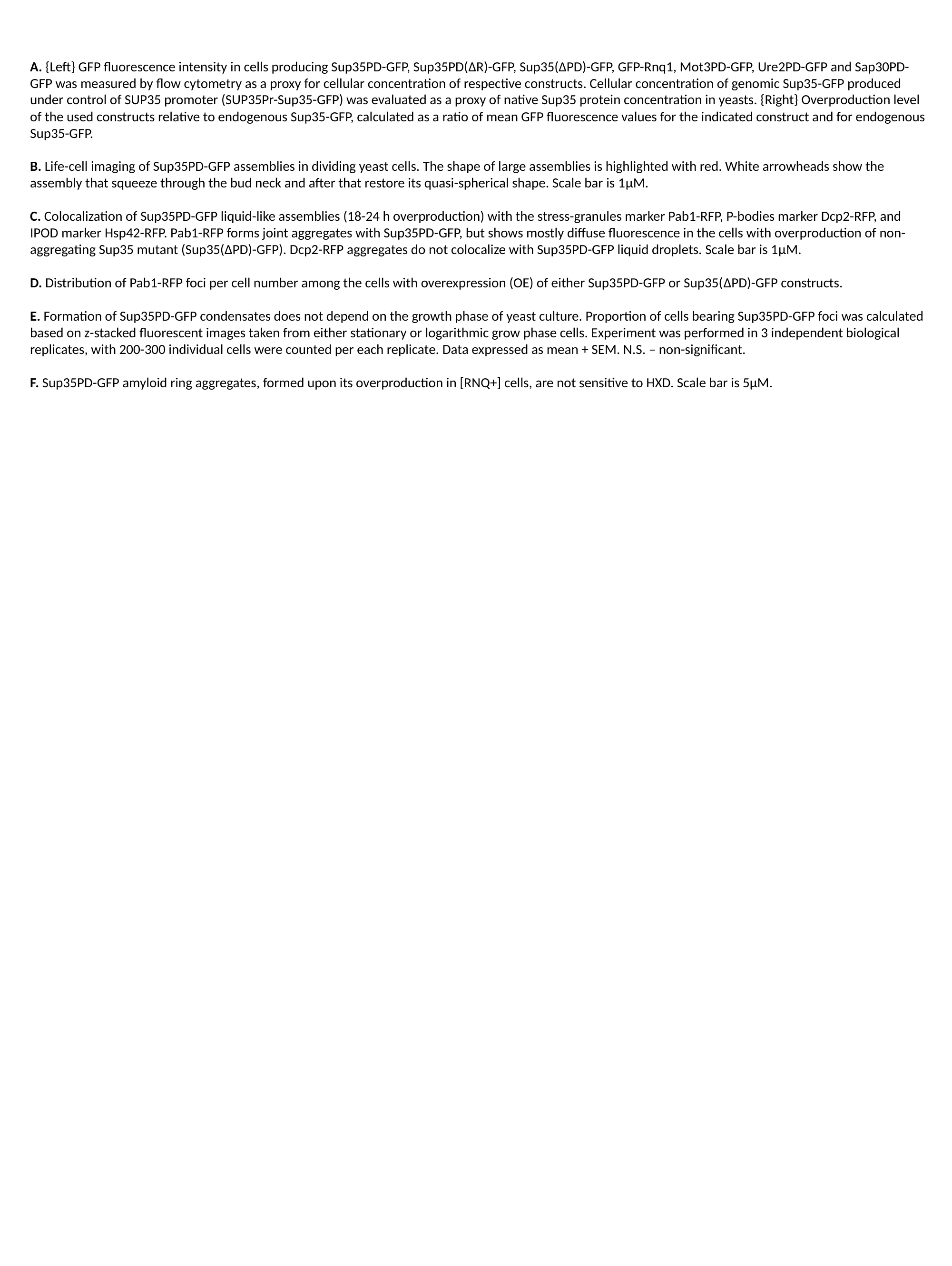

A. {Left} GFP fluorescence intensity in cells producing Sup35PD-GFP, Sup35PD(ΔR)-GFP, Sup35(ΔPD)-GFP, GFP-Rnq1, Mot3PD-GFP, Ure2PD-GFP and Sap30PD-GFP was measured by flow cytometry as a proxy for cellular concentration of respective constructs. Cellular concentration of genomic Sup35-GFP produced under control of SUP35 promoter (SUP35Pr-Sup35-GFP) was evaluated as a proxy of native Sup35 protein concentration in yeasts. {Right} Overproduction level of the used constructs relative to endogenous Sup35-GFP, calculated as a ratio of mean GFP fluorescence values for the indicated construct and for endogenous Sup35-GFP.
B. Life-cell imaging of Sup35PD-GFP assemblies in dividing yeast cells. The shape of large assemblies is highlighted with red. White arrowheads show the assembly that squeeze through the bud neck and after that restore its quasi-spherical shape. Scale bar is 1μM.
C. Colocalization of Sup35PD-GFP liquid-like assemblies (18-24 h overproduction) with the stress-granules marker Pab1-RFP, P-bodies marker Dcp2-RFP, and IPOD marker Hsp42-RFP. Pab1-RFP forms joint aggregates with Sup35PD-GFP, but shows mostly diffuse fluorescence in the cells with overproduction of non-aggregating Sup35 mutant (Sup35(ΔPD)-GFP). Dcp2-RFP aggregates do not colocalize with Sup35PD-GFP liquid droplets. Scale bar is 1μM.
D. Distribution of Pab1-RFP foci per cell number among the cells with overexpression (OE) of either Sup35PD-GFP or Sup35(ΔPD)-GFP constructs.
E. Formation of Sup35PD-GFP condensates does not depend on the growth phase of yeast culture. Proportion of cells bearing Sup35PD-GFP foci was calculated based on z-stacked fluorescent images taken from either stationary or logarithmic grow phase cells. Experiment was performed in 3 independent biological replicates, with 200-300 individual cells were counted per each replicate. Data expressed as mean + SEM. N.S. – non-significant.
F. Sup35PD-GFP amyloid ring aggregates, formed upon its overproduction in [RNQ+] cells, are not sensitive to HXD. Scale bar is 5μM.
