## Supplementary material for "A liquid-to-solid phase transition of biomolecular condensates drives *in vivo* formation of yeast amyloids and prions": Figure S2

### Slide 1
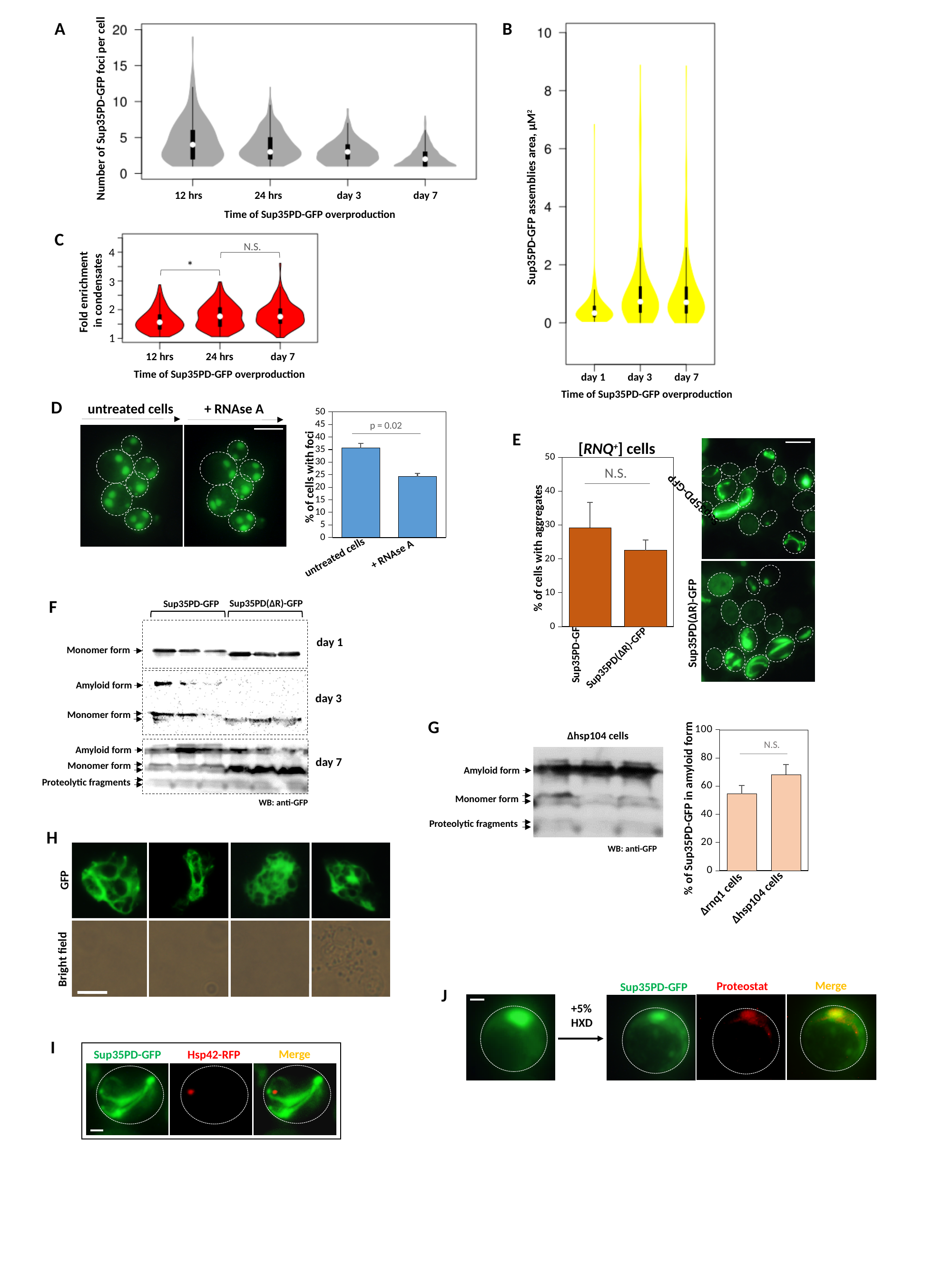

Sup35PD-GFP assemblies area, μM^2^
day 1 day 3 day 7
B
Time of Sup35PD-GFP overproduction
A
Number of Sup35PD-GFP foci per cell
12 hrs 24 hrs day 3 day 7
Time of Sup35PD-GFP overproduction
Fold enrichment
in condensates
12 hrs 24 hrs day 7
Time of Sup35PD-GFP overproduction
N.S.
4
*
3
2
1
C
D
untreated cells + RNAse A
#### Chart
| Category | |
|---|---|
| contr | 35.56815539764884 |
| rnase | 24.2236245036869 |% of cells with foci
+ RNAse A
untreated cells
p = 0.02
E
[RNQ^+^] cells
#### Chart
| Category | |
|---|---|
| NMG | 29.196261194402457 |
| R2 | 22.518846300912784 |N.S.
% of cells with aggregates
Sup35PD-GFP
Sup35PD(ΔR)-GFP
Sup35PD-GFP
Sup35PD(ΔR)-GFP
F
Sup35PD(ΔR)-GFP
Sup35PD-GFP
day 1
Monomer form
Amyloid form
day 3
Monomer form
Amyloid form
day 7
Monomer form
Proteolytic fragments
WB: anti-GFP
#### Chart
| Category | |
|---|---|% of Sup35PD-GFP in amyloid form
Δrnq1 cells
Δhsp104 cells
N.S.
G
Δhsp104 cells
Amyloid form
Monomer form
Proteolytic fragments
WB: anti-GFP
H
GFP
Bright field
Merge
Proteostat
Sup35PD-GFP
+5%
HXD
J
I
Merge
Sup35PD-GFP
Hsp42-RFP

### Slide 2
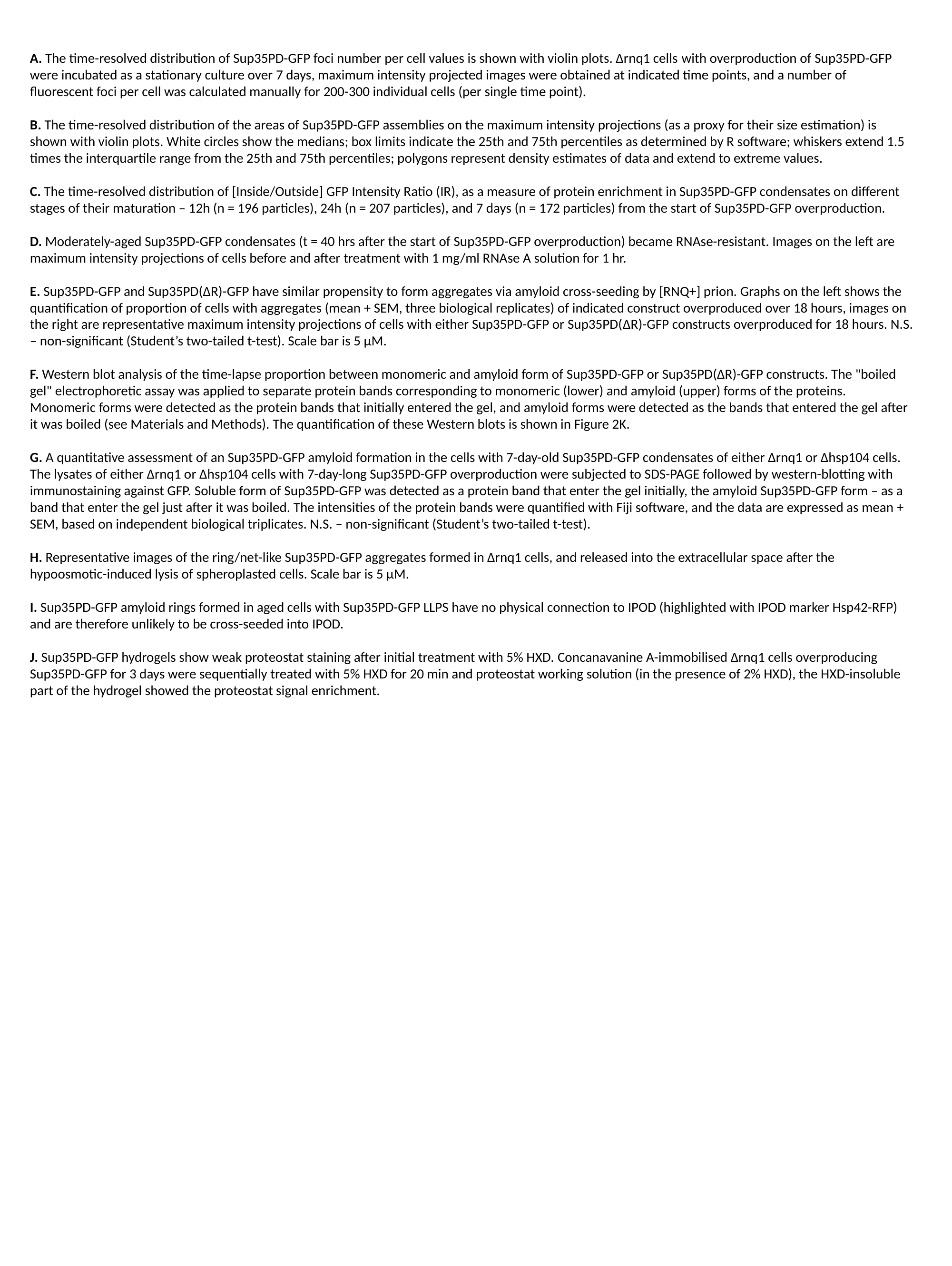

A. The time-resolved distribution of Sup35PD-GFP foci number per cell values is shown with violin plots. Δrnq1 cells with overproduction of Sup35PD-GFP were incubated as a stationary culture over 7 days, maximum intensity projected images were obtained at indicated time points, and a number of fluorescent foci per cell was calculated manually for 200-300 individual cells (per single time point).
B. The time-resolved distribution of the areas of Sup35PD-GFP assemblies on the maximum intensity projections (as a proxy for their size estimation) is shown with violin plots. White circles show the medians; box limits indicate the 25th and 75th percentiles as determined by R software; whiskers extend 1.5 times the interquartile range from the 25th and 75th percentiles; polygons represent density estimates of data and extend to extreme values.
C. The time-resolved distribution of [Inside/Outside] GFP Intensity Ratio (IR), as a measure of protein enrichment in Sup35PD-GFP condensates on different stages of their maturation – 12h (n = 196 particles), 24h (n = 207 particles), and 7 days (n = 172 particles) from the start of Sup35PD-GFP overproduction.
D. Moderately-aged Sup35PD-GFP condensates (t = 40 hrs after the start of Sup35PD-GFP overproduction) became RNAse-resistant. Images on the left are maximum intensity projections of cells before and after treatment with 1 mg/ml RNAse A solution for 1 hr.
E. Sup35PD-GFP and Sup35PD(ΔR)-GFP have similar propensity to form aggregates via amyloid cross-seeding by [RNQ+] prion. Graphs on the left shows the quantification of proportion of cells with aggregates (mean + SEM, three biological replicates) of indicated construct overproduced over 18 hours, images on the right are representative maximum intensity projections of cells with either Sup35PD-GFP or Sup35PD(ΔR)-GFP constructs overproduced for 18 hours. N.S. – non-significant (Student’s two-tailed t-test). Scale bar is 5 μM.
F. Western blot analysis of the time-lapse proportion between monomeric and amyloid form of Sup35PD-GFP or Sup35PD(ΔR)-GFP constructs. The "boiled gel" electrophoretic assay was applied to separate protein bands corresponding to monomeric (lower) and amyloid (upper) forms of the proteins. Monomeric forms were detected as the protein bands that initially entered the gel, and amyloid forms were detected as the bands that entered the gel after it was boiled (see Materials and Methods). The quantification of these Western blots is shown in Figure 2K.
G. A quantitative assessment of an Sup35PD-GFP amyloid formation in the cells with 7-day-old Sup35PD-GFP condensates of either Δrnq1 or Δhsp104 cells. The lysates of either Δrnq1 or Δhsp104 cells with 7-day-long Sup35PD-GFP overproduction were subjected to SDS-PAGE followed by western-blotting with immunostaining against GFP. Soluble form of Sup35PD-GFP was detected as a protein band that enter the gel initially, the amyloid Sup35PD-GFP form – as a band that enter the gel just after it was boiled. The intensities of the protein bands were quantified with Fiji software, and the data are expressed as mean + SEM, based on independent biological triplicates. N.S. – non-significant (Student’s two-tailed t-test).
H. Representative images of the ring/net-like Sup35PD-GFP aggregates formed in Δrnq1 cells, and released into the extracellular space after the hypoosmotic-induced lysis of spheroplasted cells. Scale bar is 5 μM.
I. Sup35PD-GFP amyloid rings formed in aged cells with Sup35PD-GFP LLPS have no physical connection to IPOD (highlighted with IPOD marker Hsp42-RFP) and are therefore unlikely to be cross-seeded into IPOD.
J. Sup35PD-GFP hydrogels show weak proteostat staining after initial treatment with 5% HXD. Concanavanine A-immobilised Δrnq1 cells overproducing Sup35PD-GFP for 3 days were sequentially treated with 5% HXD for 20 min and proteostat working solution (in the presence of 2% HXD), the HXD-insoluble part of the hydrogel showed the proteostat signal enrichment.
