## Supplementary material for "A liquid-to-solid phase transition of biomolecular condensates drives *in vivo* formation of yeast amyloids and prions": Figure S4

### Slide 1
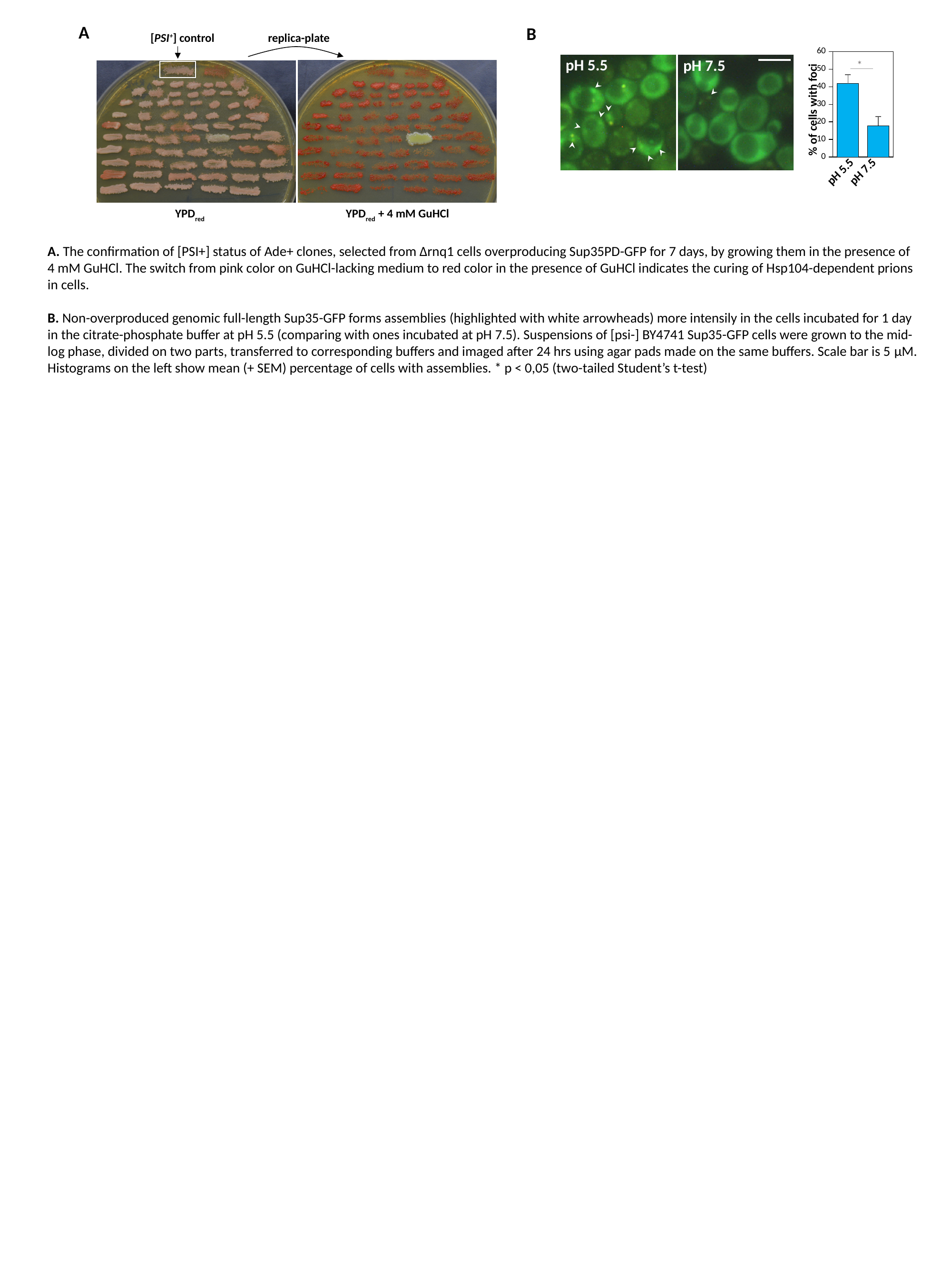

A
B
#### Chart
| Category | |
|---|---|% of cells with foci
pH 7.5
pH 5.5
*
pH 5.5
pH 7.5
replica-plate
[PSI^+^] control
YPD_red_
YPD_red_ + 4 mM GuHCl
A. The confirmation of [PSI+] status of Ade+ clones, selected from Δrnq1 cells overproducing Sup35PD-GFP for 7 days, by growing them in the presence of 4 mM GuHCl. The switch from pink color on GuHCl-lacking medium to red color in the presence of GuHCl indicates the curing of Hsp104-dependent prions in cells.
B. Non-overproduced genomic full-length Sup35-GFP forms assemblies (highlighted with white arrowheads) more intensily in the cells incubated for 1 day in the citrate-phosphate buffer at pH 5.5 (comparing with ones incubated at pH 7.5). Suspensions of [psi-] BY4741 Sup35-GFP cells were grown to the mid-log phase, divided on two parts, transferred to corresponding buffers and imaged after 24 hrs using agar pads made on the same buffers. Scale bar is 5 μM. Histograms on the left show mean (+ SEM) percentage of cells with assemblies. * p < 0,05 (two-tailed Student’s t-test)
