## Supplementary material for "A liquid-to-solid phase transition of biomolecular condensates drives *in vivo* formation of yeast amyloids and prions": Figure S5

### Slide 1
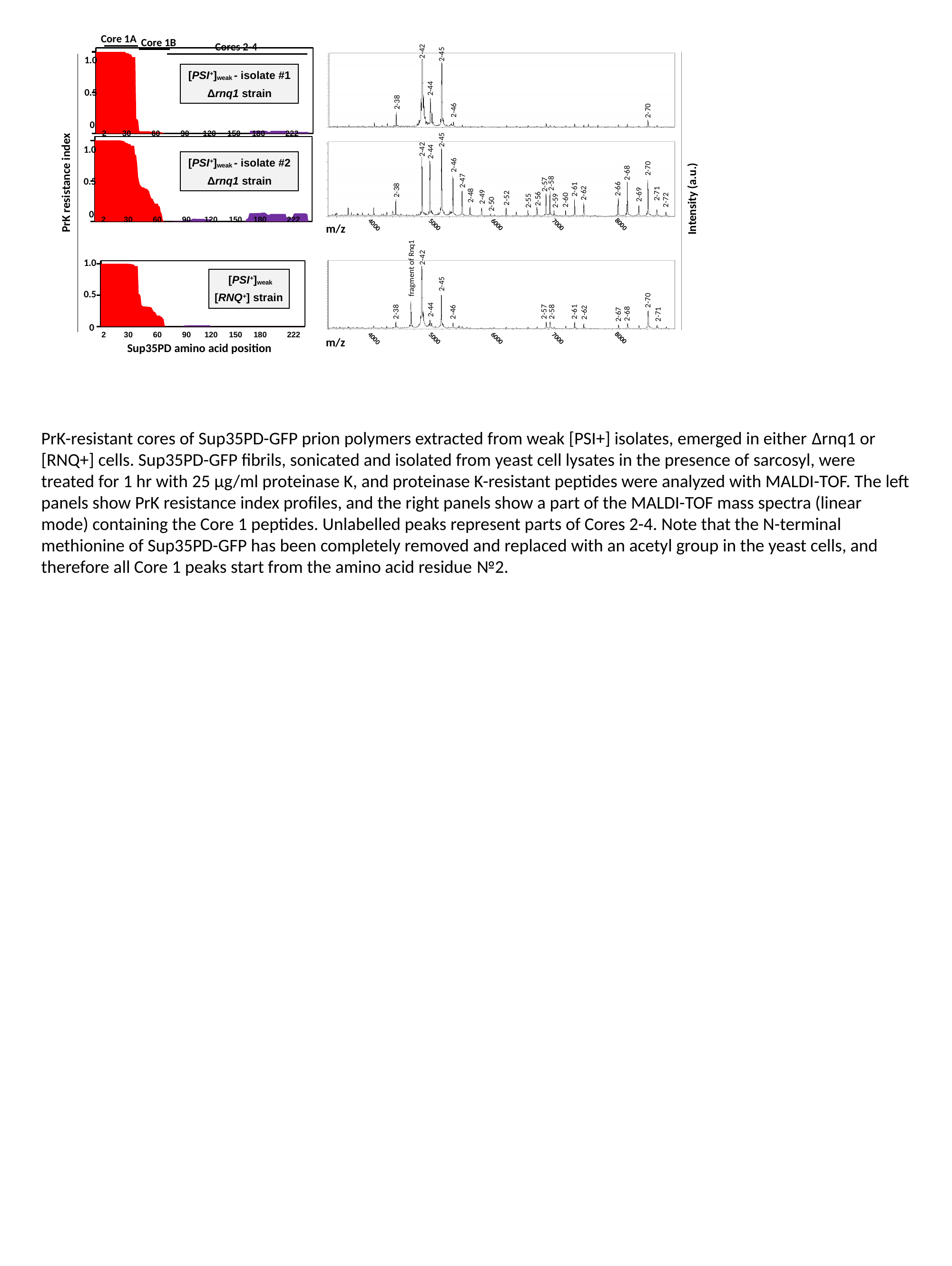

Core 1A
Core 1B
Cores 2-4
#### Chart
| Category | | |
|---|---|---|
| 1 | 0.9999998746690246 | 0.0 |
| 2 | 0.9999998746690246 | 0.0 |
| 3 | 0.9999998746690246 | 0.0 |
| 4 | 0.9999998746690246 | 0.0 |
| 5 | 0.9999998746690246 | 0.0 |
| 6 | 0.9999998746690246 | 0.0 |
| 7 | 0.9999998746690246 | 0.0 |
| 8 | 0.9999998746690246 | 0.0 |
| 9 | 0.9999998746690246 | 0.0 |
| 10 | 0.9999998746690246 | 0.0 |
| 11 | 0.9999998746690246 | 0.0 |
| 12 | 0.9999998746690246 | 0.0 |
| 13 | 0.9999998746690246 | 0.0 |
| 14 | 0.9999998746690246 | 0.0 |
| 15 | 0.9999998746690246 | 0.0 |
| 16 | 0.9999998746690246 | 0.0 |
| 17 | 0.9999998746690246 | 0.0 |
| 18 | 0.9999998746690246 | 0.0 |
| 19 | 0.9999998746690246 | 0.0 |
| 20 | 0.9999998746690246 | 0.0 |
| 21 | 0.9999998746690246 | 0.0 |
| 22 | 0.9999998746690246 | 0.0 |
| 23 | 0.9999998746690246 | 0.0 |
| 24 | 0.9999998746690246 | 0.0 |
| 25 | 0.9999998746690246 | 0.0 |
| 26 | 0.9999998746690246 | 0.0 |
| 27 | 0.9999998746690246 | 0.0 |
| 28 | 0.9999998746690246 | 0.0 |
| 29 | 0.9999998746690246 | 0.0 |
| 30 | 0.9999998746690246 | 0.0 |
| 31 | 0.9950735788709211 | 0.0 |
| 32 | 0.9832314454479659 | 0.0 |
| 33 | 0.9832314454479659 | 0.0 |
| 34 | 0.9832314454479659 | 0.0 |
| 35 | 0.9704500263418646 | 0.0 |
| 36 | 0.9704500263418646 | 0.0 |
| 37 | 0.965357274853077 | 0.0 |
| 38 | 0.9349077731500768 | 0.0 |
| 39 | 0.9349077731500768 | 0.0 |
| 40 | 0.9349077731500768 | 0.0 |
| 41 | 0.9349077731500768 | 0.0 |
| 42 | 0.17612347039729606 | 0.0 |
| 43 | 0.17612347039729606 | 0.0 |
| 44 | 0.15309709985266096 | 0.0 |
| 45 | 0.03201247457216332 | 0.0 |
| 46 | 0.026186905252159554 | 0.0 |
| 47 | 0.026186905252159554 | 0.0 |
| 48 | 0.026186905252159554 | 0.0 |
| 49 | 0.026186905252159554 | 0.0 |
| 50 | 0.026186905252159554 | 0.0 |
| 51 | 0.026186905252159554 | 0.0 |
| 52 | 0.026186905252159554 | 0.0 |
| 53 | 0.026186905252159554 | 0.0 |
| 54 | 0.026186905252159554 | 0.0 |
| 55 | 0.026186905252159554 | 0.0 |
| 56 | 0.026186905252159554 | 0.0 |
| 57 | 0.021050335023796832 | 0.0 |
| 58 | 0.021050335023796832 | 0.0 |
| 59 | 0.021050335023796832 | 0.0 |
| 60 | 0.021050335023796832 | 0.0 |
| 61 | 0.016210546164990996 | 0.0 |
| 62 | 0.016210546164990996 | 0.0 |
| 63 | 0.016210546164990996 | 0.0 |
| 64 | 0.016210546164990996 | 0.0 |
| 65 | 0.016210546164990996 | 0.0 |
| 66 | 0.016210546164990996 | 0.0 |
| 67 | 0.016210546164990996 | 0.0 |
| 68 | 0.011714762875025734 | 0.0 |
| 69 | 0.011714762875025734 | 0.0 |
| 70 | 0.0 | 0.0 |
| 71 | 0.0 | 0.0 |
| 72 | 0.0 | 0.0 |
| 73 | 0.0 | 0.0 |
| 74 | 0.0 | 0.0 |
| 75 | 0.0 | 0.0 |
| 76 | 0.0 | 0.0 |
| 77 | 0.0 | 0.0 |
| 78 | 0.0 | 0.0 |
| 79 | 0.0 | 0.0 |
| 80 | 0.0 | 0.0 |
| 81 | 0.0 | 0.0 |
| 82 | 0.0 | 0.0 |
| 83 | 0.0 | 0.0 |
| 84 | 0.0 | 0.0 |
| 85 | 0.0 | 0.0 |
| 86 | 0.0 | 0.0 |
| 87 | 0.0 | 0.0 |
| 88 | 0.0 | 0.0 |
| 89 | 0.0 | 0.0 |
| 90 | 0.0 | 0.0 |
| 91 | 0.0 | 0.0 |
| 92 | 0.0 | 0.0 |
| 93 | 0.0 | 0.0 |
| 94 | 0.0 | 0.0 |
| 95 | 0.0 | 0.0 |
| 96 | 0.0 | 0.0 |
| 97 | 0.0 | 0.0 |
| 98 | 0.0 | 0.0 |
| 99 | 0.0 | 0.0 |
| 100 | 0.0 | 0.0 |
| 101 | 0.0 | 0.0 |
| 102 | 0.0 | 0.0 |
| 103 | 0.0 | 0.0 |
| 104 | 0.0 | 0.0 |
| 105 | 0.0 | 0.0 |
| 106 | 0.0 | 0.0 |
| 107 | 0.0 | 0.0 |
| 108 | 0.0 | 0.0 |
| 109 | 0.0 | 0.0 |
| 110 | 0.0 | 0.0 |
| 111 | 0.0 | 0.0 |
| 112 | 0.0 | 0.0 |
| 113 | 0.0 | 0.0 |
| 114 | 0.0 | 0.0 |
| 115 | 0.0 | 0.0 |
| 116 | 0.0 | 0.0 |
| 117 | 0.0 | 0.0 |
| 118 | 0.0 | 0.0 |
| 119 | 0.0 | 0.0 |
| 120 | 0.0 | 0.0 |
| 121 | 0.0 | 0.0 |
| 122 | 0.0 | 0.0 |
| 123 | 0.0 | 0.0 |
| 124 | 0.0 | 0.0 |
| 125 | 0.0 | 0.0 |
| 126 | 0.0 | 0.0 |
| 127 | 0.0 | 0.0 |
| 128 | 0.0 | 0.0 |
| 129 | 0.0 | 0.0 |
| 130 | 0.0 | 0.0 |
| 131 | 0.0 | 0.0 |
| 132 | 0.0 | 0.0 |
| 133 | 0.0 | 0.0 |
| 134 | 0.0 | 0.0 |
| 135 | 0.0 | 0.0 |
| 136 | 0.0 | 0.0 |
| 137 | 0.0 | 0.0 |
| 138 | 0.0 | 0.0 |
| 139 | 0.0 | 0.0 |
| 140 | 0.0 | 0.0 |
| 141 | 0.0 | 0.0 |
| 142 | 0.0 | 0.0 |
| 143 | 0.0 | 0.0 |
| 144 | 0.0 | 0.0 |
| 145 | 0.0 | 0.0 |
| 146 | 0.0 | 0.0 |
| 147 | 0.0 | 0.0 |
| 148 | 0.0 | 0.0 |
| 149 | 0.0 | 0.0 |
| 150 | 0.0 | 0.0 |
| 151 | 0.0 | 0.0 |
| 152 | 0.0 | 0.0 |
| 153 | 0.0 | 0.0 |
| 154 | 0.0 | 0.0 |
| 155 | 0.0 | 0.0 |
| 156 | 0.0 | 0.0 |
| 157 | 0.0 | 0.0 |
| 158 | 0.0 | 0.032910667986686716 |
| 159 | 0.0 | 0.032910667986686716 |
| 160 | 0.0 | 0.032910667986686716 |
| 161 | 0.0 | 0.032910667986686716 |
| 162 | 0.0 | 0.032910667986686716 |
| 163 | 0.0 | 0.032910667986686716 |
| 164 | 0.0 | 0.032910667986686716 |
| 165 | 0.0 | 0.032910667986686716 |
| 166 | 0.0 | 0.032910667986686716 |
| 167 | 0.0 | 0.032910667986686716 |
| 168 | 0.0 | 0.032910667986686716 |
| 169 | 0.0 | 0.032910667986686716 |
| 170 | 0.0 | 0.032910667986686716 |
| 171 | 0.0 | 0.032910667986686716 |
| 172 | 0.0 | 0.032910667986686716 |
| 173 | 0.0 | 0.032910667986686716 |
| 174 | 0.0 | 0.032910667986686716 |
| 175 | 0.0 | 0.027889254651049425 |
| 176 | 0.0 | 0.022023673169799866 |
| 177 | 0.0 | 0.019641771500724877 |
| 178 | 0.0 | 0.019641771500724877 |
| 179 | 0.0 | 0.031036059851125203 |
| 180 | 0.0 | 0.031036059851125203 |
| 181 | 0.0 | 0.031036059851125203 |
| 182 | 0.0 | 0.031036059851125203 |
| 183 | 0.0 | 0.031036059851125203 |
| 184 | 0.0 | 0.031036059851125203 |
| 185 | 0.0 | 0.031036059851125203 |
| 186 | 0.0 | 0.031036059851125203 |
| 187 | 0.0 | 0.031036059851125203 |
| 188 | 0.0 | 0.031036059851125203 |
| 189 | 0.0 | 0.031036059851125203 |
| 190 | 0.0 | 0.031036059851125203 |
| 191 | 0.0 | 0.031036059851125203 |
| 192 | 0.0 | 0.031036059851125203 |
| 193 | 0.0 | 0.031036059851125203 |
| 194 | 0.0 | 0.031036059851125203 |
| 195 | 0.0 | 0.031036059851125203 |
| 196 | 0.0 | 0.012064074343867337 |
| 197 | 0.0 | 0.012064074343867337 |
| 198 | 0.0 | 0.012064074343867337 |
| 199 | 0.0 | 0.012064074343867337 |
| 200 | 0.0 | 0.012064074343867337 |
| 201 | 0.0 | 0.012064074343867337 |
| 202 | 0.0 | 0.012064074343867337 |
| 203 | 0.0 | 0.009630650675294885 |
| 204 | 0.0 | 0.018294273052329706 |
| 205 | 0.0 | 0.018294273052329706 |
| 206 | 0.0 | 0.018294273052329706 |
| 207 | 0.0 | 0.018294273052329706 |
| 208 | 0.0 | 0.018294273052329706 |
| 209 | 0.0 | 0.018294273052329706 |
| 210 | 0.0 | 0.018294273052329706 |
| 211 | 0.0 | 0.018294273052329706 |
| 212 | 0.0 | 0.018294273052329706 |
| 213 | 0.0 | 0.018294273052329706 |
| 214 | 0.0 | 0.018294273052329706 |
| 215 | 0.0 | 0.018294273052329706 |
| 216 | 0.0 | 0.018294273052329706 |
| 217 | 0.0 | 0.018294273052329706 |
| 218 | 0.0 | 0.0 |
| 219 | 0.0 | 0.0 |
| 220 | 0.0 | 0.0 |
| 221 | 0.0 | 0.0 |
| 222 | 0.0 | 0.0 |1.0
[PSI^+^]_weak_ - isolate #1
Δrnq1 strain
0.5
0
2 30 60 90 120 150 180 222
#### Chart
| Category | | |
|---|---|---|
| 1 | 1.0000000475315995 | 0.0 |
| 2 | 1.0000000475315995 | 0.0 |
| 3 | 1.0000000475315995 | 0.0 |
| 4 | 1.0000000475315995 | 0.0 |
| 5 | 1.0000000475315995 | 0.0 |
| 6 | 1.0000000475315995 | 0.0 |
| 7 | 1.0000000475315995 | 0.0 |
| 8 | 1.0000000475315995 | 0.0 |
| 9 | 1.0000000475315995 | 0.0 |
| 10 | 1.0000000475315995 | 0.0 |
| 11 | 1.0000000475315995 | 0.0 |
| 12 | 1.0000000475315995 | 0.0 |
| 13 | 1.0000000475315995 | 0.0 |
| 14 | 1.0000000475315995 | 0.0 |
| 15 | 1.0000000475315995 | 0.0 |
| 16 | 1.0000000475315995 | 0.0 |
| 17 | 1.0000000475315995 | 0.0 |
| 18 | 1.0000000475315995 | 0.0 |
| 19 | 1.0000000475315995 | 0.0 |
| 20 | 1.0000000475315995 | 0.0 |
| 21 | 1.0000000475315995 | 0.0 |
| 22 | 1.0000000475315995 | 0.0 |
| 23 | 1.0000000475315995 | 0.0 |
| 24 | 1.0000000475315995 | 0.0 |
| 25 | 1.0000000475315995 | 0.0 |
| 26 | 1.0000000475315995 | 0.0 |
| 27 | 1.0000000475315995 | 0.0 |
| 28 | 0.9970485684597487 | 0.0 |
| 29 | 0.9948603751183435 | 0.0 |
| 30 | 0.9927286616935675 | 0.0 |
| 31 | 0.9887169739009916 | 0.0 |
| 32 | 0.9735351723110657 | 0.0 |
| 33 | 0.9706869708010509 | 0.0 |
| 34 | 0.9672320136152484 | 0.0 |
| 35 | 0.9570962891481648 | 0.0 |
| 36 | 0.9570962891481648 | 0.0 |
| 37 | 0.9541415826525064 | 0.0 |
| 38 | 0.9350384611366113 | 0.0 |
| 39 | 0.9350384611366113 | 0.0 |
| 40 | 0.9350384611366113 | 0.0 |
| 41 | 0.9350384611366113 | 0.0 |
| 42 | 0.8220398987925462 | 0.0 |
| 43 | 0.8220398987925462 | 0.0 |
| 44 | 0.7183393903044769 | 0.0 |
| 45 | 0.5588672895052166 | 0.0 |
| 46 | 0.49145981662626154 | 0.0 |
| 47 | 0.4514011212342534 | 0.0 |
| 48 | 0.43601345522609847 | 0.0 |
| 49 | 0.4236403981956341 | 0.0 |
| 50 | 0.4201173353720955 | 0.0 |
| 51 | 0.4201173353720955 | 0.0 |
| 52 | 0.40691753075600673 | 0.0 |
| 53 | 0.40691753075600673 | 0.0 |
| 54 | 0.40067448469851935 | 0.0 |
| 55 | 0.39144131379717045 | 0.0 |
| 56 | 0.3759669331483713 | 0.0 |
| 57 | 0.33687617718040136 | 0.0 |
| 58 | 0.2937873661926111 | 0.0 |
| 59 | 0.28377009090538974 | 0.0 |
| 60 | 0.2750916896697153 | 0.0 |
| 61 | 0.2456264628477034 | 0.0 |
| 62 | 0.22079485835282195 | 0.0 |
| 63 | 0.22079485835282195 | 0.0 |
| 64 | 0.22079485835282195 | 0.0 |
| 65 | 0.22079485835282195 | 0.0 |
| 66 | 0.18832644848223967 | 0.0 |
| 67 | 0.18832644848223967 | 0.0 |
| 68 | 0.11835915941146694 | 0.0 |
| 69 | 0.0981407737988585 | 0.0 |
| 70 | 0.022484800267375753 | 0.0 |
| 71 | 0.00773719032439956 | 0.0 |
| 72 | 0.0 | 0.0 |
| 73 | 0.0 | 0.0 |
| 74 | 0.0 | 0.0 |
| 75 | 0.0 | 0.0 |
| 76 | 0.0 | 0.0 |
| 77 | 0.0 | 0.0 |
| 78 | 0.0 | 0.0 |
| 79 | 0.0 | 0.0 |
| 80 | 0.0 | 0.0022495143937453816 |
| 81 | 0.0 | 0.005664128781955731 |
| 82 | 0.0 | 0.005664128781955731 |
| 83 | 0.0 | 0.005664128781955731 |
| 84 | 0.0 | 0.005664128781955731 |
| 85 | 0.0 | 0.005664128781955731 |
| 86 | 0.0 | 0.005664128781955731 |
| 87 | 0.0 | 0.005664128781955731 |
| 88 | 0.0 | 0.005664128781955731 |
| 89 | 0.0 | 0.005664128781955731 |
| 90 | 0.0 | 0.012128658668142243 |
| 91 | 0.0 | 0.01804936764278486 |
| 92 | 0.0 | 0.01804936764278486 |
| 93 | 0.0 | 0.01804936764278486 |
| 94 | 0.0 | 0.01804936764278486 |
| 95 | 0.0 | 0.023126105291797032 |
| 96 | 0.0 | 0.023126105291797032 |
| 97 | 0.0 | 0.023126105291797032 |
| 98 | 0.0 | 0.023126105291797032 |
| 99 | 0.0 | 0.023126105291797032 |
| 100 | 0.0 | 0.031974087659735285 |
| 101 | 0.0 | 0.035196670331406395 |
| 102 | 0.0 | 0.035196670331406395 |
| 103 | 0.0 | 0.035196670331406395 |
| 104 | 0.0 | 0.03296652048050531 |
| 105 | 0.0 | 0.03296652048050531 |
| 106 | 0.0 | 0.03296652048050531 |
| 107 | 0.0 | 0.03296652048050531 |
| 108 | 0.0 | 0.03296652048050531 |
| 109 | 0.0 | 0.03296652048050531 |
| 110 | 0.0 | 0.03296652048050531 |
| 111 | 0.0 | 0.03296652048050531 |
| 112 | 0.0 | 0.03296652048050531 |
| 113 | 0.0 | 0.027539607348392166 |
| 114 | 0.0 | 0.027539607348392166 |
| 115 | 0.0 | 0.027539607348392166 |
| 116 | 0.0 | 0.027539607348392166 |
| 117 | 0.0 | 0.027539607348392166 |
| 118 | 0.0 | 0.027539607348392166 |
| 119 | 0.0 | 0.027539607348392166 |
| 120 | 0.0 | 0.0 |
| 121 | 0.0 | 0.0 |
| 122 | 0.0 | 0.0 |
| 123 | 0.0 | 0.0 |
| 124 | 0.0 | 0.0 |
| 125 | 0.0 | 0.0 |
| 126 | 0.0 | 0.0 |
| 127 | 0.0 | 0.0 |
| 128 | 0.0 | 0.0 |
| 129 | 0.0 | 0.0 |
| 130 | 0.0 | 0.0 |
| 131 | 0.0 | 0.0 |
| 132 | 0.0 | 0.0 |
| 133 | 0.0 | 0.0 |
| 134 | 0.0 | 0.0 |
| 135 | 0.0 | 0.0 |
| 136 | 0.0 | 0.0 |
| 137 | 0.0 | 0.0 |
| 138 | 0.0 | 0.0 |
| 139 | 0.0 | 0.0 |
| 140 | 0.0 | 0.0 |
| 141 | 0.0 | 0.0 |
| 142 | 0.0 | 0.0 |
| 143 | 0.0 | 0.0 |
| 144 | 0.0 | 0.0 |
| 145 | 0.0 | 0.0 |
| 146 | 0.0 | 0.0 |
| 147 | 0.0 | 0.0 |
| 148 | 0.0 | 0.0 |
| 149 | 0.0 | 0.0 |
| 150 | 0.0 | 0.0 |
| 151 | 0.0 | 0.0 |
| 152 | 0.0 | 0.0 |
| 153 | 0.0 | 0.0 |
| 154 | 0.0 | 0.0 |
| 155 | 0.0 | 0.028995175485521534 |
| 156 | 0.0 | 0.03422521576538436 |
| 157 | 0.0 | 0.04804827193240225 |
| 158 | 0.0 | 0.09891085742484176 |
| 159 | 0.0 | 0.10251588981768767 |
| 160 | 0.0 | 0.10251588981768767 |
| 161 | 0.0 | 0.10251588981768767 |
| 162 | 0.0 | 0.10251588981768767 |
| 163 | 0.0 | 0.10251588981768767 |
| 164 | 0.0 | 0.10251588981768767 |
| 165 | 0.0 | 0.10251588981768767 |
| 166 | 0.0 | 0.105951482460646 |
| 167 | 0.0 | 0.105951482460646 |
| 168 | 0.0 | 0.105951482460646 |
| 169 | 0.0 | 0.105951482460646 |
| 170 | 0.0 | 0.105951482460646 |
| 171 | 0.0 | 0.11108631373819106 |
| 172 | 0.0 | 0.11108631373819106 |
| 173 | 0.0 | 0.11108631373819106 |
| 174 | 0.0 | 0.11108631373819106 |
| 175 | 0.0 | 0.09996461129795205 |
| 176 | 0.0 | 0.09071158724218732 |
| 177 | 0.0 | 0.085625167321753 |
| 178 | 0.0 | 0.07930587150689869 |
| 179 | 0.0 | 0.08982565940706091 |
| 180 | 0.0 | 0.09353396936174305 |
| 181 | 0.0 | 0.09353396936174305 |
| 182 | 0.0 | 0.09353396936174305 |
| 183 | 0.0 | 0.09353396936174305 |
| 184 | 0.0 | 0.09353396936174305 |
| 185 | 0.0 | 0.09353396936174305 |
| 186 | 0.0 | 0.09353396936174305 |
| 187 | 0.0 | 0.09353396936174305 |
| 188 | 0.0 | 0.09353396936174305 |
| 189 | 0.0 | 0.09353396936174305 |
| 190 | 0.0 | 0.09353396936174305 |
| 191 | 0.0 | 0.09353396936174305 |
| 192 | 0.0 | 0.09353396936174305 |
| 193 | 0.0 | 0.09353396936174305 |
| 194 | 0.0 | 0.09353396936174305 |
| 195 | 0.0 | 0.09353396936174305 |
| 196 | 0.0 | 0.04799824686338783 |
| 197 | 0.0 | 0.04799824686338783 |
| 198 | 0.0 | 0.04799824686338783 |
| 199 | 0.0 | 0.04799824686338783 |
| 200 | 0.0 | 0.04799824686338783 |
| 201 | 0.0 | 0.04799824686338783 |
| 202 | 0.0 | 0.050447861533190905 |
| 203 | 0.0 | 0.09115535801570238 |
| 204 | 0.0 | 0.10132012929705256 |
| 205 | 0.0 | 0.10132012929705256 |
| 206 | 0.0 | 0.10132012929705256 |
| 207 | 0.0 | 0.10132012929705256 |
| 208 | 0.0 | 0.10132012929705256 |
| 209 | 0.0 | 0.10132012929705256 |
| 210 | 0.0 | 0.10132012929705256 |
| 211 | 0.0 | 0.10132012929705256 |
| 212 | 0.0 | 0.10132012929705256 |
| 213 | 0.0 | 0.10132012929705256 |
| 214 | 0.0 | 0.10132012929705256 |
| 215 | 0.0 | 0.10132012929705256 |
| 216 | 0.0 | 0.09455706270868319 |
| 217 | 0.0 | 0.09455706270868319 |
| 218 | 0.0 | 0.0 |
| 219 | 0.0 | 0.0 |
| 220 | 0.0 | 0.0 |
| 221 | 0.0 | 0.0 |
| 222 | 0.0 | 0.0 |1.0
PrK resistance index
[PSI^+^]_weak_ - isolate #2
Δrnq1 strain
0.5
0
2 30 60 90 120 150 180 222
#### Chart
| Category | | |
|---|---|---|
| 1 | 0.0 | 1.0 |
| 2 | 0.0 | 1.0 |
| 3 | 0.0 | 1.0 |
| 4 | 0.0 | 1.0 |
| 5 | 0.0 | 1.0 |
| 6 | 0.0 | 1.0 |
| 7 | 0.0 | 1.0 |
| 8 | 0.0 | 1.0 |
| 9 | 0.0 | 1.0 |
| 10 | 0.0 | 1.0 |
| 11 | 0.0 | 1.0 |
| 12 | 0.0 | 1.0 |
| 13 | 0.0 | 1.0 |
| 14 | 0.0 | 1.0 |
| 15 | 0.0 | 1.0 |
| 16 | 0.0 | 1.0 |
| 17 | 0.0 | 1.0 |
| 18 | 0.0 | 1.0 |
| 19 | 0.0 | 1.0 |
| 20 | 0.0 | 1.0 |
| 21 | 0.0 | 1.0 |
| 22 | 0.0 | 1.0 |
| 23 | 0.0 | 1.0 |
| 24 | 0.0 | 1.0 |
| 25 | 0.0 | 1.0 |
| 26 | 0.0 | 1.0 |
| 27 | 0.0 | 1.0 |
| 28 | 0.0 | 1.0 |
| 29 | 0.0 | 1.0 |
| 30 | 0.0 | 1.0 |
| 31 | 0.0 | 1.0 |
| 32 | 0.0 | 0.9950747176699571 |
| 33 | 0.0 | 0.9950747176699571 |
| 34 | 0.0 | 0.9950747176699571 |
| 35 | 0.0 | 0.9885862663684879 |
| 36 | 0.0 | 0.9885862663684879 |
| 37 | 0.0 | 0.9831248704061529 |
| 38 | 0.0 | 0.9614035644395145 |
| 39 | 0.0 | 0.9614035644395145 |
| 40 | 0.0 | 0.9614035644395145 |
| 41 | 0.0 | 0.9614035644395145 |
| 42 | 0.0 | 0.516417350690811 |
| 43 | 0.0 | 0.516417350690811 |
| 44 | 0.0 | 0.4800698159651687 |
| 45 | 0.0 | 0.3439674656368349 |
| 46 | 0.0 | 0.3268384918570443 |
| 47 | 0.0 | 0.32018323495769374 |
| 48 | 0.0 | 0.32018323495769374 |
| 49 | 0.0 | 0.31754563521312446 |
| 50 | 0.0 | 0.31754563521312446 |
| 51 | 0.0 | 0.31754563521312446 |
| 52 | 0.0 | 0.3121880245038966 |
| 53 | 0.0 | 0.3121880245038966 |
| 54 | 0.0 | 0.3121880245038966 |
| 55 | 0.0 | 0.3064033738360242 |
| 56 | 0.0 | 0.2992725997897412 |
| 57 | 0.0 | 0.27068229659357407 |
| 58 | 0.0 | 0.2407908215935906 |
| 59 | 0.0 | 0.23424309591189668 |
| 60 | 0.0 | 0.22395513480294132 |
| 61 | 0.0 | 0.1959384048845615 |
| 62 | 0.0 | 0.17234654050258064 |
| 63 | 0.0 | 0.17234654050258064 |
| 64 | 0.0 | 0.17234654050258064 |
| 65 | 0.0 | 0.17234654050258064 |
| 66 | 0.0 | 0.15389127492140636 |
| 67 | 0.0 | 0.15389127492140636 |
| 68 | 0.0 | 0.12942880430794737 |
| 69 | 0.0 | 0.11463290538993617 |
| 70 | 0.0 | 0.021762731927538942 |
| 71 | 0.0 | 0.0 |
| 72 | 0.0 | 0.0 |
| 73 | 0.0 | 0.0 |
| 74 | 0.0 | 0.0 |
| 75 | 0.0 | 0.0 |
| 76 | 0.0 | 0.0 |
| 77 | 0.0 | 0.0 |
| 78 | 0.0 | 0.0 |
| 79 | 0.0 | 0.0 |
| 80 | 0.0 | 0.0 |
| 81 | 0.0 | 0.0 |
| 82 | 0.0 | 0.0 |
| 83 | 0.0 | 0.0 |
| 84 | 0.0 | 0.0 |
| 85 | 0.0 | 0.0 |
| 86 | 0.0 | 0.0 |
| 87 | 0.0 | 0.0 |
| 88 | 0.0 | 0.0 |
| 89 | 0.0 | 0.0 |
| 90 | 0.00817589815559723 | 0.0 |
| 91 | 0.014894395750577815 | 0.0 |
| 92 | 0.014894395750577815 | 0.0 |
| 93 | 0.014894395750577815 | 0.0 |
| 94 | 0.014894395750577815 | 0.0 |
| 95 | 0.01935209356449901 | 0.0 |
| 96 | 0.01935209356449901 | 0.0 |
| 97 | 0.01935209356449901 | 0.0 |
| 98 | 0.01935209356449901 | 0.0 |
| 99 | 0.01935209356449901 | 0.0 |
| 100 | 0.01935209356449901 | 0.0 |
| 101 | 0.01935209356449901 | 0.0 |
| 102 | 0.01935209356449901 | 0.0 |
| 103 | 0.01935209356449901 | 0.0 |
| 104 | 0.01935209356449901 | 0.0 |
| 105 | 0.01935209356449901 | 0.0 |
| 106 | 0.01935209356449901 | 0.0 |
| 107 | 0.01935209356449901 | 0.0 |
| 108 | 0.01935209356449901 | 0.0 |
| 109 | 0.01935209356449901 | 0.0 |
| 110 | 0.01935209356449901 | 0.0 |
| 111 | 0.01935209356449901 | 0.0 |
| 112 | 0.01935209356449901 | 0.0 |
| 113 | 0.01935209356449901 | 0.0 |
| 114 | 0.01935209356449901 | 0.0 |
| 115 | 0.01935209356449901 | 0.0 |
| 116 | 0.01935209356449901 | 0.0 |
| 117 | 0.01935209356449901 | 0.0 |
| 118 | 0.01935209356449901 | 0.0 |
| 119 | 0.01935209356449901 | 0.0 |
| 120 | 0.004691049370270307 | 0.0 |
| 121 | 0.0 | 0.0 |
| 122 | 0.0 | 0.0 |
| 123 | 0.0 | 0.0 |
| 124 | 0.0 | 0.0 |
| 125 | 0.0 | 0.0 |
| 126 | 0.0 | 0.0 |
| 127 | 0.0 | 0.0 |
| 128 | 0.0 | 0.0 |
| 129 | 0.0 | 0.0 |
| 130 | 0.0 | 0.0 |
| 131 | 0.0 | 0.0 |
| 132 | 0.0 | 0.0 |
| 133 | 0.0 | 0.0 |
| 134 | 0.0 | 0.0 |
| 135 | 0.0 | 0.0 |
| 136 | 0.0 | 0.0 |
| 137 | 0.0 | 0.0 |
| 138 | 0.0 | 0.0 |
| 139 | 0.0 | 0.0 |
| 140 | 0.0 | 0.0 |
| 141 | 0.0 | 0.0 |
| 142 | 0.0 | 0.0 |
| 143 | 0.0 | 0.0 |
| 144 | 0.0 | 0.0 |
| 145 | 0.0 | 0.0 |
| 146 | 0.0 | 0.0 |
| 147 | 0.0 | 0.0 |
| 148 | 0.0 | 0.0 |
| 149 | 0.0 | 0.0 |
| 150 | 0.0 | 0.0 |
| 151 | 0.0 | 0.0 |
| 152 | 0.0 | 0.0 |
| 153 | 0.0 | 0.0 |
| 154 | 0.0 | 0.0 |
| 155 | 0.0 | 0.0 |
| 156 | 0.0 | 0.0 |
| 157 | 0.0 | 0.0 |
| 158 | 0.0 | 0.0 |
| 159 | 0.0 | 0.0 |
| 160 | 0.0 | 0.0 |
| 161 | 0.0 | 0.0 |
| 162 | 0.0 | 0.0 |
| 163 | 0.0 | 0.0 |
| 164 | 0.0 | 0.0 |
| 165 | 0.0 | 0.0 |
| 166 | 0.0 | 0.0 |
| 167 | 0.0 | 0.0 |
| 168 | 0.0 | 0.0 |
| 169 | 0.0 | 0.0 |
| 170 | 0.0 | 0.0 |
| 171 | 0.0 | 0.0 |
| 172 | 0.0 | 0.0 |
| 173 | 0.0 | 0.0 |
| 174 | 0.0 | 0.0 |
| 175 | 0.0 | 0.0 |
| 176 | 0.0 | 0.0 |
| 177 | 0.0 | 0.0 |
| 178 | 0.0 | 0.0 |
| 179 | 0.0 | 0.0 |
| 180 | 0.0 | 0.0 |
| 181 | 0.0 | 0.0 |
| 182 | 0.0 | 0.0 |
| 183 | 0.0 | 0.0 |
| 184 | 0.0 | 0.0 |
| 185 | 0.0 | 0.0 |
| 186 | 0.0 | 0.0 |
| 187 | 0.0 | 0.0 |
| 188 | 0.0 | 0.0 |
| 189 | 0.0 | 0.0 |
| 190 | 0.0 | 0.0 |
| 191 | 0.0 | 0.0 |
| 192 | 0.0 | 0.0 |
| 193 | 0.0 | 0.0 |
| 194 | 0.0 | 0.0 |
| 195 | 0.0 | 0.0 |
| 196 | 0.0 | 0.0 |
| 197 | 0.0 | 0.0 |
| 198 | 0.0 | 0.0 |
| 199 | 0.0 | 0.0 |
| 200 | 0.0 | 0.0 |
| 201 | 0.0 | 0.0 |
| 202 | 0.0 | 0.0 |
| 203 | 0.0 | 0.0 |
| 204 | 0.0 | 0.0 |
| 205 | 0.0 | 0.0 |
| 206 | 0.0 | 0.0 |
| 207 | 0.0 | 0.0 |
| 208 | 0.0 | 0.0 |
| 209 | 0.0 | 0.0 |
| 210 | 0.0 | 0.0 |
| 211 | 0.0 | 0.0 |
| 212 | 0.0 | 0.0 |
| 213 | 0.0 | 0.0 |
| 214 | 0.0 | 0.0 |
| 215 | 0.0 | 0.0 |
| 216 | 0.0 | 0.0 |
| 217 | 0.0 | 0.0 |
| 218 | 0.0 | 0.0 |
| 219 | 0.0 | 0.0 |
| 220 | 0.0 | 0.0 |
| 220 | 0.0 | 0.0 |
| 220 | 0.0 | 0.0 |1.0
[PSI^+^]_weak_
[RNQ^+^] strain
0.5
0
2 30 60 90 120 150 180 222
Sup35PD amino acid position
Intensity (a.u.)
4000
8000
m/z
7000
5000
6000
4000
8000
m/z
7000
5000
6000
2-42
2-45
2-44
2-38
2-46
2-70
2-45
2-42
2-44
2-46
2-70
2-68
2-47
2-58
2-57
2-66
2-61
2-38
2-62
2-71
2-69
2-48
2-49
2-52
2-56
2-60
2-72
2-59
2-55
2-50
2-42
fragment of Rnq1
2-45
2-70
2-44
2-61
2-57
2-58
2-38
2-46
2-62
2-68
2-67
2-71
PrK-resistant cores of Sup35PD-GFP prion polymers extracted from weak [PSI+] isolates, emerged in either Δrnq1 or [RNQ+] cells. Sup35PD-GFP fibrils, sonicated and isolated from yeast cell lysates in the presence of sarcosyl, were treated for 1 hr with 25 μg/ml proteinase K, and proteinase K-resistant peptides were analyzed with MALDI-TOF. The left panels show PrK resistance index profiles, and the right panels show a part of the MALDI-TOF mass spectra (linear mode) containing the Core 1 peptides. Unlabelled peaks represent parts of Cores 2-4. Note that the N-terminal methionine of Sup35PD-GFP has been completely removed and replaced with an acetyl group in the yeast cells, and therefore all Core 1 peaks start from the amino acid residue №2.

### Slide 2
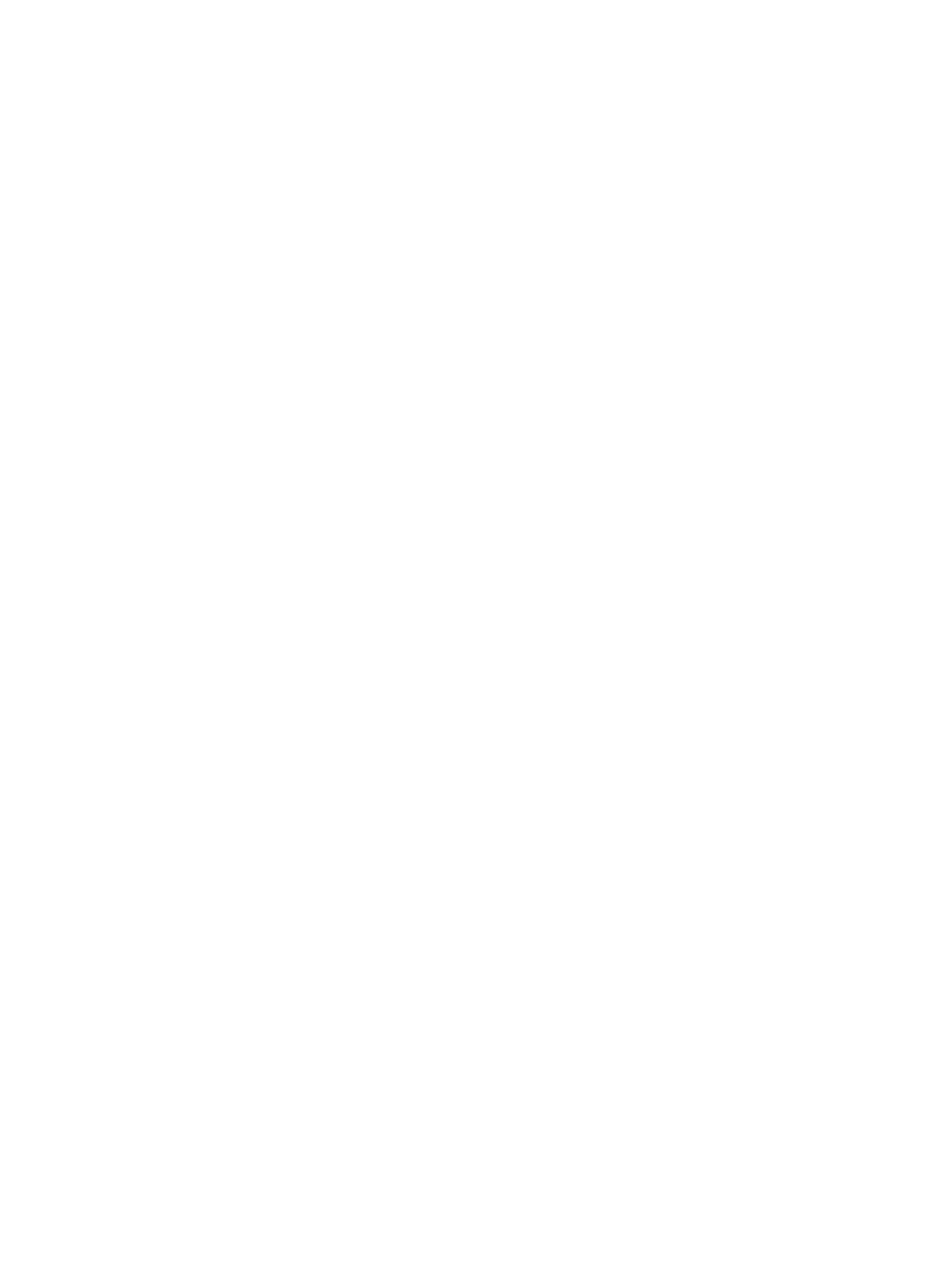
