## Supplementary material for "A liquid-to-solid phase transition of biomolecular condensates drives *in vivo* formation of yeast amyloids and prions": Figure S6

### Slide 1
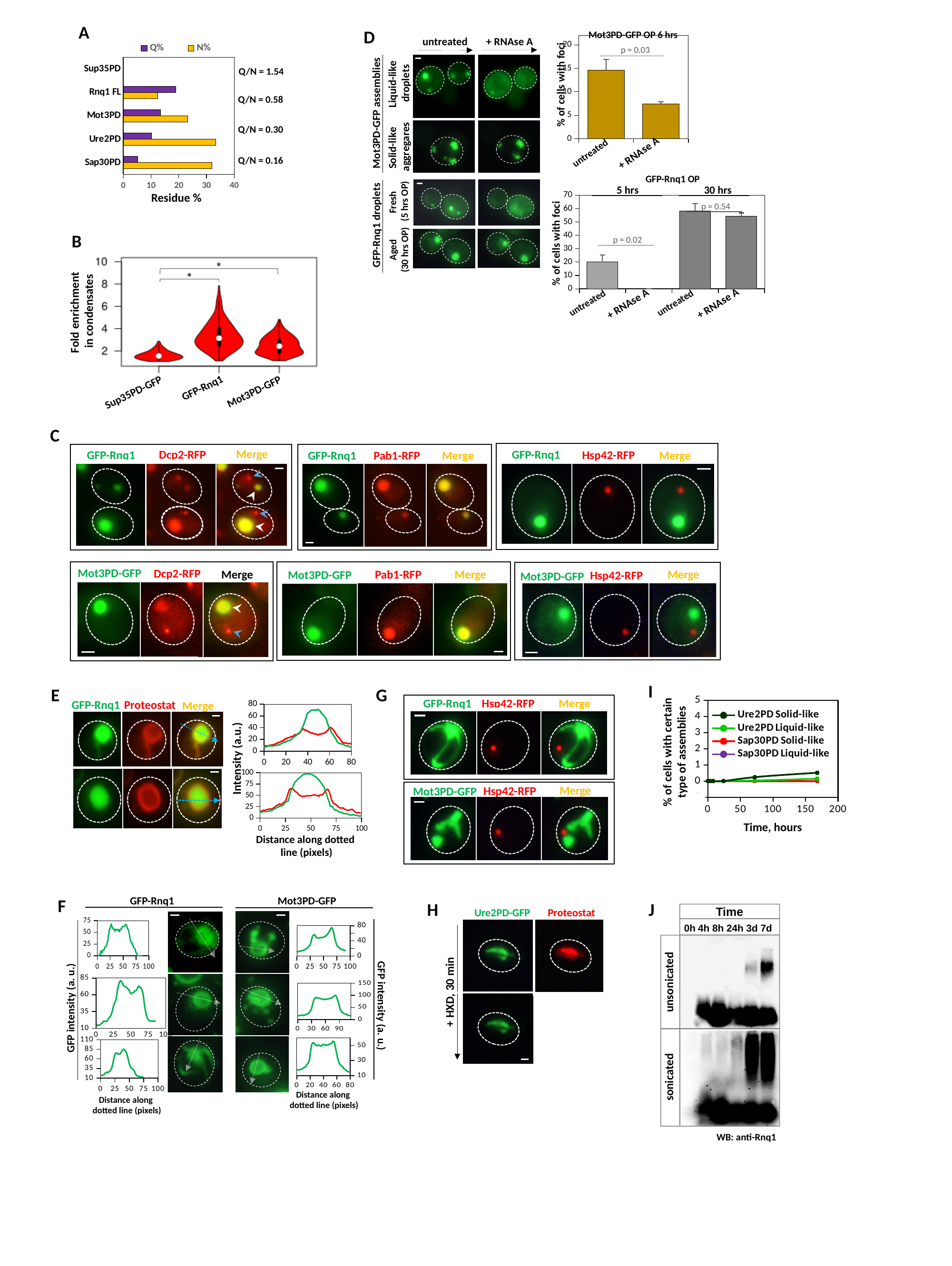

A
[unsupported chart]
Residue %
Q/N = 1.54
Q/N = 0.58
Q/N = 0.30
Q/N = 0.16
D
+ RNAse A
untreated
Liquid-like
 droplets
Mot3PD-GFP assemblies
Solid-like
aggregares
Mot3PD-GFP OP 6 hrs
#### Chart
| Category | |
|---|---|
| control | 14.619924749343332 |
| + RNAse A | 7.418256191365434 |p = 0.03
% of cells with foci
+ RNAse A
untreated
GFP-Rnq1 OP
5 hrs
30 hrs
#### Chart
| Category | |
|---|---|
| contr | 20.0598544442643 |
| RNAse | 0.0 |
| contr | 58.31392454199471 |
| RNAse | 54.483131432688715 |p = 0.54
p = 0.02
% of cells with foci
+ RNAse A
untreated
untreated
+ RNAse A
Fresh
(5 hrs OP)
GFP-Rnq1 droplets
Aged
(30 hrs OP)
B
*
*
Fold enrichment
in condensates
GFP-Rnq1
Mot3PD-GFP
Sup35PD-GFP
C
Merge
Dcp2-RFP
GFP-Rnq1
GFP-Rnq1
Hsp42-RFP
Merge
Merge
GFP-Rnq1
Pab1-RFP
Mot3PD-GFP
Dcp2-RFP
Merge
Merge
Pab1-RFP
Mot3PD-GFP
Merge
Hsp42-RFP
Mot3PD-GFP
I
#### Chart
| Category | Ure2PD Solid-like | Ure2PD Liquid-like | Sap30PD Solid-like | Sap30PD Liquid-like |
|---|---|---|---|---|% of cells with certain type of assemblies
Time, hours
G
Merge
GFP-Rnq1
Hsp42-RFP
Merge
Hsp42-RFP
Mot3PD-GFP
E
#### Chart
| Category | GFP-Rnq1 | Proteostat |
|---|---|---|GFP-Rnq1
Proteostat
Merge
Intensity (a.u.)
#### Chart
| Category | GFP-Rnq1 | Proteostat |
|---|---|---|
Distance along dotted
line (pixels)
Mot3PD-GFP
#### Chart
| Category |
|---|
#### Chart
| Category |
|---|
GFP intensity (a. u.)
#### Chart
| Category |
|---|
Distance along
dotted line (pixels)
GFP-Rnq1
#### Chart
| Category |
|---|
#### Chart
| Category |
|---|
#### Chart
| Category | |
|---|---|GFP intensity (a. u.)
Distance along
dotted line (pixels)
F
J
Time
0h 4h 8h 24h 3d 7d
unsonicated
sonicated
WB: anti-Rnq1
H
Ure2PD-GFP Proteostat
+ HXD, 30 min

### Slide 2
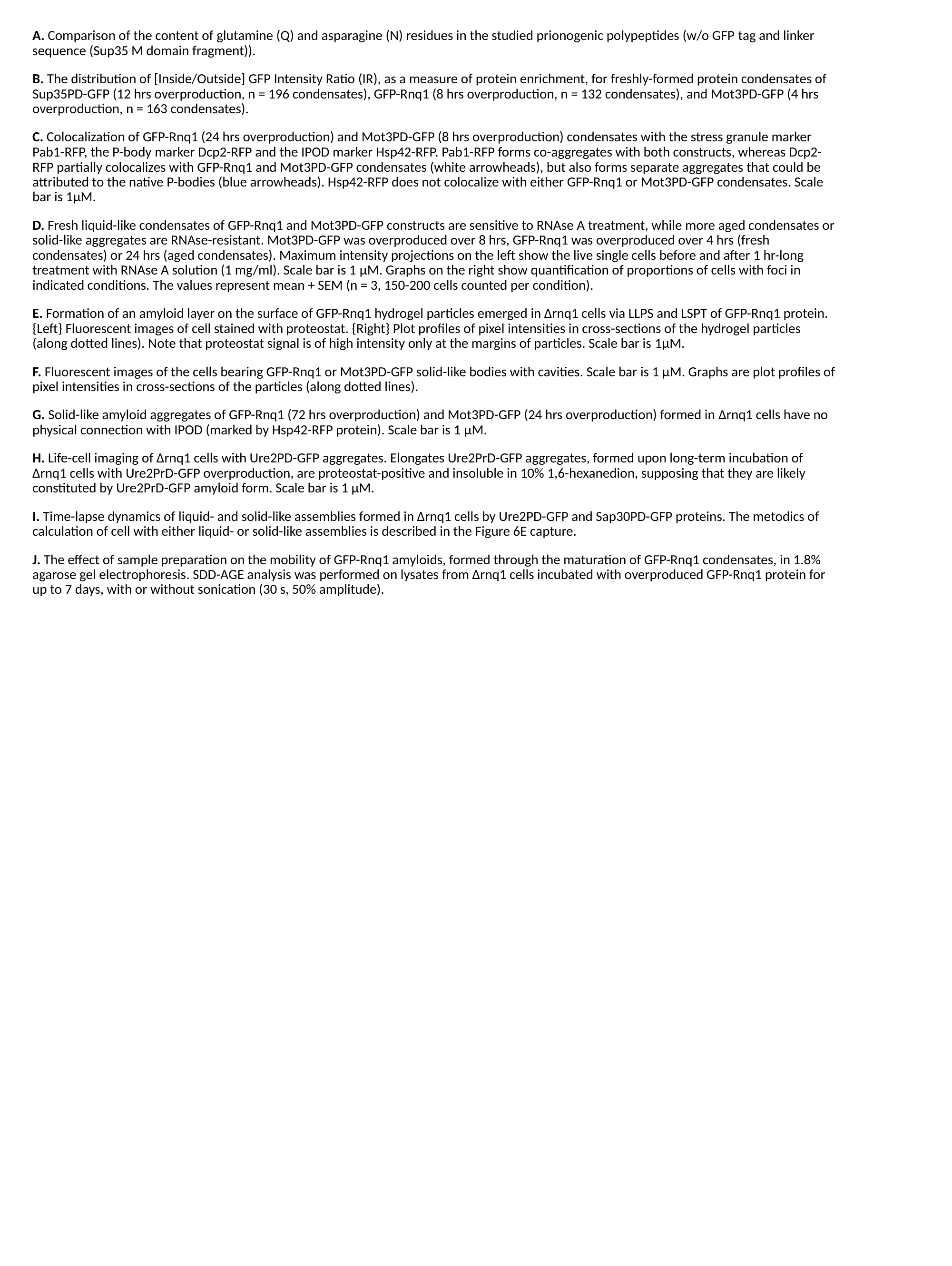

A. Comparison of the content of glutamine (Q) and asparagine (N) residues in the studied prionogenic polypeptides (w/o GFP tag and linker sequence (Sup35 M domain fragment)).
B. The distribution of [Inside/Outside] GFP Intensity Ratio (IR), as a measure of protein enrichment, for freshly-formed protein condensates of Sup35PD-GFP (12 hrs overproduction, n = 196 condensates), GFP-Rnq1 (8 hrs overproduction, n = 132 condensates), and Mot3PD-GFP (4 hrs overproduction, n = 163 condensates).
C. Colocalization of GFP-Rnq1 (24 hrs overproduction) and Mot3PD-GFP (8 hrs overproduction) condensates with the stress granule marker Pab1-RFP, the P-body marker Dcp2-RFP and the IPOD marker Hsp42-RFP. Pab1-RFP forms co-aggregates with both constructs, whereas Dcp2-RFP partially colocalizes with GFP-Rnq1 and Mot3PD-GFP condensates (white arrowheads), but also forms separate aggregates that could be attributed to the native P-bodies (blue arrowheads). Hsp42-RFP does not colocalize with either GFP-Rnq1 or Mot3PD-GFP condensates. Scale bar is 1μM.
D. Fresh liquid-like condensates of GFP-Rnq1 and Mot3PD-GFP constructs are sensitive to RNAse A treatment, while more aged condensates or solid-like aggregates are RNAse-resistant. Mot3PD-GFP was overproduced over 8 hrs, GFP-Rnq1 was overproduced over 4 hrs (fresh condensates) or 24 hrs (aged condensates). Maximum intensity projections on the left show the live single cells before and after 1 hr-long treatment with RNAse A solution (1 mg/ml). Scale bar is 1 μM. Graphs on the right show quantification of proportions of cells with foci in indicated conditions. The values represent mean + SEM (n = 3, 150-200 cells counted per condition).
E. Formation of an amyloid layer on the surface of GFP-Rnq1 hydrogel particles emerged in Δrnq1 cells via LLPS and LSPT of GFP-Rnq1 protein. {Left} Fluorescent images of cell stained with proteostat. {Right} Plot profiles of pixel intensities in cross-sections of the hydrogel particles (along dotted lines). Note that proteostat signal is of high intensity only at the margins of particles. Scale bar is 1μM.
F. Fluorescent images of the cells bearing GFP-Rnq1 or Mot3PD-GFP solid-like bodies with cavities. Scale bar is 1 μM. Graphs are plot profiles of pixel intensities in cross-sections of the particles (along dotted lines).
G. Solid-like amyloid aggregates of GFP-Rnq1 (72 hrs overproduction) and Mot3PD-GFP (24 hrs overproduction) formed in Δrnq1 cells have no physical connection with IPOD (marked by Hsp42-RFP protein). Scale bar is 1 μM.
H. Life-cell imaging of Δrnq1 cells with Ure2PD-GFP aggregates. Elongates Ure2PrD-GFP aggregates, formed upon long-term incubation of Δrnq1 cells with Ure2PrD-GFP overproduction, are proteostat-positive and insoluble in 10% 1,6-hexanedion, supposing that they are likely constituted by Ure2PrD-GFP amyloid form. Scale bar is 1 μM.
I. Time-lapse dynamics of liquid- and solid-like assemblies formed in Δrnq1 cells by Ure2PD-GFP and Sap30PD-GFP proteins. The metodics of calculation of cell with either liquid- or solid-like assemblies is described in the Figure 6E capture.
J. The effect of sample preparation on the mobility of GFP-Rnq1 amyloids, formed through the maturation of GFP-Rnq1 condensates, in 1.8% agarose gel electrophoresis. SDD-AGE analysis was performed on lysates from ∆rnq1 cells incubated with overproduced GFP-Rnq1 protein for up to 7 days, with or without sonication (30 s, 50% amplitude).
