## Supplementary material for "A liquid-to-solid phase transition of biomolecular condensates drives *in vivo* formation of yeast amyloids and prions": Figure S7

### Slide 1
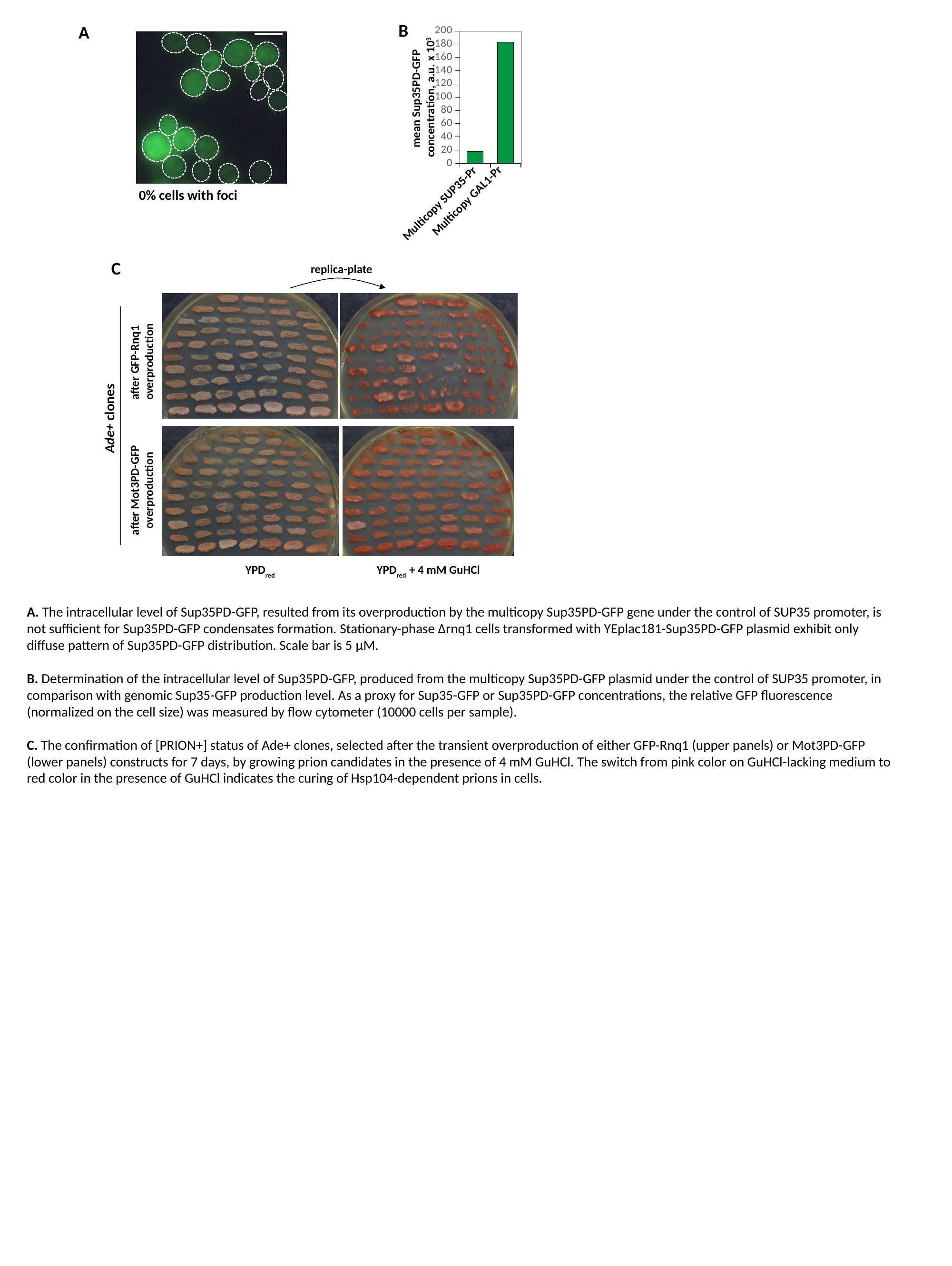

B
#### Chart
| Category | |
|---|---|
| Multicopy NMG | 17.818338297215075 |
| Gal-NMG | 183.34054260712338 |Multicopy GAL1-Pr
Multicopy SUP35-Pr
mean Sup35PD-GFP
concentration, a.u. x 10^3^
A
0% cells with foci
C
replica-plate
after GFP-Rnq1
overproduction
after Mot3PD-GFP
overproduction
YPD_red_
YPD_red_ + 4 mM GuHCl
Ade+ clones
A. The intracellular level of Sup35PD-GFP, resulted from its overproduction by the multicopy Sup35PD-GFP gene under the control of SUP35 promoter, is not sufficient for Sup35PD-GFP condensates formation. Stationary-phase Δrnq1 cells transformed with YEplac181-Sup35PD-GFP plasmid exhibit only diffuse pattern of Sup35PD-GFP distribution. Scale bar is 5 μM.
B. Determination of the intracellular level of Sup35PD-GFP, produced from the multicopy Sup35PD-GFP plasmid under the control of SUP35 promoter, in comparison with genomic Sup35-GFP production level. As a proxy for Sup35-GFP or Sup35PD-GFP concentrations, the relative GFP fluorescence (normalized on the cell size) was measured by flow cytometer (10000 cells per sample).
C. The confirmation of [PRION+] status of Ade+ clones, selected after the transient overproduction of either GFP-Rnq1 (upper panels) or Mot3PD-GFP (lower panels) constructs for 7 days, by growing prion candidates in the presence of 4 mM GuHCl. The switch from pink color on GuHCl-lacking medium to red color in the presence of GuHCl indicates the curing of Hsp104-dependent prions in cells.
