## Supplementary material for "A liquid-to-solid phase transition of biomolecular condensates drives *in vivo* formation of yeast amyloids and prions": Table 1

### Slide 1
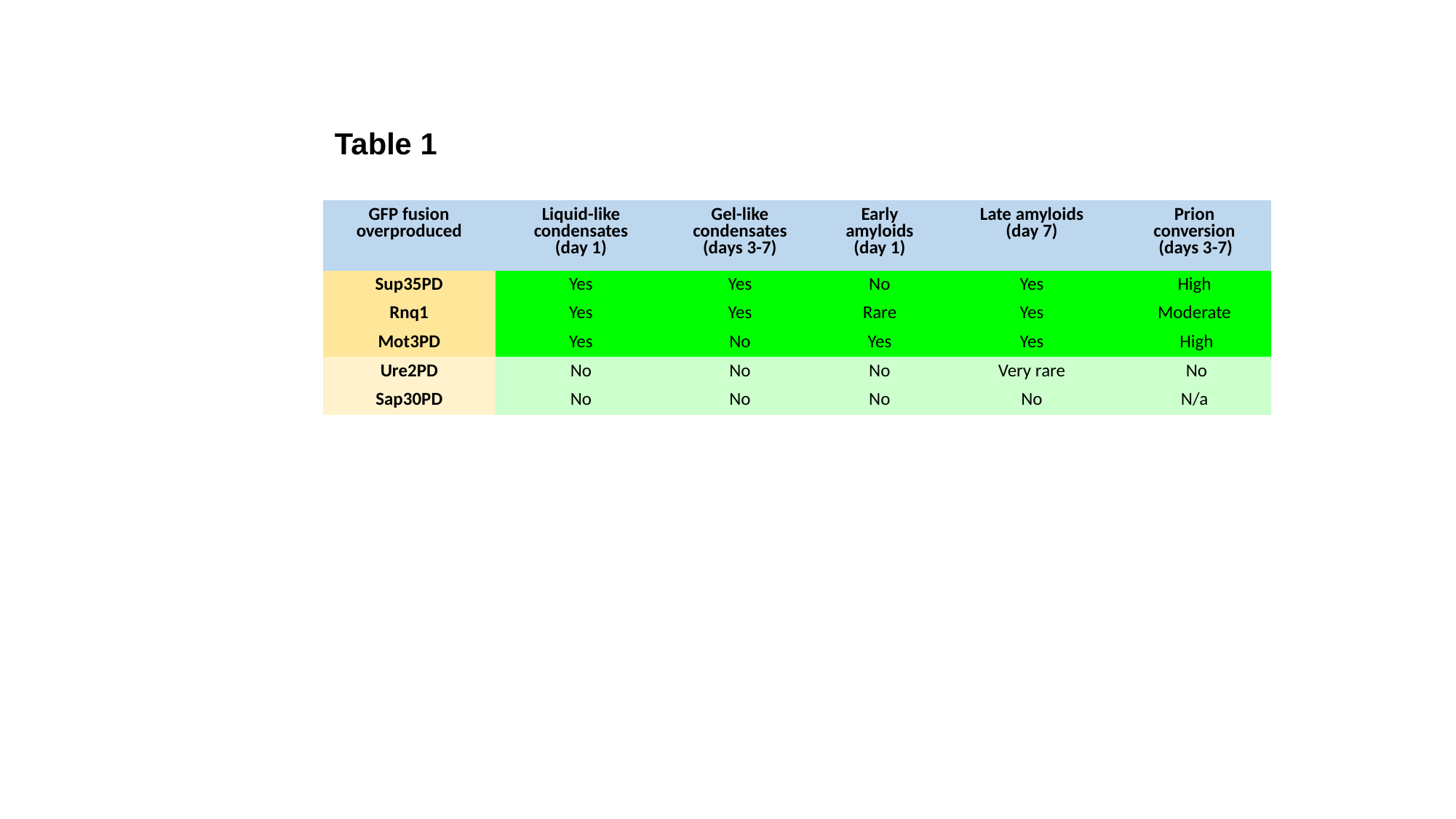

Table 1
| GFP fusion overproduced | Liquid-like condensates (day 1) | Gel-like condensates (days 3-7) | Early amyloids (day 1) | Late amyloids (day 7) | Prion conversion (days 3-7) |
| --- | --- | --- | --- | --- | --- |
| Sup35PD | Yes | Yes | No | Yes | High |
| Rnq1 | Yes | Yes | Rare | Yes | Moderate |
| Mot3PD | Yes | No | Yes | Yes | High |
| Ure2PD | No | No | No | Very rare | No |
| Sap30PD | No | No | No | No | N/a |
